# The Predicted Metabolic Function of the Gut Microbiota of *Drosophila melanogaster*

**DOI:** 10.1101/2021.01.20.427455

**Authors:** Nana Y.D. Ankrah, Brandon E. Barker, Joan Song, Cindy Wu, John G. McMullen, Angela E. Douglas

## Abstract

An important goal for many nutrition-based microbiome studies is to identify the metabolic function of microbes in complex microbial communities and its impact on host physiology. This research can be confounded by poorly-understood effects of community composition and host diet on the metabolic traits of individual taxa. Here, we investigated these multi-way interactions by constructing and analyzing metabolic models comprising every combination of five bacterial members of the *Drosophila* gut microbiome (from single taxa to the five-member community of *Acetobacter* and *Lactobacillus* species) under three nutrient regimes. We show that the metabolic function of *Drosophila* gut bacteria is dynamic, influenced by community composition and responsive to dietary modulation. Furthermore, we show that ecological interactions such as competition and mutualism identified from the growth patterns of gut bacteria are underlain by a diversity of metabolic interactions, and show that the bacteria tend to compete for amino acids and B vitamins more frequently than for carbon sources. Our results reveal that in addition to fermentation products such as acetate, intermediates of the tricarboxylic acid (TCA) cycle including 2-oxoglutarate and succinate are produced at high flux and cross-fed between bacterial taxa suggesting important roles for TCA cycle intermediates in modulating *Drosophila* gut microbe interactions and the potential to influence host traits. These metabolic models provide specific predictions of the patterns of ecological and metabolic interactions among gut bacteria under different nutrient regimes, with potentially important consequences for overall community metabolic function and nutritional interactions with the host.

**IMPORTANCE:** *Drosophila* is an important model for microbiome research partly because of the low complexity of its mostly culturable gut microbiota. Our current understanding of how *Drosophila* interacts with its gut microbes and how these interactions influence host traits derives almost entirely from empirical studies that focus on individual microbial taxa or classes of metabolites. These studies have failed to capture fully the complexity of metabolic interactions that occur between host and microbe. To overcome this limitation, we reconstructed and analyzed 31 metabolic models for every combination of the five principal bacterial taxa in the gut microbiome of *Drosophila*. This revealed that metabolic interactions between between *Drosophila* gut bacterial taxa are highly dynamic and influenced by co-occurring bacteria and nutrient availability. Our results generate testable hypothesis about among-microbe ecological interactions in the *Drosophila* gut and the diversity of metabolites available to influence host traits.

## INTRODUCTION

Microbiomes associated with animals are variable in taxonomic composition and impact on host traits (1–6). This has generated two linked challenges: to understand the causes of the variability and to predict how the microbiome may interact with other variables, such as host genotype, age, and activity, as well as diet, to determine host traits (7–11). Strategies to reduce and manage these highly complex interactions include the use of simplified microbial communities of defined taxonomic composition (12–14), and modeling to investigate patterns across larger scales of parameter space than is technically feasible by empirical study (15–19).

The gut microbiota of *Drosophila* is an attractive system to study taxonomically simple microbiomes because its microbiome is naturally of low diversity, generally with fewer than 20 bacterial species in laboratory culture, and microbiomes of defined composition can readily be generated and maintained (20, 21). In most laboratory cultures, the microbiome is dominated by acetic acid bacteria (AABs), generally of the genus *Acetobacter* (α-proteobacteria), and lactic acid bacteria (LABs) of the genus *Lactobacillus* (Firmicutes) (20), although some studies have reported strong representation of other taxa, including γ-proteobacteria (e.g. *Stenotrophomonas* spp.) and Lactobacilli (e.g. *Enterococcus* spp.) (22, 23). Studies on associations with a single bacterial isolate and with communities of 2-5 microbial taxa have demonstrated that both individual taxa and communities can, variously, contribute to B vitamin nutrition, reduce lipid content, influence development rates, lifespan and fecundity, and modulate olfactory and egg-laying behavior (14, 24–30). In several studies, these microbiome effects on host traits have been shown to vary with diet composition, and they have been linked to among-microbe interactions that influence both the abundance and metabolic activity of individual microbial taxa (14, 25, 29–31). However, the relationship between diet, community composition and metabolic function of the microbiome remains poorly-understood.

The goal of this study was twofold: first, to determine the combined effects of community composition and nutrient availability on the metabolic interactions among members of the *Drosophila* gut microbiota; and second to establish how these interactions shape the abundance of the microorganisms in the community and the production of metabolites that may influence host traits. We adopted a modeling approach, specifically to construct and analyze the metabolic model for every combination of 5 bacterial taxa isolated from the *Drosophila* gut microbiome: *Acetobacter fabarum, A. pomorum, A. tropicalis, Lactobacillus brevis,* and *L. plantarum.* This choice of taxa enabled us to examine interactions among species of different taxonomic relatedness and metabolic function: within the Acetobacteraceae, the closely-related *A. fabarum* and *A. pomorum* are assigned to a different subgroup of *Acetobacter* from *A. tropicalis* (32); and the homofermentative *L. plantarum* and heterofermentative *L. brevis* are members of different phylogenetic groups of *Lactobacillus* (33). Members of the genus *Acetobacter* and *Lactobacillus* are well represented across both field and lab *Drosophila* populations (34–37) and have been widely used to investigate the impact of gut microbes on host physiology (14, 38–40). The metabolic network for each species was reconstructed from annotated metabolism genes in the sequenced genome, and the networks for the different species in each community integrated into community models. We then applied the SteadyCom framework (41) to quantify the steady-state composition of each community and to predict the metabolic flux within and between the bacteria contributing to each community. We used SteadyCom because other community modeling approaches generally assume fixed community composition lack constraints that prevent fast-growing organisms from displacing other microbes in the community regardless of nutrient availability in their environment. SteadyCom applies more ecologically relevant constraints and imposes a single, time-averaged constant growth rate across all members of a community to ensure co-existence and stability as predicted to occur in animal guts. SteadyCom is also applicable to established constraint-based modeling approaches such as flux variability analysis, an important tool for determining the robustness of metabolic models in various simulation conditions. This approach enabled us to assess how community composition is influenced by antagonistic and mutualistic metabolic interactions, and to evaluate how among-microbe interactions can dictate overall metabolic outputs from the community.

## RESULTS

### Growth of *Drosophila* gut bacterial communities under different nutrient regimes

Our first analysis tested for growth *in silico* of the 31 possible communities of the five test bacteria under three nutrient regimes (Fig. 1). As predicted, all five single-species communities displayed growth in both the base medium, comprising the complete set of nutrients required for growth by all the bacteria, and the nutrient-rich medium, in which the complete set of nutrients was provided in excess. However, only two bacteria, the acetic acid bacteria (AAB) *A. pomorum* and *A. tropicalis*, grew on the minimal medium containing glucose, glycerol, ammonia, sulfate, and phosphate as primary sources of carbon, nitrogen, sulfur, and phosphorus respectively. Growth of the third AAB, *A. fabarum* was rescued by co-culture with any other AAB or with one of the two lactic acid bacteria (LAB), *L. plantarum*, but not with *L. brevis*. Similarly, growth of *L. plantarum* was rescued by co-culture with any AAB but not with *L. brevis*. The co-culture requirements of *L. brevis* were greater, requiring both *L. plantarum* and at least one AAB. The failure of *A. fabarum, L. plantarum,* and *L. brevis* to grow *in silico* on the minimal medium was a consequence of their auxotrophy for amino acids such as arginine and the peptidoglycan precursor diaminoheptanedioate. These metabolites are released from bacterial species, e.g. *A. pomorum* and *A. tropicalis*, that rescue growth.

**Fig. 1.**
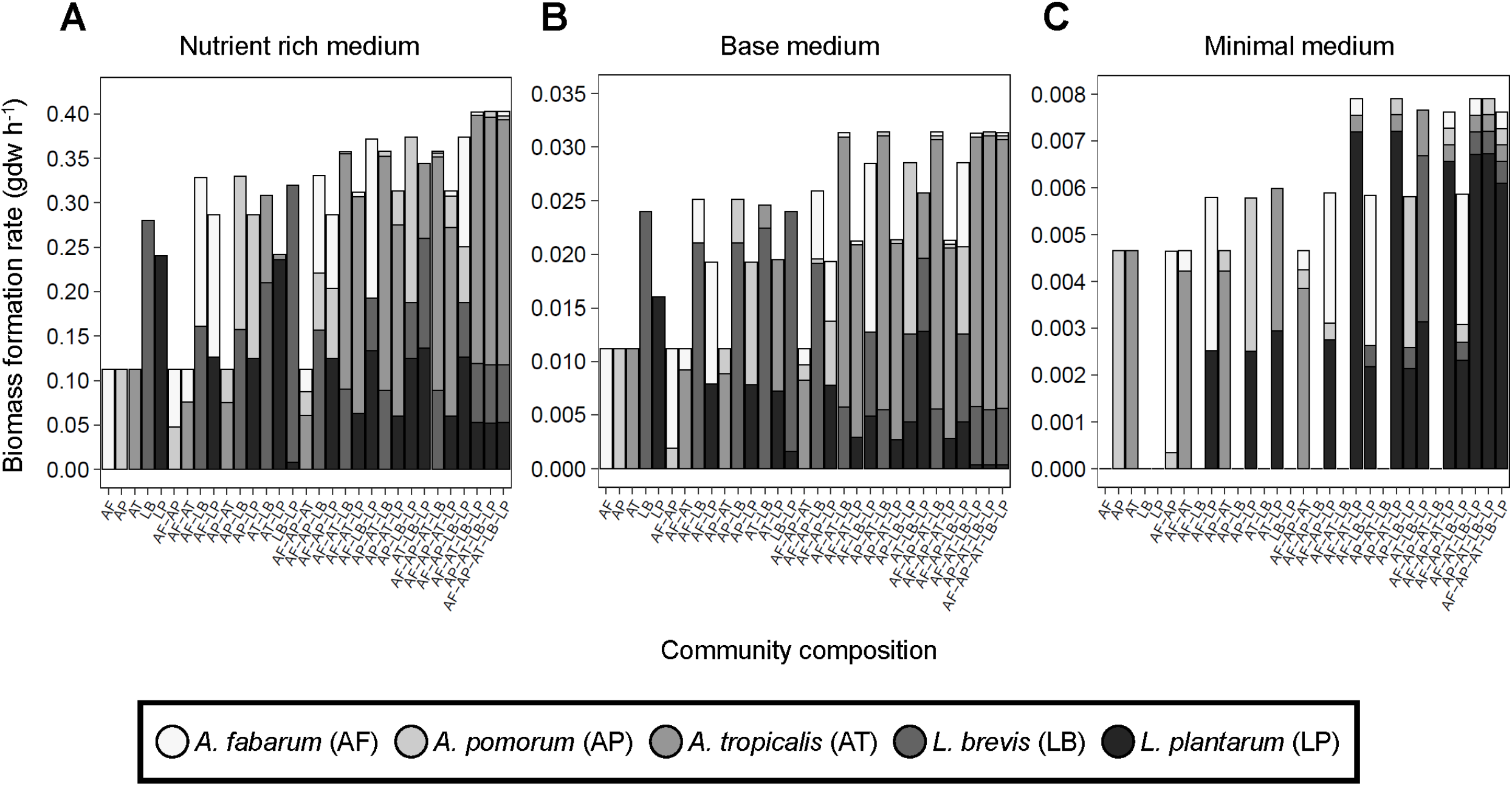
Bacterial growth dynamics on media of different nutrient content. Growth dynamics displayed as biomass formation rate predicted for growth in **(A)** nutrient rich medium, **(B)** base medium, and **(C)** minimal medium. Bar colors represents specific bacteria.

Further inspection of the data in Fig. 1 revealed the diverse effects of co-culture on growth dynamics of individual bacteria. For example, co-culture with *A. tropicalis* tended to reduce the growth of the other bacteria on the nutrient-rich medium; and the growth of AABs on the minimal medium was largely unaffected by co-culture with *L. plantarum* in two-member communities, but strongly repressed in communities of two or three AABs with *L. plantarum* (Table S1). To investigate these effects systematically, we classified co-culture interactions as sets of binary interactions: 1) competitive if both organisms displayed reduced growth in co-culture, 2) parasitic if the growth of one member was enhanced at the expense of another, 3) mutualistic if both organisms displayed increased growth in co-culture, 4) commensal if one organism displayed increased growth with no change in the other, 5) amensal if one organism displayed reduced growth with no change in the other, and 6) neutral if the growth of both organisms was unaltered. Increases and decreases in growth were determined by comparing the growth of a microbe in isolation to its growth in co-culture. In the nutrient-rich and basal media, the interactions were exclusively antagonistic: 50-70% of the interactions were competitive, and the remainder were parasitic (Fig. 2A, Table S1). In the minimal medium, parasitic interactions predominated, but competitive, mutualistic, amensal and neutral interactions also occurred (Fig. 2A). Notably, mutualistic interactions accounted for 12% of the interactions in two-member communities and increased progressively to 30% of the interactions in the five-member community. A large proportion (~80%) of neutral interactions occur between *A. fabarum* and *L. brevis* and these interactions switch to mutualistic interactions when *L. plantarum* is added to the community (Table S1C). Similarly, ~66% of amensal interactions occur between *L. brevis* and an at least one AAB (Table S1C). *L. brevis*-AAB amensal interactions switch to parasitic (L. brevis growth increases, AAB growth decreases) in more complex communities with the addition of *L. plantarum*. The switch in interaction type is facilitated by *L. plantarum* production of meso-2,6-diaminoheptanediote, an essential metabolite for *L. brevis* growth.

**Fig. 2.**
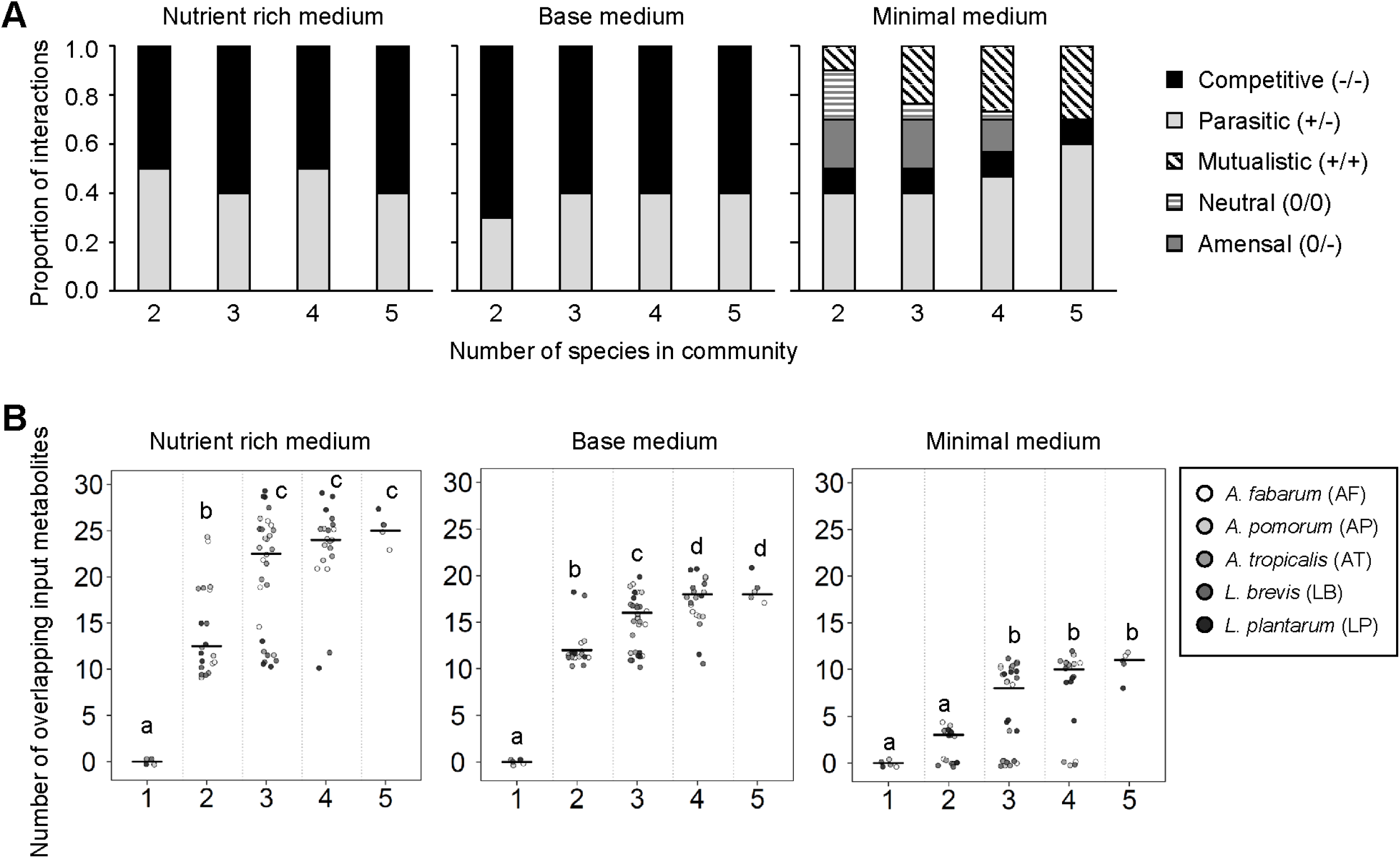
Ecological interactions in simulated communities of different diversity. **(A)** Impact of co-culture and medium on the sign of interactions between bacteria in the three test media…No commensal interactions were observed in any media type. **(B)** Overlapping metabolites consumed by bacteria in the 31 simulated communities. Symbol colors correspond to individual bacteria and the line connects the median value for the communities of different complexity. Significantly different (P < 0.05) groups by Tukey’s HSD posthoc test are indicated by different letters.

### Patterns of metabolite consumption and release

We hypothesized that the competitive interactions in the simulated bacterial communities (Fig. 2A) was underlain by the co-consumption of individual nutrients by two or more bacteria in the community; these nutrients may be constituents of the medium or, in communities with three or more members, derived from other bacteria. We, additionally, hypothesized that parasitic interactions involved the unidirectional cross-feeding of metabolites from the bacterium displaying depressed growth to the bacterium displaying increased growth in co-culture; while mutualistic interactions involved the reciprocal transfer of metabolites that were synthesized and released by one bacterium and required for growth by the other bacterium.

For our first analysis of the incidence of competition, we quantified the number of individual metabolites consumed by more than one bacterium in the simulated communities (Figure 2B, Table S2A-C, Dataset S1). As shown in Fig. 2B, the number of nutrients shared between two bacteria increased with both community complexity and nutrient content of the growth media. The number of shared metabolites increased significantly between two-member and three-member communities but increases in three to four and four to five member communities were mostly not significant for all media types (Table S2D). Consistent with the relatively low incidence of competitive interactions in the minimal medium, the greatest number of overlapping input metabolites recorded for any interaction in this medium was 12, which was half or less of the equivalent values, 21 and 29 for the basal medium and nutrient-rich medium, respectively (Fig. 2B).

To investigate the specific metabolic drivers of the antagonistic growth interactions, i.e. both competition and parasitism, we determined the input and output metabolites of each bacterium in every community. A total of 100 unique metabolites was predicted to be produced or consumed. We classified each metabolite by the frequency of its consumption by members of the gut community (Fig. 3, Table S3A-C). Our data show distinct metabolite use patterns associated with competitive, parasitic and mutualistic growth outcomes. Competitive growth interactions were significantly dominated by single-use or co-consumption in the rich and base media, but not in the minimal medium (Fig. 3A-C, Table S3D). Similarly, parasitic interactions were significantly dominated by single-use or co-consumption in rich and base media, and single-produced, and cross-fed metabolite use patterns dominated minimal medium parasitic interactions (Fig. 3D-F, Table S3D). Mutualistic growth interactions, observed only in the minimal medium, were characterized by the dominance of cross-feeding interactions which made up a significantly larger proportion of all interaction types (Fig. 3G, Table S3D).

**Fig. 3.**
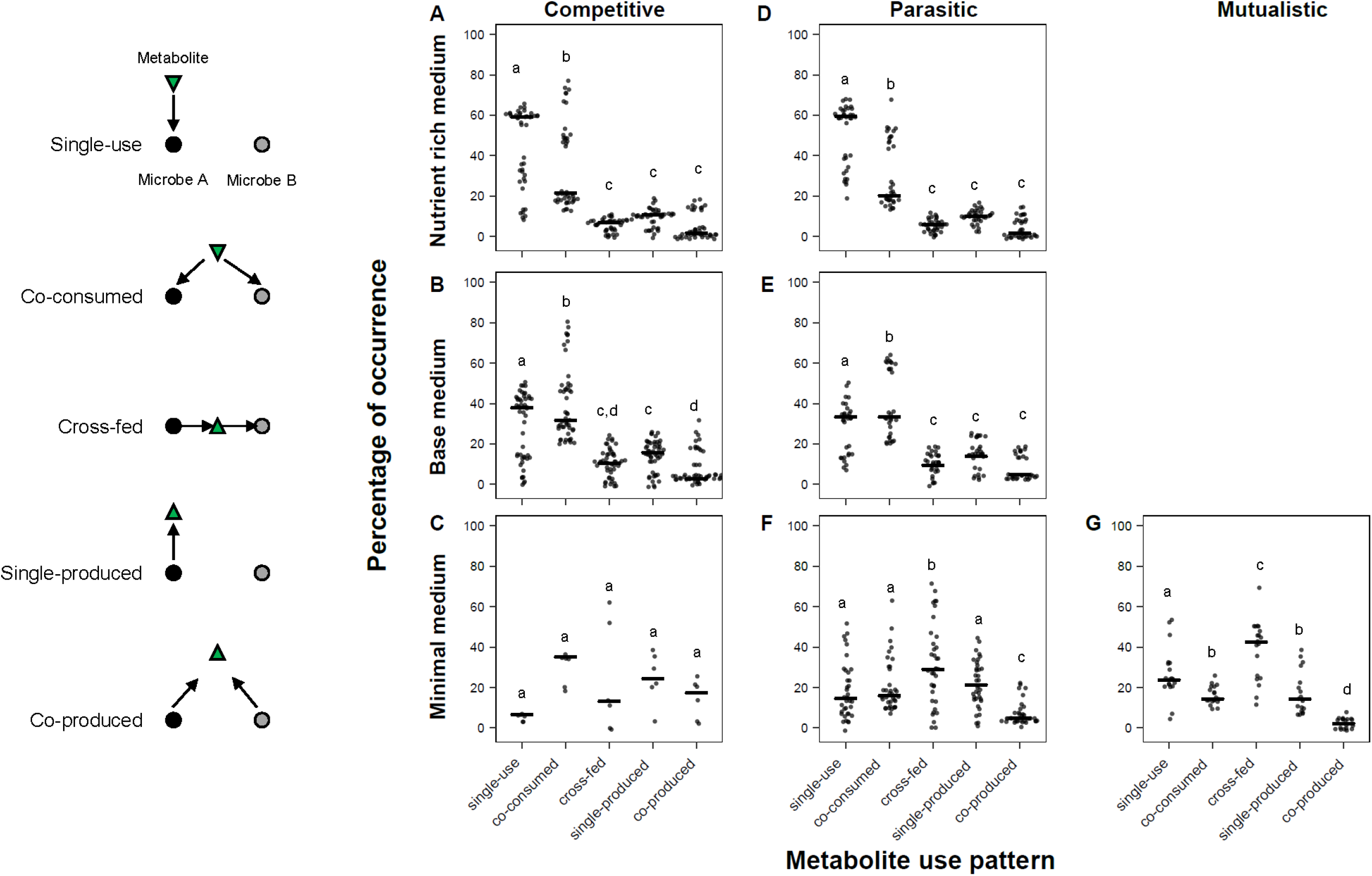
Metabolite use patterns associated with competitive, parasitic and mutualistic growth outcomes. Each dot represents the frequency of a metabolite use pattern associated between bacterial pairs in the 31 simulated communities. Black bar indicates median frequency of occurrence for each metabolite use pattern. Significantly different (P < 0.05) groups by Tukey’s HSD posthoc test are indicated by different letters.

Our data also show shifts in metabolite use profiles with depletion of nutrients in the growth medium. In the nutrient-rich medium, single-use consumption of metabolites predominated, and metabolites in the base and minimal media had more diverse metabolite use profiles with an increasing representation of cross-fed, single-produced and co-produced metabolites (Fig. 3).

We then investigated the identity of metabolites in the different metabolite use patterns. Metabolite groups with the highest number of co-consumed metabolites were amino acids and B vitamins (Fig. 4, Table S4A-C). Tyrosine, tryptophan, proline, phenylalanine, glutamine, asparagine and arginine were the most frequently co-consumed amino acids and biotin (B7) the most co-consumed B vitamin. However, some B vitamins and cofactors were cross-fed, notably thiamin (B1), pyridoxine 5-phosphate (B6), nicotinamide D-ribonucleotide, riboflavin (B2), tetrahydrofolate (B9) and coenzyme A (Fig. 4). Among the carbon compounds, only glucose and glycerol were consistently co-consumed in all three media. Other carbon compounds displayed more diverse use profiles that differed for each growth medium. For instance, malate and formate were exclusively produced in the nutrient-rich and base media and consumed or cross-fed in the minimal medium (Fig. 4); while lactate and 2,3 butanediol were exclusively produced in the nutrient-rich medium. Intermediates of the TCA cycle 2-oxoglutarate, succinate, succinyl-CoA, and fermentation products acetate, acetaldehyde and acetoin were cross-fed carbon at high frequency; and serine, glutamate, and glycine were the most cross-fed amino acids (Fig. 4). Among the nucleotides, the pyrimidine deoxyuridine 5’-phosphate (dUMP) simultaneously ranked as the most co-consumed and cross-fed nucleotide (Fig. 4).

**Fig. 4.**
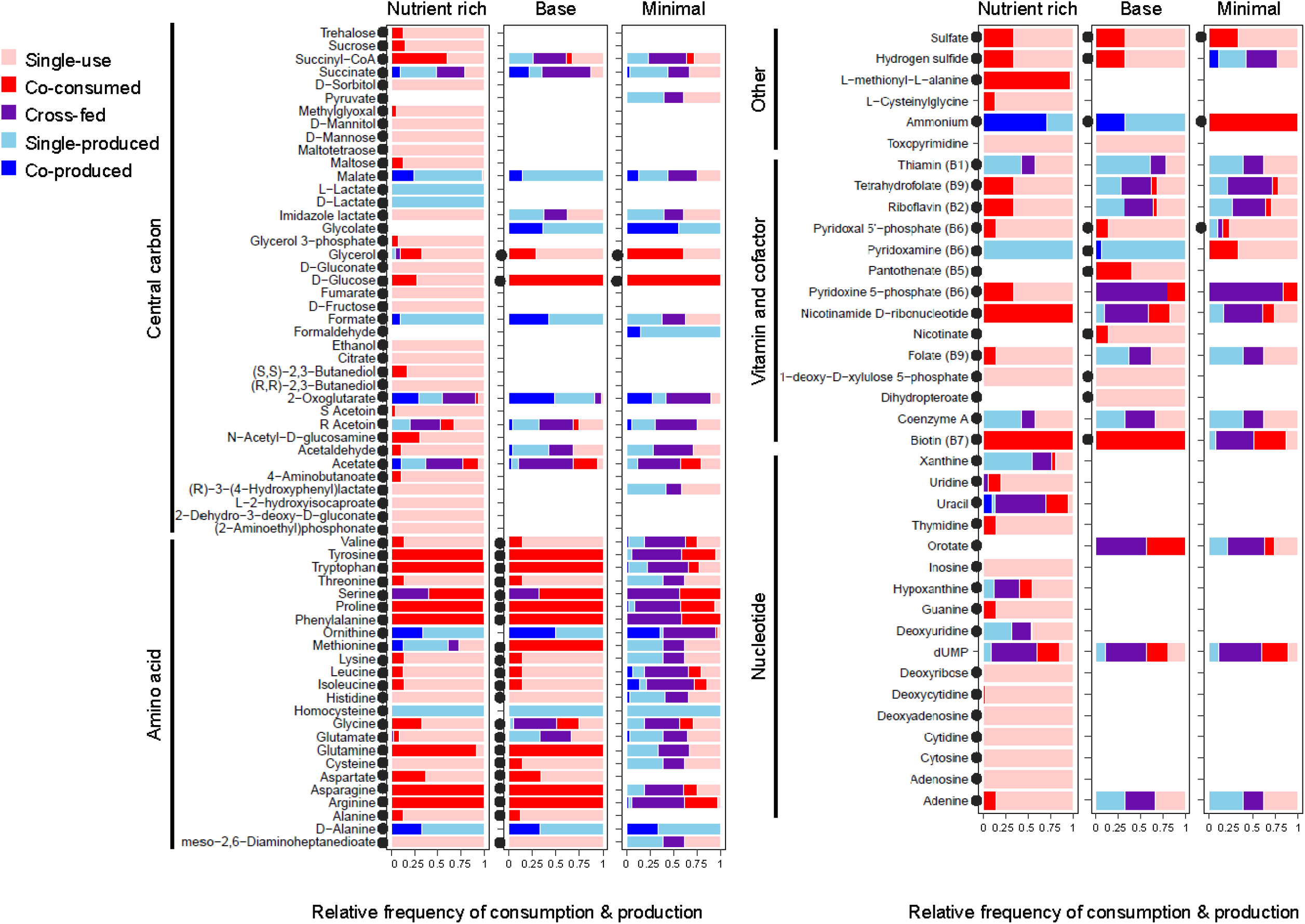
Metabolite use patterns for metabolite classes amino acid, carbon, nucleotide, vitamin and cofactors. Tick marks on X-axis indicate relative frequency of the consumption or production of a metabolite and range from 0 to 1 at 0.25 increments. The relative frequency of metabolite use is calculated by dividing the number of times a metabolite is used in a particular pattern (single-use, co-consumed, cross-fed, single-produced, co-produced) by the total number of times the metabolite is produced or consumed in the 31 simulated communities.

### Metabolic roles of individual bacteria

Our next analyses focused on metabolite production and consumption profiles of individual gut bacteria. Our simulations show that the metabolic role of individual gut bacteria as source or sink varies with the identity of co-culture microbe and across the three media for many metabolites, but is generally conserved across communities within the same media type (Fig. 5, Table S4D). For instance, all five gut bacteria produced ammonia in the nutrient-rich and base media but consumed ammonia in the minimal medium (Fig. 5A). . In base and rich media most microbes, except *A. tropicalis*, displayed committed roles for the production or consumption of all metabolites (Table S5A-C); *A. tropicalis* alternated roles as a producer and consumer for up 6% of all metabolites (Table S5A-C). In the minimal medium, all three acetic acid bacteria displayed variable roles as producers and consumers for up to ~30% of all metabolites (Table S5A-C). As an example, representatives of *Acetobacter* alternated as sources and sinks for acetate, arginine, ornithine, and succinyl-CoA in the minimal medium, depending on the number and identity of co-occurring bacteria (Fig.5B).

**Fig. 5.**
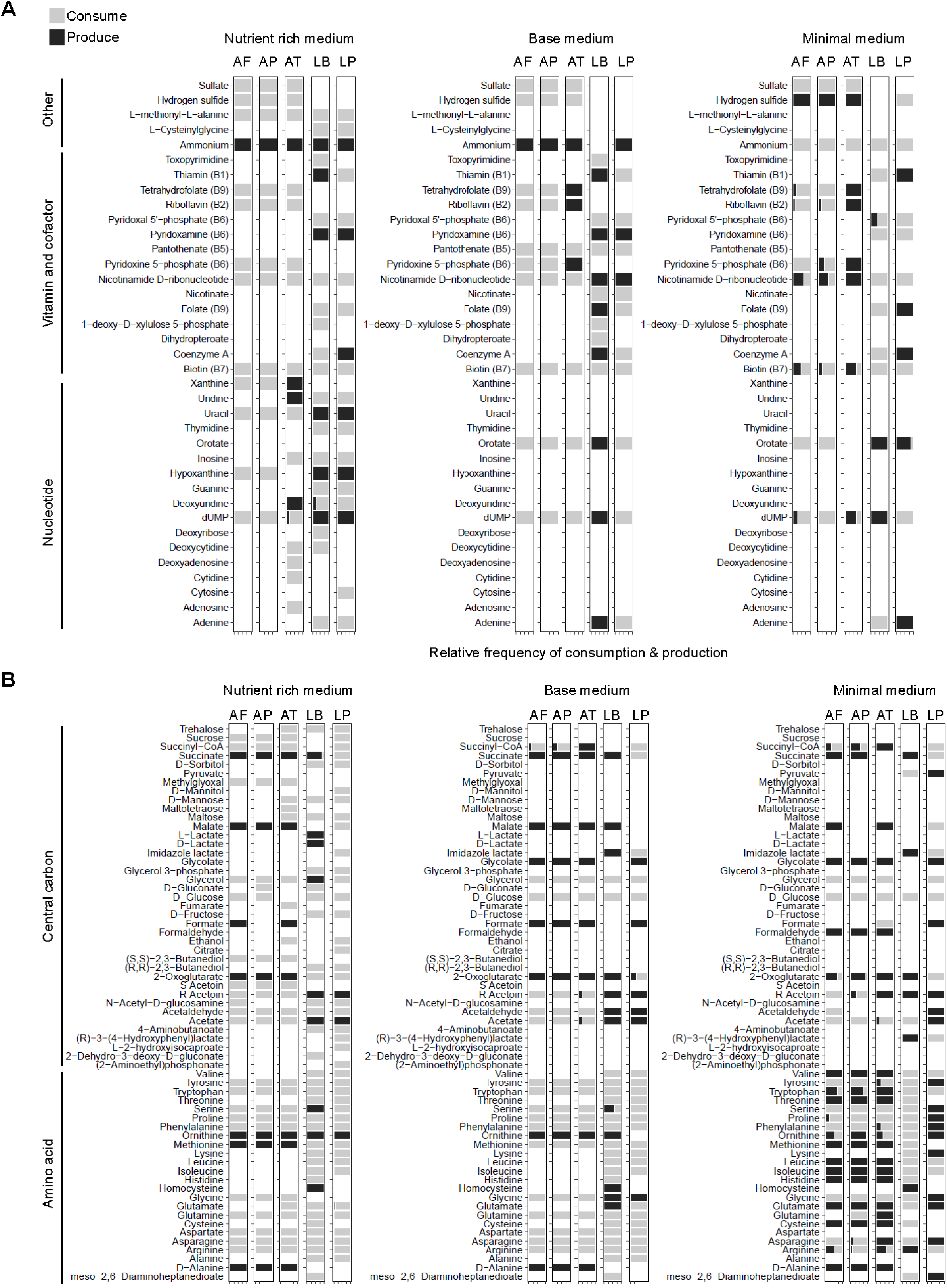
Metabolic roles of individual bacteria. Predicted metabolite production and consumption profiles for metabolite classes **(A)** nucleotide, vitamin and cofactors and **(B)** amino acid and carbon. Two letter abbreviations at the top of each plot represent individual bacteria: AF-*Acetobacter fabarum*, AP-*Acetobacter pomorum*, AT-*Acetobacter tropicalis*, LB-*Lactobacillus brevis*, LP-*Lactobacillus plantarum*. Tick marks on X-axis indicate relative frequency of the consumption or production of a metabolite and range from 0 to 1 at 0.25 increments.

Furthermore, the bacterial taxa had distinctive metabolic characteristics. All three *Acetobacter* species were sinks for glycine, serine, proline, acetaldehyde, methylglyoxal and producers of D-alanine, cysteine, histidine, isoleucine, leucine, valine, formaldehyde, malate, succinate (Fig. 5A), and *A. tropicalis* was, additionally, an important source of B vitamins including tetrahydrofolate (B9), riboflavin (B2), pyridoxine 5-phosphate (B6) and biotin (B7) (Fig. 5B). Both *Lactobacillus* species produced acetoin and were sinks for tryptophan and the branched-chain amino acids isoleucine, leucine, and valine. *L. brevis* was consistently a source of succinate, deoxyuridine 5’-phosphate (dUMP) and sink for most amino acids and meso-2,6-diaminoheptanediote (required for peptidoglycan synthesis). *L. plantarum* was a sink for arginine and succinate and source of pyruvate and meso-2,6-diaminoheptanediote.

### Effect of community size and taxa on metabolite richness

We next considered how the number of metabolites consumed or released by individual bacteria (i.e. metabolite richness) was influenced by the number and identity of other bacteria in the community. For the nutrient-rich and basal media, the number of taxa in a community did not significantly influence the number of metabolites consumed or released by individual taxa (Fig. 6A, Table S4D). However, on the minimal medium, as the number of taxa in a community increased, the number of metabolites consumed and released also increased (Fig. 6A).

**Fig. 6.**
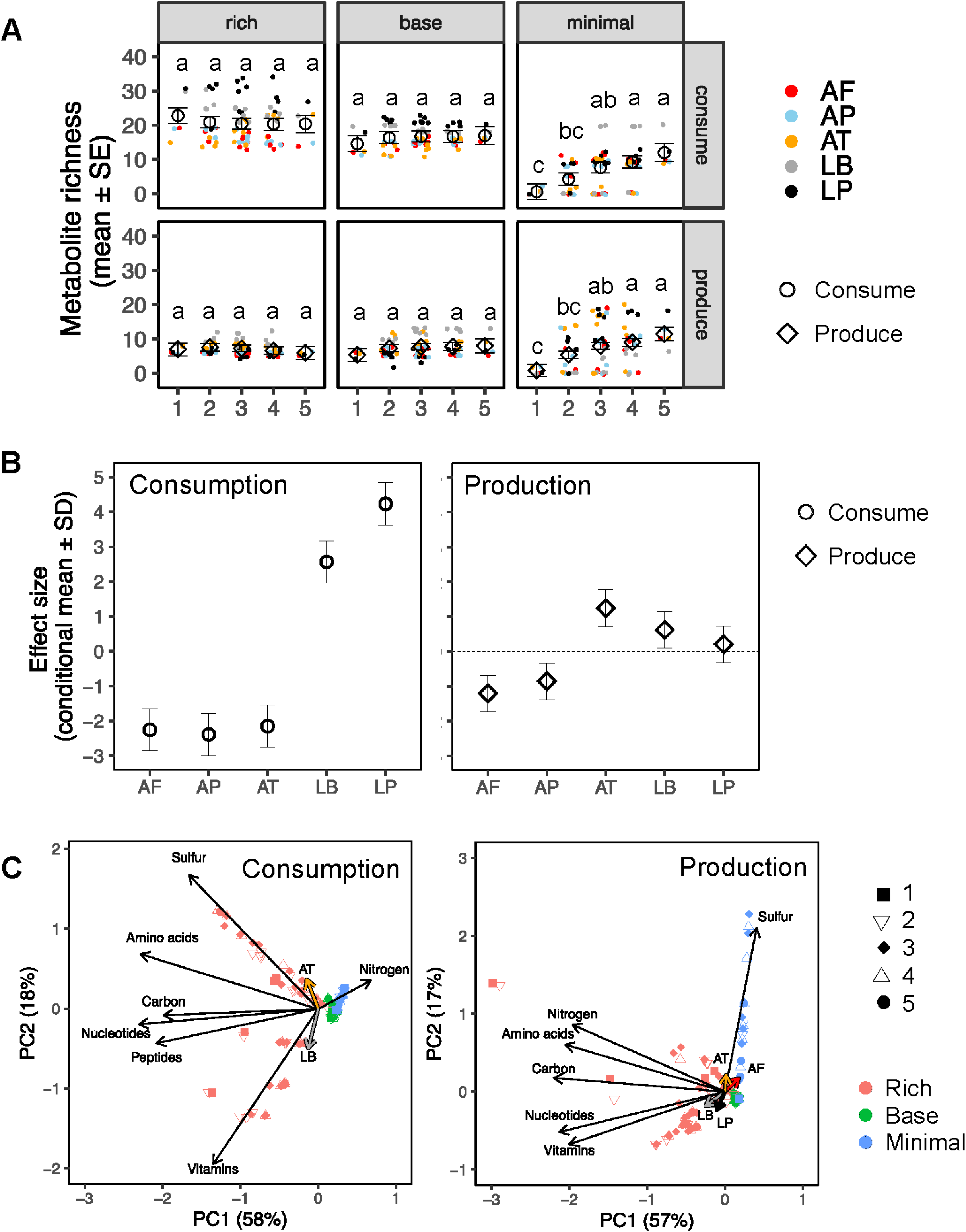
Metabolic function of simulated microbial taxa under different conditions. **(A)** Effect of community size on metabolite richness. Metabolite richness is calculated as the number of metabolites either consumed or released by a given taxon in each microbial treatment and medium combination. Effect of community size for each medium is indicated with the estimated marginal mean (open circles or diamonds) and standard error (SE) from ANOVA models. Letters indicate results from post hoc Tukey’s test, which was conducted separately for each medium. Closed, colored circles indicate individual metabolite richness values for each taxon under each condition. Effect test results are displayed in Table S4D. **(B)** Global effect of species identity on metabolite richness for the number of compounds consumed and released. The conditional mean and standard deviation are displayed for the best linear unbiased prediction. Dotted line indicates the grand mean for metabolite richness across all species. **(C)** Principal component analysis (PCA) correlating metabolite consumption or release rates with community size, medium type, and microbial presence. Black arrows indicate metabolite type scores, and colored arrow display the correlation vectors for microbial presence (only significant vectors are plotted). The percent variance explained by each axis is shown in parentheses.

The metabolic function of individual microbial taxa significantly varied across all media types (Table S4D). In particular, both *Lactobacillus* species consumed more metabolites than the three *Acetobacter* species. *A. tropicalis* released more metabolites on average than the other two *Acetobacter* species, and both *Lactobacillus* species released slightly more metabolites on average than the grand mean (Fig. 6B).

The types of metabolites consumed or released varied with the identity and number of species in the community (Fig. 6C, Table S5D). *A. tropicalis* and *L. brevis* had the strongest effect on consumption rates, especially in the nutrient-rich medium; *A. tropicalis* positively correlated with consumption of sulfur-containing metabolites, while *L. brevis* increased with vitamin consumption (Fig. 6C). For metabolite release, *Lactobacillus* species had the greatest impact in the nutrient-rich medium, correlated with vitamin and nucleotide release, while the *Acetobacter* species were more important in the minimal depleted medium and *A. tropicalis,* in particular, was strongly correlated with the release of sulfur-containing metabolites (Fig. 6C). *A. pomorum* consistently had no effect on consumption or production rates (Table S5D).

### Net outputs from the bacterial communities

Our final analysis addressed the metabolic products of the 31 bacterial communities in each of the three media. These products are candidate bacterial-derived metabolites that are available to the host animal. A total of 21 metabolites were predicted to be made available to the host (Table S6, Dataset S2), 19 of which were produced in the nutrient-rich medium, 12 in the base medium and five in the minimal medium. Some of the 21 metabolites have been demonstrated to previously play important roles in *Drosophila* physiology. For instance, acetate (26, 42, 43) and succinate (44) reduce host TAG levels, and microbe-derived amino acids rescue *Drosophila* growth on amino acid deficient diets (38, 45) and double *Drosophila* lifespan on low protein diets (46).

Three metabolites (acetate, D-alanine, homocysteine) were predicted to be made available to the host under all three diet conditions. In addition, the central carbon metabolites predicted to be released from bacteria in at least one medium include 2-oxoglutarate, formaldehyde, formate, glycolate, lactate, malate, and succinate.

## DISCUSSION

Our *in silico* study of metabolic interactions among *Drosophila* gut bacteria yielded two key findings. First, the pattern of metabolite consumption and release by individual bacteria and communities is dynamic, varying with nutrient conditions and community composition. Second, ecological interactions identified from the growth patterns of the bacteria (competition, mutualism, etc) are underpinned by a diversity of metabolic interactions, with evidence that the bacteria tend to compete for certain classes of nutrients (e.g. amino acids, B vitamins) more frequently than for others, particularly carbon sources.

Our modeling is based on several simplifying assumptions. In particular, we assume that the different bacterial species are in close proximity, such that among-species flux of metabolites is unimpeded. Although there is very limited information on the spatial organization of gut microorganisms in *Drosophila*, bacteria in other hosts can be planktonic in the gut lumen or adhere to the gut wall, often as single-species or structured multi-species colonies (47–50); and these different spatial patterns both affect and are influenced by abiotic conditions, nutrient availability, metabolite exchange and types of growth interactions between species (51–53). For example, commensal and neutral interactions which are likely to occur among spatially isolated microbes are not captured in most of our simulations. Furthermore, our models address primary metabolism exclusively and are not designed to investigate the effects of secondary metabolite mediated interactions, e.g. interference competition among bacteria mediated by toxins (54, 55), differential susceptibility of bacteria to host immune factors (56–59). Despite these limitations, many of our model outputs are consistent with published empirical data on the metabolic function of *Drosophila* gut bacteria, particularly the production by individual bacterial species of specific fermentation products (24, 60), amino acids (30, 61, 62) and B vitamins (29, 63, 64).

Furthermore, the predicted incidence of different ecological interactions, as deduced from biomass production in the various communities, agrees largely with published data reporting a predominance of antagonistic interactions among *Drosophila*-associated gut microbes (65) and other microbial communities (55, 66–68). Our observation that nutrient-poor conditions favor mutualistic interactions, especially in more complex communities, is also consistent with both predictions and empirical data for other microbial systems, e.g. (69–71). Taken together, these considerations indicate that our modeling approach is robust. It can be used with confidence to investigate patterns in the metabolic consequences of varying nutrient availability and community composition over a larger range of conditions than is technically realistic for empirical study.

Genome-scale metabolic modeling, as used here, brings into sharp focus the complexity of metabolic interactions among microorganisms. This provides a different perspective from empirical studies that, generally, focus on a single class of nutrients, e.g. short-chain fatty acids or B vitamins. In particular, the metabolic traits of an individual bacterium are not fixed but strongly influenced by the nutrient environment and the presence and identity of co-occurring microorganisms. For example, our models predict that the *Lactobacillus* species are net producers of uracil only under nutrient-rich conditions (Fig. 5). Bacterial-derived uracil has been shown to induce a pro-inflammatory state in the *Drosophila* gut via DUOX-mediated production of reactive oxygen species (72), and our data raise the possibility that the effect of *Lactobacillus* on the immunological status of the gut may be influenced by dietary factors. Similarly, the finding that the net production of several B vitamins by *Acetobacter* varies with the presence and identity of co-occurring bacteria (Fig. 5A) suggests that studies exclusively using associations with single bacterial taxa may not capture the full complexity of B vitamin provisioning by the *Drosophila* gut microbiome.

The complexity of the metabolic interactions among the bacterial species also has implications for the sign of ecological interactions. Various empirical analyses have demonstrated how an ecological interaction can be driven by a single metabolic interaction, e.g. competition for a single resource (73–75), mutualism by reciprocal cross-feeding of a pair of metabolites, each produced by one microorganism and required by the other (76, 77). However, as summarized in Fig. 3, the totality of the metabolic relationship between interacting microorganisms includes multiple classes of metabolic interaction. A relatively minor change in the uptake/release of metabolite(s), in response to a change in nutrient availability or community composition, could result in the transition to a different ecological relationship. For example, the switch from a competitive interaction to parasitism may contribute to the bloom of a microorganism previously held in check by competition (78).

Further elaboration of metabolic models, as used here and, for example by (68), offers the opportunity to investigate how subtle changes in metabolite flux in communities of different complexity and different nutrient regimes can lead to a switch between different ecological states of the microbiome and its interaction with the host.

The complexity and variation in metabolic interactions among the gut bacteria are, however, overlain by several broad patterns with respect to both metabolite class and bacterial species. Considering metabolites first, a key output of this study is that the pattern of metabolite use differed between substitutable and non-substitutable nutrient sources, i.e. where other resources can be utilized as an equivalent resource versus where no equivalent resource is available (79). Consistent with the generality that organisms tend to compete less for substitutable than non-substitutable resources (80), the *Drosophila* gut bacteria competed more for non-substitutable nutrients, such as amino acids and B vitamins, than substitutable nutrients, such as intermediates in central carbon metabolism. These patterns have implications for the health and wellbeing of the host. In particular, microbial consumption of dietary nutrients can deplete nutrient availability to the host (81), and among-microbe competition is predicted to alter host access to both critical nutrients and metabolites that influence the signaling pathways in *Drosophila* (26, 82) and other animals (83, 84).

The five bacteria selected for this analysis varied in their genetic capacity for metabolic function, particularly between the *Acetobacter* and *Lactobacillus* species (Fig. 6, see also (39)). Nevertheless, our modeling revealed substantial functional redundancy in metabolite production among the different bacteria: except for meso-2,6-diaminoheptanediote, produced by *L. plantarum*, we identified no metabolites produced exclusively by a single species. This metabolic redundancy is consistent with the evidence that taxonomically-different microbial communities can be functionally equivalent in *Drosophila* (85) and other animals (86, 87). The unique metabolic function of *L. plantarum* highlights the role of this bacterium in microbial community interactions, including its potential as a probiotic. The beneficial effects of *L. plantarum* in the *Drosophila* system have been attributed to its capacity to promote protein assimilation from the diet (82). Our observation that *L. plantarum* provisions cell wall constituents, B vitamins and amino acids for auxotrophic bacteria in the gut provides additional metabolic routes by which *L. plantarum* may promote the overall diversity of the gut microbiota.

In conclusion, this study demonstrates how *in silico* approaches can yield mechanistic insight into the metabolic traits of individual microbes and communities, and how these traits can influence metabolite levels that impact host physiology. Our models identify patterns by which microbial communities interact and respond to changes in nutrient input from the host and allow the generation of testable hypotheses for more targeted empirical studies. Substantial insight into how variations in microbiomes impact host health and metabolism have been gained from combining metabolic modeling with empirical studies (88, 89) and the simplicity of the *Drosophila* system presents an ideal model system to combine *in silico* and *in vivo* approaches to understand how gut-associated microbes impact host health.

## MATERIALS AND METHODS

### Generation of the individual bacterial metabolic models

Genomes of *Acetobacter fabarum* (JOPD01000000), *Acetobacter pomorum* (JOKL01000000), *Acetobacter tropicalis* (JOKM01000000), *Lactobacillus brevis* (JOKA01000000) and *Lactobacillus plantarum* (JOJT01000000) were downloaded from NCBI and re-annotated using the RAST annotation server (90). Two draft model reconstructions were generated for each genome and combined to generate a final model. The first models were obtained by performing reciprocal BLASTs of *Acetobacter* genomes against *Escherichia coli* str. K-12 substr. MG1655 and Lactobacilli genomes against *Lactobacillus plantarum* WCFS1. Gene orthologs identified from the reciprocal blast searches were compared to the *E. coli* str. K-12 substr. MG1655 metabolic model iML1515 (91) and the *Lactobacillus plantarum* WCFS1 metabolic model (92). Then, reactions encoded by these genes were manually extracted to create a draft model. The second draft reconstructions were generated from the automated reconstruction pipeline ModelSEED (93) using the RAST re-annotated genomes as input. For each bacterium, the two models were integrated and manually curated to remove redundant reactions and ensure correct reaction gene association, stoichiometry, and directionality. Organism-specific features and genes encoding metabolic reactions absent in *Escherichia coli* iML1515 and *Lactobacillus plantarum* WCFS1 metabolic models were identified by literature review and searches of BioCyc, KEGG, EcoCyc, BiGG and BRENDA databases (94–98), and then added to the draft model. The models were further curated using nutrient utilization BioLog data from (39) to verify and identify nutrient sources utilized by *Acetobacter* and *Lactobacillus*. All models were evaluated using MEMOTE (99).

### Model media composition

All simulations were performed in one of three media types; a minimal medium, a base medium and a rich medium. The minimal medium is a nutrient poor medium containing glucose, glycerol, ammonia, sulfate, and phosphate as primary sources of carbon, nitrogen, sulfur, and phosphorus, respectively. Components of the minimal medium were selected to investigate the complete scope of interaction possible between *Drosophila* gut microbiota in the absence of host gut and diet derived nutrients. Components of the base medium were selected to match nutrient auxotrophies for all 5 bacteria and allow the growth of all 5 bacteria in isolation. Components of the rich medium comprised the complete set of nutrients required for growth by all five bacteria, all major sources of carbon, nitrogen, sulfur, and phosphorus that all the bacteria have annotated transporters for, nutrient components of *Drosophila* holidic diet and metabolites predicted to be available in the fly gut from a metabolomic analysis (100). All media nutrient constituents and the flux bounds used for all bacteria and all simulations are provided in Table S7.

### Model constraints applied

Reaction fluxes for community members were obtained using the SteadyCom (41) Flux Variability Analysis (FVA) implementation in the OpenCOBRA Toolbox (101) with the following constraints. Medoid growth rate vectors of individual species were computed by performing FVA while maintaining 99.99% of the maximum community growth rate (*μ*_max_) and simultaneously maximizing and minimizing flux through individual reactions to obtain individual species growth rate values for each simulation. For each species, we obtained between 1000-5000 growth rate values, representing species growth rates when each reaction was performing at its minimum and maximum while maintaining the maximum community growth rate. We used the medoid predicted growth rate value for each species as a lower bound for all subsequent model simulations by constraining SteadyCom parameters BMcon, BMrhs, and BMCsense to require each species have a biomass value of at least the computed medoid in all simulations. A SteadyCom algorithm that minimizes the L1 (taxicab) norm of the predicted flux vectors was also applied to remove futile cycles and extraneous flux predictions.

### Model simulation

To find the maximum community growth rate we used the bisection algorithm in SteadyCom because it presented minimal convergence and feasibility issues with the constraints applied to our community model. Additionally, the default SteadyCom feasTol algorithm which sets the allowed error for determining if an input solution is feasible was set to 1e-8 for the solver. All parameters used to constrain SteadyCom simulations are available in the runSteadyCom and runSteadyComFVAMedoid functions.

### Analysis of Simulated Results

Simulations were run for each subset of the 5 species, i.e. 2^5^−1 = 31 communities, 5 of which are single-species communities. We constructed the single-member communities as multi-species models to facilitate the analysis and simulation pipelines, particularly SteadyCom, which requires a multi-species model. We verified that growth rates for single-species models and the single-member community models with flux balance analysis (102) were identical (see Tests/testMultiModelSingleton.m).

### Statistical analyses

All statistical analyses were performed using R v.3.6.1 (103) with an alpha of 0.05 to assess significance. Bonferroni correction for multiple tests was applied where required. Statistical differences between the number of overlapping metabolites in different community sizes and metabolite use patterns associated with different ecological interactions were investigated by one-way analysis of variance (ANOVA) followed by Tukey’s HSD post hoc test. Metabolite richness was calculated as the number of metabolites either consumed or released by a given taxon in each microbial treatment and medium combination. A mixed effect two-way ANOVA was used to assess how the medium type (rich, basal, or minimal) and community size (number of taxa) influenced metabolite richness using ‘lmer’ function in lme4 package (104). The data for metabolite consumption and release were analyzed separately. Medium type and community size were included as categorical fixed effects and microbial treatment (combination of microbes in a community, e.g. AF + LB) and taxon (AF, AP, AT, LB, or LP) were designated categorical random effects. The ‘Anova’ function in the car package (105) was used to perform a type III Wald’s F test to determine the effect of predictors with a Kenward-Rodger approximation to estimate residual degrees of freedom. A post hoc Tukey’s test was performed for each media type to determine pairwise differences across community size. An analysis of deviance was performed to assess the significance of each random effect and the best linear unbiased prediction was estimated for each taxon using the ‘ranef’ function to predict the global effect of each taxon across the different community size and medium combinations. Marginal and conditional R2 values were calculated using the MuMIn package (106).

A principal component analysis (PCA) was performed to correlate metabolite consumption or release rates with community size, medium type, and microbial presence using the vegan package (107). A correlation matrix was implemented using the total sum rate of metabolites either consumed or released for each of the metabolite type bins (carbon, amino acids, nitrogen, etc.). The function ‘envfit’ was used to correlate PC1 and PC2 with microbial presence with 999 permutations and significant vectors were plotted. In addition, a permutational multivariate analysis of variance (PERMANOVA) was used to assess the effect of community size by medium type as well as taxon on metabolite rates with the function ‘adonis’. Data was autoscaled and a Euclidean distance matrix was implemented for the model with 999 permutations. Consumption and release rates were analyzed separately for all analyses.

## Code and Data Availability

All code used in this study can be found at https://github.com/federatedcloud/DouglasMetabolicModels/releases/tag/v1.0.1. The simulations were performed using v3.0.4 of the OpenCOBRA Toolbox and v7.5.1 of the Gurobi Optimizer (108). An optional, containerized environment for running the code is available at https://github.com/federatedcloud/COBRAContainers. All results derived from simulations can be found in DouglasMetabolicModels/analysis/CMP_and_CooperativeFluxes. A tutorial is provided for performing simulations in the repository’s top-level README file. The code is available under the MPL2 license. SBML files of the models have been submitted to the BioModels database (109) with the following identifiers: MODEL2002040002, MODEL2002040003, MODEL2002040004, MODEL2002040005 and MODEL2002040006.

## ACKNOWLEDGEMENTS

We thank the Aristotle Cloud Federation (NSF OAC-1541215) for computing resources for model simulation, and Dr. Lynn Johnson (Cornell Statistical Consulting Unit) for statistical advice. This research was funded by an NIH grant R01GM095372 to AED.

There is no conflict of interest for all authors.

## Author contributions

NYDA, BEB, and AED designed research; NYDA and JS created metabolic models; NYDA, BEB and CW performed model simulations and analysis; JGM performed statistical analysis; NYDA and AED wrote the first draft of the paper, and manuscript revisions were made by all authors.

## Supplementary Tables

**Table S1.** Predicted changes to bacterial growth (gdw h-1) in co-culture compared to growth in monoculture **A)** rich medium, **B)** base medium, **C)** minimal medium.

**Table S2.** Predicted number of inputs and outputs from bacteria **A)** rich medium, **B)** base medium, **C)** minimal medium. **D)** Summary statistics for Figure 2B

**Table S3.** Predicted community metabolite use patterns for competitive, parasitic and mutualistic interactions **A)** rich medium, **B)** base medium, **C)** minimal medium. **D)** Summary statistics for Figure 3.

**Table S4.** Metabolite use pattern **A)** rich medium, **B)** base medium, **C)** minimal medium. **D)** Effect of community size and medium type on metabolite richness. Tests with significant p values are shown in bold.

**Table S5.** Predicted total number of times metabolite is consumed or produced by individual bacteria in all simulations **A)** rich medium, **B)** base medium, **C)** minimal medium. **D)** Effect of taxa, community size, and medium type on metabolite consumption and release rates. Tests with significant p values are shown in bold.

**Table S6.** Metabolites predicted to be available to the host.

**Table S7.** List of components **A)** rich medium, **B)** base medium, **C)** minimal medium.

## Datasets

**Dataset S1. A)** Metabolite abbreviation key, **B)** Predicted number of inputs and outputs from bacteria - rich medium, **C)** Predicted number of inputs and outputs from bacteria - base medium, **D)** Predicted number of inputs and outputs from bacteria - minimal medium.

**Dataset S2.** Predicted metabolite flux available to to the host.

## REFERENCES

1. Adair KL, Douglas AE. 2017. Making a microbiome: the many determinants of host-associated microbial community composition. Curr Opin Microbiol 35:23–29.

2. Costello EK, Stagaman K, Dethlefsen L, Bohannan BJ, Relman DA. 2012. The application of ecological theory toward an understanding of the human microbiome. Science 336:1255–1262.

3. Douglas AE. 2018. Fundamentals of Microbiome Science: How Microbes Shape Animal Biology. Princeton University Press.

4. Lozupone CA, Stombaugh JI, Gordon JI, Jansson JK, Knight R. 2012. Diversity, stability and resilience of the human gut microbiota. Nature 489:220–230.

5. McFall-Ngai M, Hadfield MG, Bosch TCG, Carey HV, Domazet-Lošo T, Douglas AE, Dubilier N, Eberl G, Fukami T, Gilbert SF, Hentschel U, King N, Kjelleberg S, Knoll AH, Kremer N, Mazmanian SK, Metcalf JL, Nealson K, Pierce NE, Rawls JF, Reid A, Ruby EG, Rumpho M, Sanders JG, Tautz D, Wernegreen JJ. 2013. Animals in a bacterial world, a new imperative for the life sciences. Proc Natl Acad Sci USA 110:3229–3236.

6. Ankrah NY, Douglas AE. 2018. Nutrient factories: metabolic function of beneficial microorganisms associated with insects. Environ Microbiol 20:2002–2011.

7. Aleman FDD, Valenzano DR. 2019. Microbiome evolution during host aging. PLoS Pathog 15: e1007727.

8. Dobson AJ, Chaston JM, Newell PD, Donahue L, Hermann SL, Sannino DR, Westmiller S, Wong AC-N, Clark AG, Lazzaro BP. 2015. Host genetic determinants of microbiota-dependent nutrition revealed by genome-wide analysis of Drosophila melanogaster. Nature Comm 6:6312.

9. Goodrich JK, Davenport ER, Beaumont M, Jackson MA, Knight R, Ober C, Spector TD, Bell JT, Clark AG, Ley RE. 2016. Genetic determinants of the gut microbiome in UK twins. Cell Host Microbe 19:731–743.

10. Lamoureux EV, Grandy SA, Langille MG. 2017. Moderate exercise has limited but distinguishable effects on the mouse microbiome. mSystems 2:e00006–17.

11. Zmora N, Suez J, Elinav E. 2019. You are what you eat: diet, health and the gut microbiota. Nature Rev Gastroent Hepatol 16:35–56.

12. Wymore Brand M, Wannemuehler MJ, Phillips GJ, Proctor A, Overstreet A-M, Jergens AE, Orcutt RP, Fox JG. 2015. The altered Schaedler flora: continued applications of a defined murine microbial community. ILAR J 56:169–178.

13. Elzinga J, van der Oost J, de Vos WM, Smidt H. 2019. The use of defined microbial communities to model host-microbe interactions in the human gut. Microbiol Mol Biol Rev 83:e00054–18.

14. Gould AL, Zhang V, Lamberti L, Jones EW, Obadia B, Korasidis N, Gavryushkin A, Carlson JM, Beerenwinkel N, Ludington WB. 2018. Microbiome interactions shape host fitness. Proc Natl Acad Sci USA 115:E11951–E11960.

15. Ankrah NY, Chouaia B, Douglas AE. 2018. The cost of metabolic interactions in symbioses between insects and bacteria with reduced genomes. mBio 9:e01433–18.

16. Baldini F, Heinken A, Heirendt L, Magnusdottir S, Fleming RM, Thiele I. 2019. The microbiome modeling toolbox: from microbial interactions to personalized microbial communities. Bioinformatics 35:2332–2334.

17. Bauer E, Thiele I. 2018. From network analysis to functional metabolic modeling of the human gut microbiota. mSystems 3:e00209–17.

18. Levy R, Borenstein E. 2013. Metabolic modeling of species interaction in the human microbiome elucidates community-level assembly rules. Proc Natl Acad Sci USA 110:12804–12809.

19. Noecker C, Chiu H-C, McNally CP, Borenstein E. 2019. Defining and evaluating microbial contributions to metabolite variation in microbiome-metabolome association studies. mSystems 4:e00579–19.

20. Broderick NA, Lemaitre B. 2012. Gut-associated microbes of Drosophila melanogaster. Gut Microbes 3:307–321.

21. Douglas AE. 2019. Simple animal models for microbiome research. Nature Rev Microbiol 17:764–775.

22. Chaston JM, Newell PD, Douglas AE. 2014. Metagenome-wide association of microbial determinants of host phenotype in Drosophila melanogaster. mBio 5:e01631–14.

23. Cox CR, Gilmore MS. 2007. Native microbial colonization of Drosophila melanogaster and its use as a model of Enterococcus faecalis pathogenesis. Infect Immun 75:1565–1576.

24. Fischer C, Trautman EP, Crawford JM, Stabb EV, Handelsman J, Broderick NA. 2017. Metabolite exchange between microbiome members produces compounds that influence Drosophila behavior. eLife 6:e18855.

25. Newell PD, Douglas AE. 2014. Interspecies interactions determine the impact of the gut microbiota on nutrient allocation in Drosophila melanogaster. Appl Environ Microbiol 80:788–796.

26. Shin SC, Kim S-H, You H, Kim B, Kim AC, Lee K-A, Yoon J-H, Ryu J-H, Lee W-J. 2011. Drosophila microbiome modulates host developmental and metabolic homeostasis via insulin signaling. Science 334:670–674.

27. Storelli G, Defaye A, Erkosar B, Hols P, Royet J, Leulier F. 2011. Lactobacillus plantarum promotes Drosophila systemic growth by modulating hormonal signals through TOR-dependent nutrient sensing. Cell Metab 14:403–414.

28. Wong AC-N, Wang Q-P, Morimoto J, Senior AM, Lihoreau M, Neely GG, Simpson SJ, Ponton F. 2017. Gut microbiota modifies olfactory-guided microbial preferences and foraging decisions in Drosophila. Curr Biol 27:2397–2404. e4.

29. Wong AC-N, Dobson AJ, Douglas AE. 2014. Gut microbiota dictates the metabolic response of Drosophila to diet. J Exp Biol 217:1894–1901.

30. Consuegra J, Grenier T, Baa-Puyoulet P, Rahioui I, Akherraz H, Gervais H, Parisot N, da Silva P, Charles H, Calevro F, Leulier F. 2020. Drosophila-associated bacteria differentially shape the nutritional requirements of their host during juvenile growth. PLoS Biol 18:e3000681.

31. Sommer AJ, Newell PD. 2019. Metabolic basis for mutualism between gut bacteria and its impact on the Drosophila melanogaster host. Appl Environ Microbiol 85:e01882–18.

32. Cleenwerck I, Gonzalez A, Camu N, Engelbeen K, De Vos P, De Vuyst L. 2008. Acetobacter fabarum sp. nov., an acetic acid bacterium from a Ghanaian cocoa bean heap fermentation. Int J Syst Evol Microbiol 58:2180–2185.

33. Zheng J, Ruan L, Sun M, Gänzle M. 2015. A genomic view of lactobacilli and pediococci demonstrates that phylogeny matches ecology and physiology. Appl Environ Microbiol 81:7233–7243.

34. Chaston JM, Dobson AJ, Newell PD, Douglas AE. 2016. Host genetic control of the microbiota mediates the Drosophila nutritional phenotype. Appl Environ Microbiol 82:671–679.

35. Wang Y, Kapun M, Waidele L, Kuenzel S, Bergland AO, Staubach F. 2020. Common structuring principles of the Drosophila melanogaster microbiome on a continental scale and between host and substrate. Environ Microbiol Rep 12:220–228.

36. Douglas AE. 2018. The Drosophila model for microbiome research. Lab Anim 47:157–164.

37. Adair KL, Wilson M, Bost A, Douglas AE. 2018. Microbial community assembly in wild populations of the fruit fly Drosophila melanogaster. ISME J 12:959–972.

38. Consuegra J, Grenier T, Baa-Puyoulet P, Rahioui I, Akherraz H, Gervais H, Parisot N, da Silva P, Charles H, Calevro F. 2020. Drosophila-associated bacteria differentially shape the nutritional requirements of their host during juvenile growth. PLoS Biol 18:e3000681.

39. Newell PD, Chaston JM, Wang Y, Winans NJ, Sannino DR, Wong AC, Dobson AJ, Kagle J, Douglas AE. 2014. In vivo function and comparative genomic analyses of the Drosophila gut microbiota identify candidate symbiosis factors. Front Microbiol 5:576.

40. Sannino DR, Dobson AJ, Edwards K, Angert ER, Buchon N. 2018. The Drosophila melanogaster gut microbiota provisions thiamine to its host. mBio 9:e00155–18.

41. Chan SHJ, Simons MN, Maranas CD. 2017. SteadyCom: Predicting microbial abundances while ensuring community stability. PLoS Comput Biol 13:e1005539.

42. McMullen JG, Peters-Schulze G, Cai J, Patterson AD, Douglas AE. 2020. How gut microbiome interactions affect nutritional traits of Drosophila melanogaster. J Exp Biol 223: jeb227843.

43. Kamareddine L, Robins WP, Berkey CD, Mekalanos JJ, Watnick PI. 2018. The Drosophila immune deficiency pathway modulates enteroendocrine function and host metabolism. Cell Metab 28:449–462.e5.

44. Zhang FQ, McMullen JG, Douglas AE, Ankrah NYD. accepted. Succinate: a microbial product that modulates Drosophila nutritional physiology. Insect Science.

45. Consuegra J, Grenier T, Akherraz H, Rahioui I, Gervais H, da Silva P, Leulier F. 2020. Metabolic cooperation among commensal bacteria supports Drosophila juvenile growth under nutritional stress. iScience 23:101232.

46. Leitão-Gonçalves R, Carvalho-Santos Z, Francisco AP, Fioreze GT, Anjos M, Baltazar C, Elias AP, Itskov PM, Piper MD, Ribeiro C. 2017. Commensal bacteria and essential amino acids control food choice behavior and reproduction. PLoS Biol 15:e2000862.

47. Pedron T, Mulet C, Dauga C, Frangeul L, Chervaux C, Grompone G, Sansonetti PJ. 2012. A crypt-specific core microbiota resides in the mouse colon. mBio 3:e00116–12.

48. Yasuda K, Oh K, Ren B, Tickle TL, Franzosa EA, Wachtman LM, Miller AD, Westmoreland SV, Mansfield KG, Vallender EJ, Miller GM, Rowlett JK, Gevers D, Huttenhower C, Morgan XC. 2015. Biogeography of the intestinal mucosal and lumenal microbiome in the rhesus macaque. Cell Host Microbe 17:385–391.

49. Mark Welch JL, Hasegawa Y, McNulty NP, Gordon JI, Borisy GG. 2017. Spatial organization of a model 15-member human gut microbiota established in gnotobiotic mice. Proc Natl Acad Sci USA 114:E9105–E9114.

50. Schlomann BH, Wiles TJ, Wall ES, Guillemin K, Parthasarathy R. 2018. Bacterial Cohesion Predicts Spatial Distribution in the Larval Zebrafish Intestine. Biophys J 115:2271–2277.

51. Borer B, Tecon R, Or D. 2018. Spatial organization of bacterial populations in response to oxygen and carbon counter-gradients in pore networks. Nat Commun 9:769.

52. Bridier A, Piard JC, Pandin C, Labarthe S, Dubois-Brissonnet F, Briandet R. 2017. Spatial organization plasticity as an adaptive driver of surface microbial communities. Front Microbiol 8:1364.

53. Nadell CD, Drescher K, Foster KR. 2016. Spatial structure, cooperation and competition in biofilms. Nat Rev Microbiol 14:589–600.

54. Granato ET, Meiller-Legrand TA, Foster KR. 2019. The Evolution and Ecology of Bacterial Warfare. Curr Biol 29:R521–R537.

55. Hibbing ME, Fuqua C, Parsek MR, Peterson SB. 2010. Bacterial competition: surviving and thriving in the microbial jungle. Nat Rev Microbiol 8:15–25.

56. Earley AM, Graves CL, Shiau CE. 2018. Critical role for a subset of intestinal macrophages in shaping gut microbiota in adult zebrafish. Cell Rep 25:424–436.

57. Broderick NA. 2016. Friend, foe or food? Recognition and the role of antimicrobial peptides in gut immunity and Drosophila-microbe interactions. Phil Trans R Soc Lond B Biol Sci 371:20150295.

58. Liu X, Hodgson JJ, Buchon N. 2017. Drosophila as a model for homeostatic, antibacterial, and antiviral mechanisms in the gut. PLoS Pathog 13:e1006277.

59. Pabst O, Slack E. 2020. IgA and the intestinal microbiota: the importance of being specific. Mucosal Immunol 13:12–21.

60. Kim G, Huang J, McMullen II JG, Newell PD, Douglas AE. 2017. Physiological responses of insects to microbial fermentation products: insights from the interactions between Drosophila and acetic acid. J Insect Physiol 106:13–19.

61. Leitao-Goncalves R, Carvalho-Santos Z, Francisco AP, Fioreze GT, Anjos M, Baltazar C, Elias AP, Itskov PM, Piper MDW, Ribeiro C. 2017. Commensal bacteria and essential amino acids control food choice behavior and reproduction. PLoS Biol 15:e2000862.

62. Judd AM, Matthews MK, Hughes R, Veloz M, Sexton CE, Chaston JM. 2018. Bacterial methionine metabolism genes influence Drosophila melanogaster starvation resistance. Appl Environ Microbiol 84:e00662–18.

63. Blatch S, Meyer KW, Harrison JF. 2010. Effects of dietary folic acid level and symbiotic folate production on fitness and development in the fruit fly Drosophila melanogaster. Fly 4:312–319.

64. Sannino DR, Dobson AJ, Edwards K, Angert ER, Buchon N. 2018. The Drosophila melanogaster gut microbiota provisions thiamine to its host. mBio 9:e00155–18.

65. Gould AL, Zhang V, Lamberti L, Jones EW, Obadia B, Korasidis N, Gavryushkin A, Carlson JM, Beerenwinkel N, Ludington WB. 2018. Microbiome interactions shape host fitness. Proc Natl Acad Sci USA 115:E11951–E11960.

66. Foster KR, Bell T. 2012. Competition, not cooperation, dominates interactions among culturable microbial species. Curr Biol 22:1845–50.

67. Faust K, Bauchinger F, Laroche B, de Buyl S, Lahti L, Washburne AD, Gonze D, Widder S. 2018. Signatures of ecological processes in microbial community time series. Microbiome 6:120.

68. Venturelli OS, Carr AC, Fisher G, Hsu RH, Lau R, Bowen BP, Hromada S, Northen T, Arkin AP. 2018. Deciphering microbial interactions in synthetic human gut microbiome communities. Mol Syst Biol 14:e8157.

69. Zelezniak A, Andrejev S, Ponomarova O, Mende DR, Bork P, Patil KR. 2015. Metabolic dependencies drive species co-occurrence in diverse microbial communities. Proc Natl Acad Sci USA 112:6449–54.

70. Freilich S, Zarecki R, Eilam O, Segal ES, Henry CS, Kupiec M, Gophna U, Sharan R, Ruppin E. 2011. Competitive and cooperative metabolic interactions in bacterial communities. Nat Commun 2:589.

71. Hoek TA, Axelrod K, Biancalani T, Yurtsev EA, Liu J, Gore J. 2016. Resource availability modulates the cooperative and competitive nature of a microbial cross-feeding mutualism. PLoS Biol 14:e1002540.

72. Lee KA, Kim SH, Kim EK, Ha EM, You H, Kim B, Kim MJ, Kwon Y, Ryu JH, Lee WJ. 2013. Bacterial-derived uracil as a modulator of mucosal immunity and gut-microbe homeostasis in Drosophila. Cell 153:797–811.

73. MacLean RC, Gudelj I. 2006. Resource competition and social conflict in experimental populations of yeast. Nature 441:498–501.

74. Vulic M, Kolter R. 2001. Evolutionary cheating in Escherichia coli stationary phase cultures. Genetics 158:519–26.

75. Kehe J, Kulesa A, Ortiz A, Ackerman CM, Thakku SG, Sellers D, Kuehn S, Gore J, Friedman J, Blainey PC. 2019. Massively parallel screening of synthetic microbial communities. Proc Natl Acad Sci USA 116:12804–12809.

76. Harcombe W. 2010. Novel cooperation experimentally evolved between species. Evolution 64:2166–72.

77. D’Souza G, Shitut S, Preussger D, Yousif G, Waschina S, Kost C. 2018. Ecology and evolution of metabolic cross-feeding interactions in bacteria. Nat Prod Rep 35:455–488.

78. Selber-Hnatiw S, Rukundo B, Ahmadi M, Akoubi H, Al-Bizri H, Aliu AF, Ambeaghen TU, Avetisyan L, Bahar I, Baird A, Begum F, Ben Soussan H, Blondeau-Ethier V, Bordaries R, Bramwell H, Briggs A, Bui R, Carnevale M, Chancharoen M, Chevassus T, Choi JH, Coulombe K, Couvrette F, D’Abreau S, Davies M, Desbiens MP, Di Maulo T, Di Paolo SA, Do Ponte S, Dos Santos Ribeiro P, Dubuc-Kanary LA, Duncan PK, Dupuis F, El-Nounou S, Eyangos CN, Ferguson NK, Flores-Chinchilla NR, Fotakis T, Gado Oumarou HDM, Georgiev M, Ghiassy S, Glibetic N, Gregoire Bouchard J, Hassan T, Huseen I, Ibuna Quilatan MF, Iozzo T, Islam S, Jaunky DB, Jeyasegaram A, et al. 2017. Human gut microbiota: toward an ecology of disease. Front Microbiol 8:1265.

79. Tilman D. 1982. Resource Competition and Community Structure. Princeton University Press, Princeton, NJ.

80. Leon JA, Tumpson DB. 1975. Competition between two species for two complementary or substitutable resources. J Theo Biol 50:185–201.

81. Huang JH, Douglas AE. 2015. Consumption of dietary sugar by gut bacteria determines Drosophila lipid content. Biol Lett 11:20150469.

82. Storelli G, Defaye A, Erkosar B, Hols P, Royet J, Leulier F. 2011. Lactobacillus plantarum promotes Drosophila systemic growth by modulating hormonal signals through TOR-dependent nutrient sensing. Cell Metab 14:403–14.

83. Levy M, Thaiss CA, Elinav E. 2016. Metabolites: messengers between the microbiota and the immune system. Genes Dev 30:1589–97.

84. Rowland I, Gibson G, Heinken A, Scott K, Swann J, Thiele I, Tuohy K. 2018. Gut microbiota functions: metabolism of nutrients and other food components. Eur J Nutr 57:1–24.

85. Kang D, Douglas AE. 2020. Functional traits of the gut microbiome correlated with host lipid content in a natural population of Drosophila melanogaster. Biol Lett 16:20190803.

86. Lozupone CA, Stombaugh JI, Gordon JI, Jansson JK, Knight R. 2012. Diversity, stability and resilience of the human gut microbiota. Nature 489:220–30.

87. Moya A, Ferrer M. 2016. Functional redundancy-induced stability of gut microbiota subjected to disturbance. Trends Microbiol 24:402–413.

88. Heinken A, Ravcheev DA, Baldini F, Heirendt L, Fleming RMT, Thiele I. 2019. Systematic assessment of secondary bile acid metabolism in gut microbes reveals distinct metabolic capabilities in inflammatory bowel disease. Microbiome 7:75.

89. Henson MA, Orazi G, Phalak P, O’Toole GA. 2019. Metabolic modeling of cystic fibrosis airway communities predicts mechanisms of pathogen dominance. mSystems 4:e00026–19.

90. Aziz RK, Bartels D, Best AA, DeJongh M, Disz T, Edwards RA, Formsma K, Gerdes S, Glass EM, Kubal M. 2008. The RAST Server: rapid annotations using subsystems technology. BMC Genomics 9:75.

91. Monk JM, Lloyd CJ, Brunk E, Mih N, Sastry A, King Z, Takeuchi R, Nomura W, Zhang Z, Mori H. 2017. iML1515, a knowledgebase that computes Escherichia coli traits. Nature Biotech 35:904.

92. Teusink B, van Enckevort FH, Francke C, Wiersma A, Wegkamp A, Smid EJ, Siezen RJ. 2005. In silico reconstruction of the metabolic pathways of Lactobacillus plantarum: comparing predictions of nutrient requirements with those from growth experiments. Appl Env Microbiol 71:7253–7262.

93. Henry CS, DeJongh M, Best AA, Frybarger PM, Linsay B, Stevens RL. 2010. High-throughput generation, optimization and analysis of genome-scale metabolic models. Nature Biotech 28:977–982.

94. Caspi R, Foerster H, Fulcher CA, Kaipa P, Krummenacker M, Latendresse M, Paley S, Rhee SY, Shearer AG, Tissier C. 2008. The MetaCyc Database of metabolic pathways and enzymes and the BioCyc collection of Pathway/Genome Databases. Nuc Acids Res 36:D623–D631.

95. Kanehisa M, Goto S. 2000. KEGG: kyoto encyclopedia of genes and genomes. Nuc Acids Res 28:27–30.

96. Keseler IM, Mackie A, Peralta-Gil M, Santos-Zavaleta A, Gama-Castro S, Bonavides-Martínez C, Fulcher C, Huerta AM, Kothari A, Krummenacker M. 2013. EcoCyc: fusing model organism databases with systems biology. Nuc Acids Res 41:D605–D612.

97. Schellenberger J, Park JO, Conrad TM, Palsson BØ. 2010. BiGG: a Biochemical Genetic and Genomic knowledgebase of large scale metabolic reconstructions. BMC Bioinformatics 11:213.

98. Schomburg I, Chang A, Ebeling C, Gremse M, Heldt C, Huhn G, Schomburg D. 2004. BRENDA, the enzyme database: updates and major new developments. Nuc Acids Res 32:D431–D433.

99. Lieven C, Beber ME, Olivier BG, Bergmann FT, Ataman M, Babaei P, Bartell JA, Blank LM, Chauhan S, Correia K. 2020. MEMOTE for standardized genome-scale metabolic model testing. Nat Biotechnol 38:272–276.

100. McMullen JG. 2020. Effect of among-microbe interactions on Drosophila melanogaster nutrition. PhD Thesis. Cornell University.

101. Heirendt L, Arreckx S, Pfau T, Mendoza SN, Richelle A, Heinken A, Haraldsdottir HS, Wachowiak J, Keating SM, Vlasov V. 2019. Creation and analysis of biochemical constraint-based models using the COBRA Toolbox v. 3.0. Nature Protoc 14:639.

102. Orth JD, Thiele I, Palsson BØ. 2010. What is flux balance analysis? Nature Biotech 28:245–248.

103. R Core Team. 2019. R: A Language and Environment for Statistical Computing, R Foundation for Statistical Computing, Vienna, Austria. https://www.R-project.org.

104. Bates D, Maechler M, Bolker B, Walker S. 2015. Fitting Linear Mixed-Effects Models Using lme4. JStatist Software 67:1–48.

105. Fox J, Weisberg S. 2019. An {R} Companion to Applied Regression, Third ed. Sage, Thousand Oaks CA.

106. Bartoń K. 2019. MuMIn: Multi-Model Inference, vR package version 1.43.6. https://CRAN.R-project.org/package=MuMIn

107. Oksanen J, Blanchet FG, Friendly M, Kindt R, Legendre P, McGlinn D, Minchin PR, O’Hara RB, Simpson GL, Solymos P, Henry M, Stevens H, Szoecs E, Wagner H. 2019. vegan: Community Ecology Package, vR package version 2.5-5. https://CRAN.R-project.org/package=vegan.

108. Gurobi Optimization L. 2019. Gurobi Optimizer Reference Manual. http://www.gurobi.com.

109. Chelliah V, Juty N, Ajmera I, Ali R, Dumousseau M, Glont M, Hucka M, Jalowicki G, Keating S, Knight-Schrijver V. 2015. BioModels: ten-year anniversary. Nucleic Acids Res 43:D542–D548.

