## Supplementary Tables for "The Predicted Metabolic Function of the Gut Microbiota of *Drosophila melanogaster*"

**Table S1A. Predicted changes to bacterial growth (gdw $h^{-1}$) in co-culture compared to growth in monoculture - rich medium.**

| Community size | Community composition | Interaction type | Bacterium 1 | Change in growth [a] | Bacterium 2 | Change in growth [a] |
|---|---|---|---|---|---|---|
| 2 | AF-AP | Competitive | AF | -0.047543062 | AP | -0.065061688 |
| | AF-AT | Competitive | AF | -0.076238667 | AT | -0.036366088 |
| | AF-LB | Parasitic | AF | 0.055074439 | LB | -0.119156081 |
| | AF-LP | Parasitic | AF | 0.047204914 | LP | -0.1135639 |
| | AP-AT | Competitive | AP | -0.074874192 | AT | -0.037730678 |
| | AP-LB | Parasitic | AP | 0.059915684 | LB | -0.122596289 |
| | AP-LP | Parasitic | AP | 0.049206063 | LP | -0.114985903 |
| | AT-LB | Competitive | AT | -0.014359178 | LB | -0.069815219 |
| | AT-LP | Competitive | AT | -0.107234294 | LP | -0.003816429 |
| | LB-LP | Parasitic | LB | 0.032214901 | LP | -0.232262772 |
| 3 | AF-AP-AT | Competitive | AF | -0.087082938 | AP | -0.086340684 |
| | | Competitive | AF | -0.087082938 | AT | -0.051786046 |
| | | Competitive | AP | -0.086340684 | AT | -0.051786046 |
| | AF-AP-LB | Competitive | AF | -0.002853789 | AP | -0.048262407 |
| | | Competitive | AF | -0.002853789 | LB | -0.123714149 |
| | | Competitive | AP | -0.048262407 | LB | -0.123714149 |
| | AF-AP-LP | Competitive | AF | -0.029562466 | AP | -0.033782893 |
| | | Competitive | AF | -0.029562466 | LP | -0.115023822 |
| | | Competitive | AP | -0.033782893 | LP | -0.115023822 |
| | AF-AT-LB | Parasitic | AF | -0.110219045 | AT | 0.152376904 |
| | | Competitive | AF | -0.110219045 | LB | -0.189996451 |
| | | Parasitic | AT | 0.152376904 | LB | -0.189996451 |
| | AF-AT-LP | Parasitic | AF | -0.107561003 | AT | 0.131553873 |
| | | Competitive | AF | -0.107561003 | LP | -0.177088036 |
| | | Parasitic | AT | 0.131553873 | LP | -0.177088036 |
| | AF-LB-LP | Parasitic | AF | 0.06660369 | LB | -0.221068991 |
| | | Parasitic | AF | 0.06660369 | LP | -0.106327846 |
| | | Competitive | LB | -0.221068991 | LP | -0.106327846 |
| | AP-AT-LB | Parasitic | AP | -0.106949154 | AT | 0.150741462 |
| | | Competitive | AP | -0.106949154 | LB | -0.191157911 |
| | | Parasitic | AT | 0.150741462 | LB | -0.191157911 |
| | AP-AT-LP | Parasitic | AP | -0.074220679 | AT | 0.102520301 |
| | | Competitive | AP | -0.074220679 | LP | -0.180148566 |
| | | Parasitic | AT | 0.102520301 | LP | -0.180148566 |
| | AP-LB-LP | Parasitic | AP | 0.073164849 | LB | -0.216806519 |
| | | Parasitic | AP | 0.073164849 | LP | -0.115252826 |
| | | Competitive | LB | -0.216806519 | LP | -0.115252826 |
| | AT-LB-LP | Competitive | AT | -0.028038695 | LB | -0.156802785 |
| | | Competitive | AT | -0.028038695 | LP | -0.103339428 |
| | | Competitive | LB | -0.156802785 | LP | -0.103339428 |
| 4 | AF-AP-AT-LB | Competitive | AF | -0.110695191 | AP | -0.108231736 |
| | | Parasitic | AF | -0.110695191 | AT | 0.150034022 |
| | | Competitive | AF | -0.110695191 | LB | -0.191100792 |
| | | Parasitic | AP | -0.108231736 | AT | 0.150034022 |
| | | Competitive | AP | -0.108231736 | LB | -0.191100792 |
| | | Parasitic | AT | 0.150034022 | LB | -0.191100792 |
| | AF-AP-AT-LP | Competitive | AF | -0.106694143 | AP | -0.077213775 |
| | | Parasitic | AF | -0.106694143 | AT | 0.099597843 |
| | | Competitive | AF | -0.106694143 | LP | -0.180144987 |
| | | Parasitic | AP | -0.077213775 | AT | 0.099597843 |
| | | Competitive | AP | -0.077213775 | LP | -0.180144987 |
| | | Parasitic | AT | 0.099597843 | LP | -0.180144987 |
| | AF-AP-LB-LP | Parasitic | AF | 0.011016456 | AP | -0.050282481 |
| | | Parasitic | AF | 0.011016456 | LB | -0.218304048 |
| | | Parasitic | AF | 0.011016456 | LP | -0.113878797 |

| | | | | | | |
|---|---|---|---|---|---|---|
| | | Competitive | AP | -0.050282481 | LB | -0.218304048 |
| | | Competitive | AP | -0.050282481 | LP | -0.113878797 |
| | | Competitive | LB | -0.218304048 | LP | -0.113878797 |
| | AF-AT-LB-LP | Parasitic | AF | -0.109146218 | AT | 0.166705235 |
| | | Competitive | AF | -0.109146218 | LB | -0.213566522 |
| | | Competitive | AF | -0.109146218 | LP | -0.187422146 |
| | | Parasitic | AT | 0.166705235 | LB | -0.213566522 |
| | | Parasitic | AT | 0.166705235 | LP | -0.187422146 |
| | | Competitive | LB | -0.213566522 | LP | -0.187422146 |
| | AP-AT-LB-LP | Parasitic | AP | -0.106756685 | AT | 0.166029534 |
| | | Competitive | AP | -0.106756685 | LB | -0.214101504 |
| | | Competitive | AP | -0.106756685 | LP | -0.188105092 |
| | | Parasitic | AT | 0.166029534 | LB | -0.214101504 |
| | | Parasitic | AT | 0.166029534 | LP | -0.188105092 |
| | | Competitive | LB | -0.214101504 | LP | -0.188105092 |
| 5 | AF-AP-AT-LB-LP | Competitive | AF | -0.108186463 | AP | -0.107792338 |
| | | Parasitic | AF | -0.108186463 | AT | 0.162643559 |
| | | Competitive | AF | -0.108186463 | LB | -0.215194119 |
| | | Competitive | AF | -0.108186463 | LP | -0.187010075 |
| | | Parasitic | AP | -0.107792338 | AT | 0.162643559 |
| | | Competitive | AP | -0.107792338 | LB | -0.215194119 |
| | | Competitive | AP | -0.107792338 | LP | -0.187010075 |
| | | Parasitic | AT | 0.162643559 | LB | -0.215194119 |
| | | Parasitic | AT | 0.162643559 | LP | -0.187010075 |
| | | Competitive | LB | -0.215194119 | LP | -0.187010075 |

AF-*Acetobacter fabarum*; AP-*Acetobacter pomorum*; AT-*Acetobacter tropicalis*; LB-*Lactobacillus brevis*; LP-*Lactobacillus plantarum*

[a]Change in growth is calculated by subtracting growth of a microbe in mono-culture to growth of the microbe in co-culture.

**Table S1B. Predicted changes to bacterial growth (gdw $h^{-1}$) in co-culture compared to growth in isolation - base medium.**

| Community size | Community composition | Interaction type | Bacterium 1 | Change in growth [a] | Bacterium 2 | Change in growth [a] |
|---|---|---|---|---|---|---|
| 2 | AF-AP | Competitive | AF | -0.001884309 | AP | -0.009319108 |
| | AF-AT | Competitive | AF | -0.009198881 | AT | -0.002004535 |
| | AF-LB | Competitive | AF | -0.007072457 | LB | -0.002948602 |
| | AF-LP | Parasitic | AF | 0.000173024 | LP | -0.008120277 |
| | AP-AT | Competitive | AP | -0.008869707 | AT | -0.002333709 |
| | AP-LB | Competitive | AP | -0.007072456 | LB | -0.002948602 |
| | AP-LP | Parasitic | AP | 0.000247659 | LP | -0.008173545 |
| | AT-LB | Competitive | AT | -0.009023334 | LB | -0.001556102 |
| | AT-LP | Parasitic | AT | 0.001117801 | LP | -0.008794637 |
| | LB-LP | Competitive | LB | -0.001573434 | LP | -0.014420165 |
| 3 | AF-AP-AT | Competitive | AF | -0.009702916 | AP | -0.009721573 |
| | | Competitive | AF | -0.009702916 | AT | -0.002982338 |
| | | Competitive | AP | -0.009721573 | AT | -0.002982338 |
| | AF-AP-LB | Competitive | AF | -0.004808531 | AP | -0.010774966 |
| | | Competitive | AF | -0.004808531 | LB | -0.004870388 |
| | | Competitive | AP | -0.010774966 | LB | -0.004870388 |
| | AF-AP-LP | Competitive | AF | -0.005672564 | AP | -0.005215904 |
| | | Competitive | AF | -0.005672564 | LP | -0.008221575 |
| | | Competitive | AP | -0.005215904 | LP | -0.008221575 |
| | AF-AT-LB | Parasitic | AF | -0.010831436 | AT | 0.014005427 |
| | | Competitive | AF | -0.010831436 | LB | -0.018259108 |
| | | Parasitic | AT | 0.014005427 | LB | -0.018259108 |
| | AF-AT-LP | Parasitic | AF | -0.010836787 | AT | 0.006768986 |
| | | Competitive | AF | -0.010836787 | LP | -0.013090051 |
| | | Parasitic | AT | 0.006768986 | LP | -0.013090051 |
| | AF-LB-LP | Parasitic | AF | 0.00453964 | LB | -0.016130989 |

| | | | | | | |
|---|---|---|---|---|---|---|
| | | Parasitic | AF | 0.00453964 | LP | -0.011099678 |
| | | Competitive | LB | -0.016130989 | LP | -0.011099678 |
| | AP-AT-LB | Parasitic | AP | -0.010833167 | AT | 0.014395807 |
| | | Competitive | AP | -0.010833167 | LB | -0.018536512 |
| | | Parasitic | AT | 0.014395807 | LB | -0.018536512 |
| | AP-AT-LP | Parasitic | AP | -0.010835518 | AT | 0.007082089 |
| | | Competitive | AP | -0.010835518 | LP | -0.013314428 |
| | | Parasitic | AT | 0.007082089 | LP | -0.013314428 |
| | AP-LB-LP | Parasitic | AP | 0.004762498 | LB | -0.015717421 |
| | | Parasitic | AP | 0.004762498 | LP | -0.01167233 |
| | | Competitive | LB | -0.015717421 | LP | -0.01167233 |
| | AT-LB-LP | Competitive | AT | -0.005040301 | LB | -0.017196784 |
| | | Competitive | AT | -0.005040301 | LP | -0.003195905 |
| | | Competitive | LB | -0.017196784 | LP | -0.003195905 |
| 4 | AF-AP-AT-LB | Competitive | AF | -0.010834665 | AP | -0.010850631 |
| | | Parasitic | AF | -0.010834665 | AT | 0.013922938 |
| | | Competitive | AF | -0.010834665 | LB | -0.018449745 |
| | | Parasitic | AP | -0.010850631 | AT | 0.013922938 |
| | | Competitive | AP | -0.010850631 | LB | -0.018449745 |
| | | Parasitic | AT | 0.013922938 | LB | -0.018449745 |
| | AF-AP-AT-LP | Competitive | AF | -0.010850423 | AP | -0.010839312 |
| | | Parasitic | AF | -0.010850423 | AT | 0.006582039 |
| | | Competitive | AF | -0.010850423 | LP | -0.013206757 |
| | | Parasitic | AP | -0.010839312 | AT | 0.006582039 |
| | | Competitive | AP | -0.010839312 | LP | -0.013206757 |
| | | Parasitic | AT | 0.006582039 | LP | -0.013206757 |
| | AF-AP-LB-LP | Competitive | AF | -0.003309859 | AP | -0.003135658 |
| | | Competitive | AF | -0.003309859 | LB | -0.01575855 |
| | | Competitive | AF | -0.003309859 | LP | -0.01162791 |
| | | Competitive | AP | -0.003135658 | LB | -0.01575855 |
| | | Competitive | AP | -0.003135658 | LP | -0.01162791 |
| | | Competitive | LB | -0.01575855 | LP | -0.01162791 |
| | AF-AT-LB-LP | Parasitic | AF | -0.010848141 | AT | 0.013942745 |
| | | Competitive | AF | -0.010848141 | LB | -0.018583218 |
| | | Competitive | AF | -0.010848141 | LP | -0.01561282 |
| | | Parasitic | AT | 0.013942745 | LB | -0.018583218 |
| | | Parasitic | AT | 0.013942745 | LP | -0.01561282 |
| | | Competitive | LB | -0.018583218 | LP | -0.01561282 |
| | AP-AT-LB-LP | Parasitic | AP | -0.010838311 | AT | 0.014321286 |
| | | Competitive | AP | -0.010838311 | LB | -0.0188563 |
| | | Competitive | AP | -0.010838311 | LP | -0.01561693 |
| | | Parasitic | AT | 0.014321286 | LB | -0.0188563 |
| | | Parasitic | AT | 0.014321286 | LP | -0.01561693 |
| | | Competitive | LB | -0.0188563 | LP | -0.01561693 |
| 5 | AF-AP-AT-LB-LP | Competitive | AF | -0.010850631 | AP | -0.01083872 |
| | | Parasitic | AF | -0.010850631 | AT | 0.013847162 |
| | | Competitive | AF | -0.010850631 | LB | -0.018770392 |
| | | Competitive | AF | -0.010850631 | LP | -0.015615938 |
| | | Parasitic | AP | -0.01083872 | AT | 0.013847162 |
| | | Competitive | AP | -0.01083872 | LB | -0.018770392 |
| | | Competitive | AP | -0.01083872 | LP | -0.015615938 |
| | | Parasitic | AT | 0.013847162 | LB | -0.018770392 |
| | | Parasitic | AT | 0.013847162 | LP | -0.015615938 |
| | | Competitive | LB | -0.018770392 | LP | -0.015615938 |

AF-*Acetobacter fabarum*; AP-*Acetobacter pomorum*; AT-*Acetobacter tropicalis*; LB-*Lactobacillus brevis*; LP-*Lactobacillus plantarum*

[a] Change in growth is calculated by subtracting growth of a microbe in mono-culture to growth of the microbe in co-culture.

**Table S1C. Predicted changes to bacterial growth (gdw h$^{-1}$) in co-culture compared to growth in isolation - minimal medium.**

| Community size | Community composition | Interaction type | Bacterium 1 | Change in growth [a] | Bacterium 2 | Change in growth [a] |
|---|---|---|---|---|---|---|
| 2 | AF-AP | Parasitic | AF | 0.004306849 | AP | -0.004306849 |
| | AF-AT | Parasitic | AF | 0.000426858 | AT | -0.000426858 |
| | AF-LB | Neutral | AF | 0 | LB | 0 |
| | AF-LP | Mutualistic | AF | 0.003271789 | LP | 0.002523747 |
| | AP-AT | Competitive | AP | -0.004216446 | AT | -0.000438274 |
| | AP-LB | Amensal | AP | -0.00465472 | LB | 0 |
| | AP-LP | Parasitic | AP | -0.001375683 | LP | 0.00251052 |
| | AT-LB | Amensal | AT | -0.00465472 | LB | 0 |
| | AT-LP | Parasitic | AT | -0.001614864 | LP | 0.002947009 |
| | LB-LP | Neutral | LB | 0 | LP | 0 |
| 3 | AF-AP-AT | Parasitic | AF | 0.000404493 | AP | -0.004250006 |
| | | Parasitic | AF | 0.000404493 | AT | -0.000809207 |
| | | Competitive | AP | -0.004250006 | AT | -0.000809207 |
| | AF-AP-LB | Amensal | AF | 0 | AP | -0.00465472 |
| | | Neutral | AF | 0 | LB | 0 |
| | | Amensal | AP | -0.00465472 | LB | 0 |
| | AF-AP-LP | Parasitic | AF | 0.002786746 | AP | -0.004291046 |
| | | Mutual | AF | 0.002786746 | LP | 0.002745237 |
| | | Parasitic | AP | -0.004291046 | LP | 0.002745237 |
| | AF-AT-LB | Amensal | AF | 0 | AT | -0.00465472 |
| | | Neutral | AF | 0 | LB | 0 |
| | | Amensal | AT | -0.00465472 | LB | 0 |
| | AF-AT-LP | Parasitic | AF | 0.000355333 | AT | -0.00429841 |
| | | Mutual | AF | 0.000355333 | LP | 0.007195837 |
| | | Parasitic | AT | -0.00429841 | LP | 0.007195837 |
| | AF-LB-LP | Mutual | AF | 0.003216672 | LB | 0.000451943 |
| | | Mutual | AF | 0.003216672 | LP | 0.002168724 |
| | | Mutual | LB | 0.000451943 | LP | 0.002168724 |
| | AP-AT-LB | Competitive | AP | -0.00465472 | AT | -0.00465472 |
| | | Amensal | AP | -0.00465472 | LB | 0 |
| | | Amensal | AT | -0.00465472 | LB | 0 |
| | AP-AT-LP | Competitive | AP | -0.004301871 | AT | -0.004301009 |
| | | Parasitic | AP | -0.004301871 | LP | 0.007205113 |
| | | Parasitic | AT | -0.004301009 | LP | 0.007205113 |
| | AP-LB-LP | Parasitic | AP | -0.001419561 | LB | 0.000449321 |
| | | Parasitic | AP | -0.001419561 | LP | 0.002137628 |
| | | Mutual | LB | 0.000449321 | LP | 0.002137628 |
| | AT-LB-LP | Parasitic | AT | -0.003674414 | LB | 0.003541452 |
| | | Parasitic | AT | -0.003674414 | LP | 0.003135363 |
| | | Mutual | LB | 0.003541452 | LP | 0.003135363 |
| 4 | AF-AP-AT-LB | Amensal | AF | 0 | AP | -0.00465472 |
| | | Amensal | AF | 0 | AT | -0.00465472 |
| | | Neutral | AF | 0 | LB | 0 |
| | | Competitive | AP | -0.00465472 | AT | -0.00465472 |
| | | Amensal | AP | -0.00465472 | LB | 0 |
| | | Amensal | AT | -0.00465472 | LB | 0 |
| | AF-AP-AT-LP | Parasitic | AF | 0.000352853 | AP | -0.004301867 |
| | | Parasitic | AF | 0.000352853 | AT | -0.004301188 |
| | | Mutual | AF | 0.000352853 | LP | 0.006561499 |
| | | Competitive | AP | -0.004301867 | AT | -0.004301188 |
| | | Parasitic | AP | -0.004301867 | LP | 0.006561499 |
| | | Parasitic | AT | -0.004301188 | LP | 0.006561499 |
| | AF-AP-LB-LP | Parasitic | AF | 0.002791665 | AP | -0.004269056 |
| | | Mutual | AF | 0.002791665 | LB | 0.000377067 |

| | | | | | | |
|---|---|---|---|---|---|---|
| | | Mutual | AF | 0.002791665 | LP | 0.002316005 |
| | | Parasitic | AP | -0.004269056 | LB | 0.000377067 |
| | | Parasitic | AP | -0.004269056 | LP | 0.002316005 |
| | | Mutual | LB | 0.000377067 | LP | 0.002316005 |
| | AF-AT-LB-LP | Parasitic | AF | 0.000355324 | AT | -0.004299397 |
| | | Mutual | AF | 0.000355324 | LB | 0.000481636 |
| | | Mutual | AF | 0.000355324 | LP | 0.006712112 |
| | | Parasitic | AT | -0.004299397 | LB | 0.000481636 |
| | | Parasitic | AT | -0.004299397 | LP | 0.006712112 |
| | | Mutual | LB | 0.000481636 | LP | 0.006712112 |
| | AP-AT-LB-LP | Competitive | AP | -0.004301471 | AT | -0.004301906 |
| | | Parasitic | AP | -0.004301471 | LB | 0.000478799 |
| | | Parasitic | AP | -0.004301471 | LP | 0.006723336 |
| | | Parasitic | AT | -0.004301906 | LB | 0.000478799 |
| | | Parasitic | AT | -0.004301906 | LP | 0.006723336 |
| | | Mutual | LB | 0.000478799 | LP | 0.006723336 |
| 5 | AF-AP-AT-LB-LP | Parasitic | AF | 0.000352834 | AP | -0.004301887 |
| | | Parasitic | AF | 0.000352834 | AT | -0.004301887 |
| | | Mutual | AF | 0.000352834 | LB | 0.000474397 |
| | | Mutual | AF | 0.000352834 | LP | 0.006084601 |
| | | Competitive | AP | -0.004301887 | AT | -0.004301887 |
| | | Parasitic | AP | -0.004301887 | LB | 0.000474397 |
| | | Parasitic | AP | -0.004301887 | LP | 0.006084601 |
| | | Parasitic | AT | -0.004301887 | LB | 0.000474397 |
| | | Parasitic | AT | -0.004301887 | LP | 0.006084601 |
| | | Mutual | LB | 0.000474397 | LP | 0.006084601 |

AF-*Acetobacter fabarum*; AP-*Acetobacter pomorum*; AT-*Acetobacter tropicalis*; LB-*Lactobacillus brevis*; LP-*Lactobacillus plantarum*

[a] Change in growth is calculated by subtracting growth of a microbe in mono-culture to growth of the microbe in co-culture.

**Table S2A. Predicted number of inputs and outputs from bacteria - rich medium.**

| Community | Number of species | Organism | Number of overlapping inputs | Number of overlapping outputs | Total input count | Total output count |
|---|---|---|---|---|---|---|
| AF | 1 | AF | 0 | 0 | 29 | 6 |
| AF_AP | 2 | AF | 24 | 6 | 28 | 6 |
| AF_AP | 2 | AP | 24 | 6 | 26 | 6 |
| AF_AP_AT | 3 | AF | 24 | 6 | 26 | 6 |
| AF_AP_AT | 3 | AP | 24 | 6 | 26 | 6 |
| AF_AP_AT | 3 | AT | 19 | 6 | 26 | 9 |
| AF_AP_AT_LB | 4 | AF | 21 | 5 | 24 | 6 |
| AF_AP_AT_LB | 4 | AP | 24 | 5 | 26 | 6 |
| AF_AP_AT_LB | 4 | AT | 24 | 4 | 36 | 7 |
| AF_AP_AT_LB | 4 | LB | 12 | 1 | 40 | 12 |
| AF_AP_AT_LB_LP | 5 | AF | 23 | 5 | 25 | 5 |
| AF_AP_AT_LB_LP | 5 | AP | 25 | 5 | 26 | 6 |
| AF_AP_AT_LB_LP | 5 | AT | 25 | 4 | 37 | 6 |
| AF_AP_AT_LB_LP | 5 | LB | 25 | 3 | 37 | 8 |
| AF_AP_AT_LB_LP | 5 | LP | 27 | 2 | 39 | 5 |
| AF_AP_AT_LP | 4 | AF | 24 | 5 | 25 | 5 |
| AF_AP_AT_LP | 4 | AP | 25 | 5 | 26 | 6 |
| AF_AP_AT_LP | 4 | AT | 22 | 4 | 32 | 8 |
| AF_AP_AT_LP | 4 | LP | 11 | 1 | 36 | 7 |
| AF_AP_LB | 3 | AF | 25 | 5 | 28 | 5 |
| AF_AP_LB | 3 | AP | 24 | 5 | 27 | 6 |
| AF_AP_LB | 3 | LB | 10 | 1 | 41 | 12 |
| AF_AP_LB_LP | 4 | AF | 25 | 5 | 28 | 5 |
| AF_AP_LB_LP | 4 | AP | 23 | 5 | 26 | 6 |
| AF_AP_LB_LP | 4 | LB | 25 | 4 | 36 | 10 |
| AF_AP_LB_LP | 4 | LP | 29 | 2 | 46 | 5 |
| AF_AP_LP | 3 | AF | 26 | 6 | 27 | 6 |
| AF_AP_LP | 3 | AP | 26 | 6 | 30 | 7 |
| AF_AP_LP | 3 | LP | 13 | 0 | 44 | 7 |
| AF_AT | 2 | AF | 19 | 6 | 26 | 6 |
| AF_AT | 2 | AT | 19 | 6 | 25 | 9 |
| AF_AT_LB | 3 | AF | 19 | 6 | 24 | 6 |
| AF_AT_LB | 3 | AT | 20 | 6 | 35 | 8 |
| AF_AT_LB | 3 | LB | 12 | 1 | 40 | 12 |
| AF_AT_LB_LP | 4 | AF | 21 | 4 | 25 | 5 |
| AF_AT_LB_LP | 4 | AT | 23 | 4 | 37 | 6 |
| AF_AT_LB_LP | 4 | LB | 25 | 3 | 37 | 8 |
| AF_AT_LB_LP | 4 | LP | 27 | 2 | 39 | 5 |
| AF_AT_LP | 3 | AF | 22 | 5 | 25 | 6 |
| AF_AT_LP | 3 | AT | 21 | 5 | 33 | 9 |
| AF_AT_LP | 3 | LP | 11 | 1 | 36 | 7 |
| AF_LB | 2 | AF | 10 | 1 | 30 | 6 |
| AF_LB | 2 | LB | 10 | 1 | 41 | 12 |
| AF_LB_LP | 3 | AF | 15 | 2 | 29 | 5 |
| AF_LB_LP | 3 | LB | 25 | 4 | 36 | 10 |
| AF_LB_LP | 3 | LP | 29 | 2 | 46 | 5 |
| AF_LP | 2 | AF | 11 | 0 | 30 | 7 |
| AF_LP | 2 | LP | 11 | 0 | 44 | 7 |
| AP | 1 | AP | 0 | 0 | 29 | 6 |
| AP_AT | 2 | AP | 19 | 6 | 27 | 6 |
| AP_AT | 2 | AT | 19 | 6 | 26 | 9 |
| AP_AT_LB | 3 | AP | 23 | 4 | 26 | 6 |
| AP_AT_LB | 3 | AT | 25 | 4 | 36 | 7 |
| AP_AT_LB | 3 | LB | 11 | 1 | 40 | 12 |

| | | | | | | |
|---|---|---|---|---|---|---|
| AP_AT_LB_LP | 4 | AP | 22 | 4 | 25 | 6 |
| AP_AT_LB_LP | 4 | AT | 25 | 4 | 37 | 6 |
| AP_AT_LB_LP | 4 | LB | 26 | 3 | 37 | 8 |
| AP_AT_LB_LP | 4 | LP | 28 | 2 | 40 | 5 |
| AP_AT_LP | 3 | AP | 22 | 4 | 26 | 6 |
| AP_AT_LP | 3 | AT | 23 | 4 | 33 | 8 |
| AP_AT_LP | 3 | LP | 11 | 1 | 36 | 7 |
| AP_LB | 2 | AP | 9 | 1 | 30 | 6 |
| AP_LB | 2 | LB | 9 | 1 | 41 | 12 |
| AP_LB_LP | 3 | AP | 12 | 2 | 28 | 6 |
| AP_LB_LP | 3 | LB | 25 | 4 | 36 | 10 |
| AP_LB_LP | 3 | LP | 28 | 2 | 45 | 5 |
| AP_LP | 2 | AP | 12 | 0 | 30 | 7 |
| AP_LP | 2 | LP | 12 | 0 | 45 | 7 |
| AT | 1 | AT | 0 | 0 | 26 | 8 |
| AT_LB | 2 | AT | 9 | 1 | 26 | 7 |
| AT_LB | 2 | LB | 9 | 1 | 41 | 10 |
| AT_LB_LP | 3 | AT | 12 | 1 | 29 | 7 |
| AT_LB_LP | 3 | LB | 27 | 3 | 38 | 8 |
| AT_LB_LP | 3 | LP | 28 | 2 | 42 | 4 |
| AT_LP | 2 | AT | 13 | 1 | 28 | 7 |
| AT_LP | 2 | LP | 13 | 1 | 44 | 6 |
| LB | 1 | LB | 0 | 0 | 45 | 9 |
| LB_LP | 2 | LB | 15 | 4 | 45 | 9 |
| LB_LP | 2 | LP | 15 | 4 | 30 | 6 |
| LP | 1 | LP | 0 | 0 | 44 | 6 |

AF-*Acetobacter fabarum*; AP-*Acetobacter pomorum*; AT-*Acetobacter tropicalis*; LB-*Lactobacillus brevis*; LP-*Lactobacillus plantarum*

**Table S2B. Predicted number of inputs and outputs from bacteria - base medium.**

| Community | Number of species | Organism | Number of overlapping inputs | Number of overlapping outputs | Total input count | Total output count |
|---|---|---|---|---|---|---|
| AF | 1 | AF | 0 | 0 | 15 | 6 |
| AF_AP | 2 | AF | 13 | 6 | 14 | 6 |
| AF_AP | 2 | AP | 13 | 6 | 14 | 6 |
| AF_AP_AT | 3 | AF | 15 | 4 | 20 | 5 |
| AF_AP_AT | 3 | AP | 15 | 4 | 21 | 5 |
| AF_AP_AT | 3 | AT | 12 | 4 | 15 | 12 |
| AF_AP_AT_LB | 4 | AF | 17 | 4 | 22 | 5 |
| AF_AP_AT_LB | 4 | AP | 18 | 4 | 22 | 5 |
| AF_AP_AT_LB | 4 | AT | 18 | 4 | 22 | 9 |
| AF_AP_AT_LB | 4 | LB | 10 | 1 | 25 | 13 |
| AF_AP_AT_LB_LP | 5 | AF | 17 | 4 | 22 | 5 |
| AF_AP_AT_LB_LP | 5 | AP | 18 | 4 | 21 | 6 |
| AF_AP_AT_LB_LP | 5 | AT | 18 | 5 | 22 | 9 |
| AF_AP_AT_LB_LP | 5 | LB | 19 | 5 | 25 | 13 |
| AF_AP_AT_LB_LP | 5 | LP | 21 | 6 | 25 | 7 |
| AF_AP_AT_LP | 4 | AF | 16 | 4 | 19 | 5 |
| AF_AP_AT_LP | 4 | AP | 16 | 4 | 19 | 5 |
| AF_AP_AT_LP | 4 | AT | 15 | 5 | 18 | 9 |
| AF_AP_AT_LP | 4 | LP | 12 | 1 | 24 | 7 |
| AF_AP_LB | 3 | AF | 19 | 6 | 21 | 7 |
| AF_AP_LB | 3 | AP | 18 | 5 | 21 | 6 |
| AF_AP_LB | 3 | LB | 11 | 2 | 26 | 11 |
| AF_AP_LB_LP | 4 | AF | 20 | 7 | 22 | 7 |
| AF_AP_LB_LP | 4 | AP | 20 | 7 | 21 | 7 |
| AF_AP_LB_LP | 4 | LB | 17 | 2 | 25 | 13 |

| | | | | | | |
|---|---|---|---|---|---|---|
| AF_AP_LB_LP | 4 | LP | 18 | 1 | 28 | 5 |
| AF_AP_LP | 3 | AF | 18 | 7 | 19 | 7 |
| AF_AP_LP | 3 | AP | 19 | 7 | 20 | 8 |
| AF_AP_LP | 3 | LP | 12 | 0 | 25 | 6 |
| AF_AT | 2 | AF | 12 | 4 | 19 | 5 |
| AF_AT | 2 | AT | 12 | 4 | 15 | 11 |
| AF_AT_LB | 3 | AF | 16 | 4 | 22 | 5 |
| AF_AT_LB | 3 | AT | 17 | 4 | 22 | 9 |
| AF_AT_LB | 3 | LB | 10 | 1 | 25 | 13 |
| AF_AT_LB_LP | 4 | AF | 16 | 4 | 22 | 5 |
| AF_AT_LB_LP | 4 | AT | 18 | 5 | 22 | 9 |
| AF_AT_LB_LP | 4 | LB | 19 | 5 | 25 | 12 |
| AF_AT_LB_LP | 4 | LP | 21 | 6 | 26 | 8 |
| AF_AT_LP | 3 | AF | 16 | 4 | 20 | 5 |
| AF_AT_LP | 3 | AT | 15 | 5 | 18 | 9 |
| AF_AT_LP | 3 | LP | 12 | 1 | 24 | 7 |
| AF_LB | 2 | AF | 12 | 2 | 21 | 7 |
| AF_LB | 2 | LB | 12 | 2 | 27 | 10 |
| AF_LB_LP | 3 | AF | 12 | 3 | 23 | 7 |
| AF_LB_LP | 3 | LB | 17 | 2 | 25 | 13 |
| AF_LB_LP | 3 | LP | 18 | 1 | 28 | 5 |
| AF_LP | 2 | AF | 12 | 0 | 20 | 8 |
| AF_LP | 2 | LP | 12 | 0 | 25 | 6 |
| AP | 1 | AP | 0 | 0 | 14 | 7 |
| AP_AT | 2 | AP | 12 | 5 | 21 | 5 |
| AP_AT | 2 | AT | 12 | 5 | 15 | 12 |
| AP_AT_LB | 3 | AP | 16 | 4 | 21 | 5 |
| AP_AT_LB | 3 | AT | 17 | 4 | 22 | 8 |
| AP_AT_LB | 3 | LB | 10 | 1 | 25 | 13 |
| AP_AT_LB_LP | 4 | AP | 16 | 4 | 20 | 5 |
| AP_AT_LB_LP | 4 | AT | 18 | 5 | 22 | 7 |
| AP_AT_LB_LP | 4 | LB | 19 | 5 | 25 | 13 |
| AP_AT_LB_LP | 4 | LP | 21 | 6 | 26 | 7 |
| AP_AT_LP | 3 | AP | 16 | 4 | 20 | 5 |
| AP_AT_LP | 3 | AT | 15 | 5 | 18 | 9 |
| AP_AT_LP | 3 | LP | 12 | 1 | 24 | 7 |
| AP_LB | 2 | AP | 12 | 2 | 21 | 7 |
| AP_LB | 2 | LB | 12 | 2 | 27 | 10 |
| AP_LB_LP | 3 | AP | 12 | 2 | 22 | 7 |
| AP_LB_LP | 3 | LB | 17 | 1 | 25 | 12 |
| AP_LB_LP | 3 | LP | 18 | 1 | 28 | 5 |
| AP_LP | 2 | AP | 12 | 0 | 20 | 8 |
| AP_LP | 2 | LP | 12 | 0 | 25 | 6 |
| AT | 1 | AT | 0 | 0 | 16 | 6 |
| AT_LB | 2 | AT | 11 | 1 | 20 | 6 |
| AT_LB | 2 | LB | 11 | 1 | 27 | 9 |
| AT_LB_LP | 3 | AT | 14 | 1 | 20 | 6 |
| AT_LB_LP | 3 | LB | 17 | 1 | 25 | 12 |
| AT_LB_LP | 3 | LP | 20 | 0 | 31 | 3 |
| AT_LP | 2 | AT | 12 | 0 | 19 | 7 |
| AT_LP | 2 | LP | 12 | 0 | 24 | 5 |
| LB | 1 | LB | 0 | 0 | 27 | 4 |
| LB_LP | 2 | LB | 18 | 0 | 27 | 10 |
| LB_LP | 2 | LP | 18 | 0 | 30 | 2 |
| LP | 1 | LP | 0 | 0 | 22 | 4 |

AF-*Acetobacter fabarum*; AP-*Acetobacter pomorum*; AT-*Acetobacter tropicalis*; LB-*Lactobacillus brevis*; LP-*Lactobacillus plantarum*

**Table S2C. Predicted number of inputs and outputs from bacteria - minimal medium.**

| Community | Number of species | Organism | Number of overlapping inputs | Number of overlapping outputs | Total input count | Total output count |
|---|---|---|---|---|---|---|
| AF | 1 | AF | 0 | 0 | 0 | 0 |
| AF_AP | 2 | AF | 4 | 1 | 9 | 7 |
| AF_AP | 2 | AP | 4 | 1 | 11 | 5 |
| AF_AP_AT | 3 | AF | 8 | 1 | 18 | 1 |
| AF_AP_AT | 3 | AP | 8 | 1 | 16 | 2 |
| AF_AP_AT | 3 | AT | 3 | 1 | 5 | 17 |
| AF_AP_AT_LB | 4 | AF | 0 | 0 | 0 | 0 |
| AF_AP_AT_LB | 4 | AP | 0 | 0 | 0 | 0 |
| AF_AP_AT_LB | 4 | AT | 0 | 0 | 0 | 0 |
| AF_AP_AT_LB | 4 | LB | 0 | 0 | 0 | 0 |
| AF_AP_AT_LB_LP | 5 | AF | 12 | 5 | 14 | 10 |
| AF_AP_AT_LB_LP | 5 | AP | 12 | 5 | 12 | 13 |
| AF_AP_AT_LB_LP | 5 | AT | 11 | 4 | 12 | 10 |
| AF_AP_AT_LB_LP | 5 | LB | 11 | 1 | 28 | 6 |
| AF_AP_AT_LB_LP | 5 | LP | 8 | 1 | 18 | 18 |
| AF_AP_AT_LP | 4 | AF | 11 | 4 | 14 | 7 |
| AF_AP_AT_LP | 4 | AP | 11 | 4 | 13 | 8 |
| AF_AP_AT_LP | 4 | AT | 10 | 5 | 11 | 11 |
| AF_AP_AT_LP | 4 | LP | 4 | 1 | 16 | 10 |
| AF_AP_LB | 3 | AF | 0 | 0 | 0 | 0 |
| AF_AP_LB | 3 | AP | 0 | 0 | 0 | 0 |
| AF_AP_LB | 3 | LB | 0 | 0 | 0 | 0 |
| AF_AP_LB_LP | 4 | AF | 11 | 2 | 12 | 10 |
| AF_AP_LB_LP | 4 | AP | 11 | 3 | 12 | 11 |
| AF_AP_LB_LP | 4 | LB | 9 | 1 | 27 | 5 |
| AF_AP_LB_LP | 4 | LP | 9 | 1 | 15 | 18 |
| AF_AP_LP | 3 | AF | 10 | 2 | 15 | 8 |
| AF_AP_LP | 3 | AP | 11 | 3 | 12 | 10 |
| AF_AP_LP | 3 | LP | 4 | 1 | 14 | 10 |
| AF_AT | 2 | AF | 3 | 1 | 17 | 1 |
| AF_AT | 2 | AT | 3 | 1 | 4 | 14 |
| AF_AT_LB | 3 | AF | 0 | 0 | 0 | 0 |
| AF_AT_LB | 3 | AT | 0 | 0 | 0 | 0 |
| AF_AT_LB | 3 | LB | 0 | 0 | 0 | 0 |
| AF_AT_LB_LP | 4 | AF | 11 | 7 | 15 | 9 |
| AF_AT_LB_LP | 4 | AT | 10 | 8 | 11 | 20 |
| AF_AT_LB_LP | 4 | LB | 12 | 1 | 27 | 7 |
| AF_AT_LB_LP | 4 | LP | 8 | 1 | 19 | 17 |
| AF_AT_LP | 3 | AF | 10 | 6 | 14 | 9 |
| AF_AT_LP | 3 | AT | 10 | 6 | 12 | 17 |
| AF_AT_LP | 3 | LP | 3 | 0 | 17 | 9 |
| AF_LB | 2 | AF | 0 | 0 | 0 | 0 |
| AF_LB | 2 | LB | 0 | 0 | 0 | 0 |
| AF_LB_LP | 3 | AF | 9 | 3 | 14 | 19 |
| AF_LB_LP | 3 | LB | 10 | 1 | 27 | 4 |
| AF_LB_LP | 3 | LP | 9 | 2 | 15 | 19 |
| AF_LP | 2 | AF | 3 | 0 | 13 | 13 |
| AF_LP | 2 | LP | 3 | 0 | 14 | 10 |
| AP | 1 | AP | 0 | 0 | 5 | 2 |
| AP_AT | 2 | AP | 3 | 1 | 16 | 2 |
| AP_AT | 2 | AT | 3 | 1 | 5 | 13 |
| AP_AT_LB | 3 | AP | 0 | 0 | 0 | 0 |
| AP_AT_LB | 3 | AT | 0 | 0 | 0 | 0 |
| AP_AT_LB | 3 | LB | 0 | 0 | 0 | 0 |

| | | | | | | |
|---|---|---|---|---|---|---|
| AP_AT_LB_LP | 4 | AP | 12 | 4 | 12 | 11 |
| AP_AT_LB_LP | 4 | AT | 11 | 4 | 12 | 13 |
| AP_AT_LB_LP | 4 | LB | 10 | 1 | 28 | 6 |
| AP_AT_LB_LP | 4 | LP | 8 | 1 | 17 | 18 |
| AP_AT_LP | 3 | AP | 11 | 6 | 13 | 9 |
| AP_AT_LP | 3 | AT | 10 | 5 | 12 | 14 |
| AP_AT_LP | 3 | LP | 4 | 1 | 15 | 10 |
| AP_LB | 2 | AP | 0 | 0 | 0 | 0 |
| AP_LB | 2 | LB | 0 | 0 | 0 | 0 |
| AP_LB_LP | 3 | AP | 8 | 2 | 11 | 18 |
| AP_LB_LP | 3 | LB | 9 | 2 | 28 | 5 |
| AP_LB_LP | 3 | LP | 9 | 2 | 15 | 17 |
| AP_LP | 2 | AP | 3 | 0 | 10 | 13 |
| AP_LP | 2 | LP | 3 | 0 | 14 | 7 |
| AT | 1 | AT | 0 | 0 | 4 | 2 |
| AT_LB | 2 | AT | 0 | 0 | 0 | 0 |
| AT_LB | 2 | LB | 0 | 0 | 0 | 0 |
| AT_LB_LP | 3 | AT | 11 | 2 | 13 | 18 |
| AT_LB_LP | 3 | LB | 12 | 0 | 28 | 7 |
| AT_LB_LP | 3 | LP | 9 | 2 | 18 | 16 |
| AT_LP | 2 | AT | 3 | 0 | 12 | 14 |
| AT_LP | 2 | LP | 3 | 0 | 15 | 9 |
| LB | 1 | LB | 0 | 0 | 0 | 0 |
| LB_LP | 2 | LB | 0 | 0 | 0 | 0 |
| LB_LP | 2 | LP | 0 | 0 | 0 | 0 |
| LP | 1 | LP | 0 | 0 | 0 | 0 |

AF-*Acetobacter fabarum*; AP-*Acetobacter pomorum*; AT-*Acetobacter tropicalis*; LB-*Lactobacillus brevis*; LP-*Lactobacillus plantarum*

**Table S2D. Summary statistics for Figure 2B**

| | f.value | p.value | Tukey comparison | p.value |
|---|---|---|---|---|
| **Rich** | 27.11 | 6.27E-14 | four-five | 0.9457374 |
| | | | one-five | 0 |
| | | | three-five | 0.3474896 |
| | | | two-five | 0.0005505 |
| | | | one-four | 0 |
| | | | three-four | 0.3658145 |
| | | | two-four | 0.0000039 |
| | | | three-one | 0 |
| | | | two-one | 0.0000047 |
| | | | two-three | 0.0005415 |
| **Base** | 56.06 | 2.00E-16 | four-five | 0.8101439 |
| | | | one-five | 0 |
| | | | three-five | 0.0438476 |
| | | | two-five | 0.0000622 |
| | | | one-four | 0 |
| | | | three-four | 0.0410586 |
| | | | two-four | 0.0000008 |
| | | | three-one | 0 |
| | | | two-one | 0 |
| | | | two-three | 0.0043816 |
| **Minimal** | 12.19 | 0.00000011 | four-five | 0.5243205 |
| | | | one-five | 0.0001562 |
| | | | three-five | 0.0606031 |
| | | | two-five | 0.0000753 |
| | | | one-four | 0.0005432 |
| | | | three-four | 0.359816 |

| | | |
|---|---|---|
| | two-four | 0.0000223 |
| | three-one | 0.0120029 |
| | two-one | 0.8429174 |
| | two-three | 0.002909 |

**Table S3A. Predicted community metabolite use patterns for competitive, parasitic and mutualistic interactions - rich medium.**

| Community size | Community composition | Pairwise comparison | Interaction Type | Single-use | Co-consumed | Cross-fed | Single-produced | Co-produced |
|---|---|---|---|---|---|---|---|---|
| 2 | AF_AP | AF-AP | Competitive | 4 | 25 | 0 | 0 | 6 |
| | AF_AT | AF-AT | Competitive | 10 | 20 | 1 | 2 | 6 |
| | AF_LB | AF-LB | Parasitic | 43 | 11 | 6 | 10 | 1 |
| | AF_LP | AF-LP | Parasitic | 43 | 12 | 7 | 7 | 0 |
| | AP_AT | AP-AT | Competitive | 11 | 20 | 2 | 1 | 6 |
| | AP_LB | AP-LB | Parasitic | 44 | 10 | 7 | 9 | 1 |
| | AP_LP | AP-LP | Parasitic | 42 | 13 | 7 | 7 | 0 |
| | AT_LB | AT-LB | Competitive | 44 | 10 | 3 | 12 | 1 |
| | AT_LP | AT-LP | Competitive | 41 | 14 | 3 | 8 | 1 |
| | LB_LP | LB-LP | Parasitic | 35 | 18 | 4 | 3 | 4 |
| 3 | AF-AP-AT | AF-AP | Competitive | 2 | 25 | 0 | 0 | 6 |
| | | AF-AT | Competitive | 11 | 20 | 1 | 2 | 6 |
| | | AP-AT | Competitive | 11 | 20 | 1 | 2 | 6 |
| | AF-AP-LB | AF-AP | Competitive | 5 | 25 | 0 | 1 | 5 |
| | | AF-LB | Competitive | 41 | 11 | 6 | 9 | 1 |
| | | AP-LB | Competitive | 42 | 10 | 6 | 10 | 1 |
| | AF-AP-LP | AF-AP | Competitive | 5 | 26 | 0 | 1 | 6 |
| | | AF-LP | Competitive | 38 | 13 | 7 | 6 | 0 |
| | | AP-LP | Competitive | 41 | 13 | 7 | 7 | 0 |
| | AF-AT-LB | AF-AT | Parasitic | 18 | 20 | 1 | 1 | 6 |
| | | AF-LB | Competitive | 37 | 11 | 5 | 11 | 1 |
| | | AT-LB | Parasitic | 46 | 12 | 5 | 13 | 1 |
| | AF-AT-LP | AF-AT | Parasitic | 13 | 22 | 1 | 4 | 5 |
| | | AF-LP | Competitive | 33 | 12 | 4 | 7 | 1 |
| | | AT-LP | Parasitic | 40 | 11 | 7 | 7 | 1 |
| | AF-LB-LP | AF-LB | Parasitic | 36 | 12 | 5 | 6 | 2 |
| | | AF-LP | Parasitic | 39 | 16 | 4 | 6 | 0 |
| | | LB-LP | Competitive | 21 | 28 | 5 | 6 | 2 |
| | AP-AT-LB | AP-AT | Parasitic | 13 | 24 | 1 | 4 | 4 |
| | | AP-LB | Competitive | 39 | 10 | 7 | 9 | 1 |
| | | AT-LB | Parasitic | 46 | 12 | 6 | 11 | 1 |
| | AP-AT-LP | AP-AT | Parasitic | 12 | 23 | 1 | 5 | 4 |
| | | AP-LP | Competitive | 36 | 11 | 4 | 7 | 1 |
| | | AT-LP | Parasitic | 38 | 12 | 7 | 6 | 1 |
| | AP-LB-LP | AP-LB | Parasitic | 39 | 10 | 5 | 7 | 2 |
| | | AP-LP | Parasitic | 43 | 13 | 4 | 7 | 0 |
| | | LB-LP | Competitive | 20 | 28 | 5 | 6 | 2 |
| | AT-LB-LP | AT-LB | Competitive | 41 | 11 | 4 | 9 | 1 |
| | | AT-LP | Competitive | 43 | 12 | 4 | 7 | 0 |
| | | LB-LP | Competitive | 22 | 27 | 4 | 4 | 2 |
| 4 | AF-AP-AT-LB | AF-AP | Competitive | 4 | 23 | 0 | 2 | 5 |
| | | AF-AT | Parasitic | 17 | 21 | 1 | 2 | 5 |
| | | AF-LB | Competitive | 37 | 11 | 5 | 11 | 1 |
| | | AP-AT | Parasitic | 13 | 24 | 1 | 4 | 4 |
| | | AP-LB | Competitive | 39 | 10 | 7 | 9 | 1 |
| | | AT-LB | Parasitic | 46 | 12 | 6 | 11 | 1 |
| | AF-AP-AT-LP | AF-AP | Competitive | 3 | 24 | 0 | 1 | 5 |
| | | AF-AT | Parasitic | 14 | 21 | 1 | 4 | 4 |
| | | AF-LP | Competitive | 33 | 12 | 4 | 6 | 1 |
| | | AP-AT | Parasitic | 11 | 23 | 1 | 5 | 4 |
| | | AP-LP | Competitive | 36 | 11 | 4 | 7 | 1 |
| | | AT-LP | Parasitic | 39 | 11 | 7 | 6 | 1 |
| | AF-AP-LB-LP | AF-AP | Parasitic | 6 | 24 | 0 | 1 | 5 |
| | | AF-LB | Parasitic | 35 | 12 | 5 | 6 | 2 |

| | | | | | | | | |
|---|---|---|---|---|---|---|---|---|
| | | AF-LP | Parasitic | 38 | 16 | 4 | 6 | 0 |
| | | AP-LB | Competitive | 37 | 10 | 5 | 7 | 2 |
| | | AP-LP | Competitive | 40 | 14 | 4 | 7 | 0 |
| | | LB-LP | Competitive | 21 | 28 | 5 | 6 | 2 |
| | AF-AT-LB-LP | AF-AT | Parasitic | 18 | 22 | 0 | 3 | 4 |
| | | AF-LB | Competitive | 36 | 11 | 4 | 7 | 1 |
| | | AF-LP | Competitive | 34 | 13 | 4 | 6 | 0 |
| | | AT-LB | Parasitic | 47 | 12 | 3 | 9 | 1 |
| | | AT-LP | Parasitic | 44 | 14 | 4 | 7 | 0 |
| | | LB-LP | Competitive | 18 | 27 | 4 | 5 | 2 |
| | AP-AT-LB-LP | AP-AT | Parasitic | 16 | 23 | 0 | 4 | 4 |
| | | AP-LB | Competitive | 37 | 10 | 5 | 7 | 1 |
| | | AP-LP | Competitive | 37 | 12 | 4 | 7 | 0 |
| | | AT-LB | Parasitic | 47 | 12 | 3 | 9 | 1 |
| | | AT-LP | Parasitic | 45 | 14 | 4 | 7 | 0 |
| | | LB-LP | Competitive | 17 | 28 | 4 | 5 | 2 |
| 5 | AF-AP-AT-LB-LP | AF-AP | Competitive | 3 | 24 | 0 | 1 | 5 |
| | | AF-AT | Parasitic | 18 | 22 | 0 | 3 | 4 |
| | | AF-LB | Competitive | 36 | 11 | 4 | 7 | 1 |
| | | AF-LP | Competitive | 35 | 13 | 3 | 7 | 0 |
| | | AP-AT | Parasitic | 15 | 24 | 0 | 4 | 4 |
| | | AP-LB | Competitive | 36 | 11 | 5 | 7 | 1 |
| | | AP-LP | Competitive | 36 | 13 | 3 | 8 | 0 |
| | | AT-LB | Parasitic | 47 | 12 | 3 | 9 | 1 |
| | | AT-LP | Parasitic | 44 | 14 | 4 | 7 | 0 |
| | | LB-LP | Competitive | 18 | 27 | 4 | 5 | 2 |

AF-*Acetobacter fabarum*; AP-*Acetobacter pomorum*; AT-*Acetobacter tropicalis*; LB-*Lactobacillus brevis*; LP-*Lactobacillus plantarum*

**Table S3B. Predicted community metabolite use patterns for competitive, parasitic and mutualistic interactions - base medium**

| Community size | Community composition | Pairwise comparison | Interaction Type | Single-use | Co-consumed | Cross-fed | Single-produced | Co-produced |
|---|---|---|---|---|---|---|---|---|
| 2 | AF_AP | AF-AP | Competitive | 0 | 14 | 0 | 0 | 6 |
| | AF_AT | AF-AT | Competitive | 3 | 13 | 5 | 1 | 5 |
| | AF_LB | AF-LB | Competitive | 17 | 13 | 5 | 8 | 2 |
| | AF_LP | AF-LP | Parasitic | 12 | 13 | 7 | 5 | 1 |
| | AP_AT | AP-AT | Competitive | 4 | 13 | 6 | 1 | 5 |
| | AP_LB | AP-LB | Competitive | 17 | 13 | 5 | 8 | 2 |
| | AP_LP | AP-LP | Parasitic | 12 | 13 | 7 | 5 | 1 |
| | AT_LB | AT-LB | Competitive | 18 | 12 | 5 | 8 | 1 |
| | AT_LP | AT-LP | Parasitic | 12 | 13 | 5 | 5 | 1 |
| | LB_LP | LB-LP | Competitive | 7 | 21 | 8 | 4 | 0 |
| 3 | AF-AP-AT | AF-AP | Competitive | 1 | 20 | 0 | 0 | 5 |
| | | AF-AT | Competitive | 4 | 13 | 5 | 2 | 5 |
| | | AP-AT | Competitive | 4 | 13 | 6 | 1 | 5 |
| | AF-AP-LB | AF-AP | Competitive | 0 | 21 | 0 | 1 | 6 |
| | | AF-LB | Competitive | 17 | 12 | 6 | 8 | 2 |
| | | AP-LB | Competitive | 17 | 12 | 6 | 9 | 1 |
| | AF-AP-LP | AF-AP | Competitive | 1 | 19 | 0 | 1 | 7 |
| | | AF-LP | Competitive | 14 | 12 | 6 | 5 | 1 |
| | | AP-LP | Competitive | 12 | 13 | 7 | 5 | 1 |
| | AF-AT-LB | AF-AT | Parasitic | 5 | 18 | 3 | 1 | 5 |
| | | AF-LB | Competitive | 21 | 10 | 6 | 10 | 1 |
| | | AT-LB | Parasitic | 17 | 11 | 8 | 12 | 1 |
| | AF-AT-LP | AF-AT | Parasitic | 2 | 18 | 0 | 4 | 5 |
| | | AF-LP | Competitive | 14 | 13 | 4 | 6 | 1 |
| | | AT-LP | Parasitic | 13 | 12 | 5 | 7 | 2 |

| | | | | | | | | |
|---|---|---|---|---|---|---|---|---|
| | AF-LB-LP | AF-LB | Parasitic | 22 | 11 | 4 | 12 | 2 |
| | | AF-LP | Parasitic | 23 | 12 | 4 | 6 | 1 |
| | | LB-LP | Competitive | 6 | 19 | 9 | 9 | 0 |
| | AP-AT-LB | AP-AT | Parasitic | 5 | 18 | 2 | 1 | 5 |
| | | AP-LB | Competitive | 20 | 10 | 6 | 10 | 1 |
| | | AT-LB | Parasitic | 17 | 11 | 8 | 11 | 1 |
| | AP-AT-LP | AP-AT | Parasitic | 2 | 18 | 0 | 4 | 5 |
| | | AP-LP | Competitive | 14 | 13 | 4 | 6 | 1 |
| | | AT-LP | Parasitic | 13 | 12 | 5 | 7 | 2 |
| | AP-LB-LP | AP-LB | Parasitic | 21 | 11 | 4 | 13 | 1 |
| | | AP-LP | Parasitic | 22 | 12 | 4 | 6 | 1 |
| | | LB-LP | Competitive | 6 | 19 | 9 | 8 | 0 |
| | AT-LB-LP | AT-LB | Competitive | 19 | 11 | 4 | 12 | 1 |
| | | AT-LP | Competitive | 19 | 14 | 4 | 5 | 0 |
| | | LB-LP | Competitive | 10 | 18 | 10 | 5 | 0 |
| 4 | AF-AP-AT-LB | AF-AP | Competitive | 2 | 21 | 0 | 0 | 5 |
| | | AF-AT | Parasitic | 5 | 18 | 3 | 1 | 5 |
| | | AF-LB | Competitive | 21 | 10 | 6 | 10 | 1 |
| | | AP-AT | Parasitic | 4 | 19 | 2 | 2 | 5 |
| | | AP-LB | Competitive | 21 | 10 | 6 | 10 | 1 |
| | | AT-LB | Parasitic | 17 | 11 | 8 | 12 | 1 |
| | AF-AP-AT-LP | AF-AP | Competitive | 0 | 19 | 0 | 0 | 5 |
| | | AF-AT | Parasitic | 3 | 17 | 0 | 4 | 5 |
| | | AF-LP | Competitive | 15 | 12 | 4 | 6 | 1 |
| | | AP-AT | Parasitic | 3 | 17 | 0 | 4 | 5 |
| | | AP-LP | Competitive | 15 | 12 | 4 | 6 | 1 |
| | | AT-LP | Parasitic | 13 | 12 | 5 | 7 | 2 |
| | AF-AP-LB-LP | AF-AP | Competitive | 1 | 21 | 0 | 0 | 7 |
| | | AF-LB | Competitive | 21 | 11 | 4 | 12 | 2 |
| | | AF-LP | Competitive | 22 | 12 | 4 | 6 | 1 |
| | | AP-LB | Competitive | 20 | 11 | 4 | 12 | 2 |
| | | AP-LP | Competitive | 22 | 12 | 3 | 7 | 1 |
| | | LB-LP | Competitive | 6 | 19 | 9 | 9 | 0 |
| | AF-AT-LB-LP | AF-AT | Parasitic | 5 | 18 | 3 | 1 | 5 |
| | | AF-LB | Competitive | 22 | 10 | 5 | 10 | 1 |
| | | AF-LP | Competitive | 21 | 12 | 3 | 6 | 2 |
| | | AT-LB | Parasitic | 19 | 11 | 6 | 13 | 1 |
| | | AT-LP | Parasitic | 15 | 14 | 5 | 8 | 2 |
| | | LB-LP | Competitive | 6 | 20 | 5 | 7 | 4 |
| | AP-AT-LB-LP | AP-AT | Parasitic | 5 | 18 | 1 | 1 | 5 |
| | | AP-LB | Competitive | 20 | 10 | 5 | 11 | 1 |
| | | AP-LP | Competitive | 20 | 12 | 2 | 8 | 1 |
| | | AT-LB | Parasitic | 18 | 11 | 7 | 11 | 1 |
| | | AT-LP | Parasitic | 16 | 14 | 4 | 6 | 2 |
| | | LB-LP | Competitive | 6 | 20 | 5 | 7 | 4 |
| 5 | AF-AP-AT-LB-LP | AF-AP | Competitive | 3 | 20 | 0 | 1 | 5 |
| | | AF-AT | Parasitic | 5 | 18 | 3 | 1 | 5 |
| | | AF-LB | Competitive | 22 | 10 | 5 | 11 | 1 |
| | | AF-LP | Competitive | 21 | 12 | 2 | 6 | 2 |
| | | AP-AT | Parasitic | 4 | 19 | 1 | 2 | 6 |
| | | AP-LB | Competitive | 21 | 10 | 5 | 12 | 1 |
| | | AP-LP | Competitive | 18 | 13 | 2 | 7 | 2 |
| | | AT-LB | Parasitic | 18 | 11 | 7 | 13 | 1 |
| | | AT-LP | Parasitic | 15 | 14 | 4 | 8 | 2 |
| | | LB-LP | Competitive | 6 | 20 | 4 | 8 | 4 |

AF-*Acetobacter fabarum*; AP-*Acetobacter pomorum*; AT-*Acetobacter tropicalis*; LB-*Lactobacillus brevis*; LP-*Lactobacillus plantarum*

**Table S3C. Predicted community metabolite use patterns for competitive, parasitic and mutualistic interactions - minimal medium.**

| Community size | Community composition | Pairwise comparison | Interaction Type | Single-use | Co-consumed | Cross-fed | Single-produced | Co-produced |
|---|---|---|---|---|---|---|---|---|
| 2 | AF_AP | AF-AP | Parasitic | 0 | 5 | 10 | 0 | 1 |
| | AF_AT | AF-AT | Parasitic | 1 | 4 | 12 | 1 | 1 |
| | AF_LP | AF-LP | Mutualistic | 2 | 3 | 19 | 2 | 1 |
| | AP_AT | AP-AT | Competitive | 1 | 4 | 12 | 1 | 1 |
| | AP_LP | AP-LP | Parasitic | 2 | 3 | 16 | 2 | 1 |
| | AT_LP | AT-LP | Parasitic | 2 | 3 | 19 | 2 | 1 |
| 3 | AF-AP-AT | AF-AP | Parasitic | 6 | 14 | 0 | 1 | 1 |
| | | AF-AT | Parasitic | 2 | 4 | 13 | 3 | 1 |
| | | AP-AT | Competitive | 1 | 4 | 12 | 5 | 1 |
| | AF-AP-LP | AF-AP | Parasitic | 2 | 10 | 5 | 7 | 3 |
| | | AF-LP | Mutualistic | 7 | 4 | 14 | 2 | 1 |
| | | AP-LP | Parasitic | 4 | 5 | 12 | 4 | 2 |
| | AF-AT-LP | AF-AT | Parasitic | 2 | 10 | 4 | 8 | 7 |
| | | AF-LP | Mutualistic | 10 | 4 | 13 | 5 | 0 |
| | | AT-LP | Parasitic | 4 | 3 | 19 | 5 | 1 |
| | AF-LB-LP | AF-LB | Mutualistic | 14 | 6 | 15 | 6 | 1 |
| | | AF-LP | Mutualistic | 2 | 5 | 17 | 15 | 3 |
| | | LB-LP | Mutualistic | 9 | 8 | 17 | 6 | 0 |
| | AP-AT-LP | AP-AT | Competitive | 2 | 10 | 3 | 6 | 7 |
| | | AP-LP | Parasitic | 8 | 4 | 12 | 3 | 2 |
| | | AT-LP | Parasitic | 3 | 3 | 18 | 4 | 1 |
| | AP-LB-LP | AP-LB | Parasitic | 14 | 5 | 15 | 6 | 1 |
| | | AP-LP | Parasitic | 2 | 5 | 14 | 17 | 2 |
| | | LB-LP | Mutualistic | 8 | 9 | 17 | 3 | 1 |
| | AT-LB-LP | AT-LB | Parasitic | 10 | 8 | 15 | 10 | 0 |
| | | AT-LP | Parasitic | 6 | 5 | 15 | 13 | 3 |
| | | LB-LP | Mutualistic | 8 | 9 | 20 | 3 | 0 |
| 4 | AF-AP-AT-LB | AP-AT | Competitive | 0 | 0 | 0 | 0 | 0 |
| | AF-AP-AT-LP | AF-AP | Parasitic | 1 | 12 | 2 | 7 | 3 |
| | | AF-AT | Parasitic | 3 | 10 | 2 | 6 | 5 |
| | | AF-LP | Mutualistic | 9 | 4 | 13 | 2 | 1 |
| | | AP-AT | Competitive | 2 | 9 | 4 | 7 | 4 |
| | | AP-LP | Parasitic | 8 | 5 | 11 | 3 | 2 |
| | | AT-LP | Parasitic | 7 | 3 | 14 | 5 | 1 |
| | AF-AP-LB-LP | AF-AP | Parasitic | 1 | 11 | 1 | 14 | 3 |
| | | AF-LB | Mutualistic | 21 | 5 | 8 | 7 | 0 |
| | | AF-LP | Mutualistic | 8 | 5 | 9 | 15 | 2 |
| | | AP-LB | Parasitic | 18 | 6 | 9 | 5 | 1 |
| | | AP-LP | Parasitic | 4 | 6 | 11 | 14 | 2 |
| | | LB-LP | Mutualistic | 8 | 8 | 18 | 5 | 0 |
| | AF-AT-LB-LP | AF-AT | Parasitic | 3 | 9 | 5 | 10 | 7 |
| | | AF-LB | Mutualistic | 20 | 8 | 6 | 10 | 0 |
| | | AF-LP | Mutualistic | 12 | 6 | 10 | 14 | 1 |
| | | AT-LB | Parasitic | 12 | 7 | 12 | 13 | 1 |
| | | AT-LP | Parasitic | 7 | 3 | 17 | 16 | 2 |
| | | LB-LP | Mutualistic | 10 | 8 | 20 | 4 | 0 |
| | AP-AT-LB-LP | AP-AT | Competitive | 2 | 11 | 0 | 12 | 6 |
| | | AP-LB | Parasitic | 18 | 7 | 8 | 7 | 1 |
| | | AP-LP | Parasitic | 8 | 5 | 11 | 14 | 2 |
| | | AT-LB | Parasitic | 19 | 6 | 9 | 8 | 1 |
| | | AT-LP | Parasitic | 7 | 5 | 12 | 15 | 2 |
| | | LB-LP | Mutualistic | 9 | 8 | 20 | 4 | 0 |
| 5 | AF-AP-AT-LB-LP | AF-AP | Parasitic | 4 | 11 | 0 | 9 | 7 |
| | | AF-AT | Parasitic | 2 | 11 | 2 | 6 | 6 |

| | | | | | | |
|---|---|---|---|---|---|---|
| AF-LB | Mutualistic | 23 | 7 | 5 | 9 | 1 |
| AF-LP | Mutualistic | 10 | 6 | 10 | 14 | 2 |
| AP-AT | Competitive | 2 | 11 | 0 | 11 | 6 |
| AP-LB | Parasitic | 17 | 7 | 9 | 8 | 1 |
| AP-LP | Parasitic | 8 | 5 | 12 | 15 | 2 |
| AT-LB | Parasitic | 23 | 6 | 5 | 9 | 1 |
| AT-LP | Parasitic | 9 | 5 | 11 | 13 | 2 |
| LB-LP | Mutualistic | 10 | 8 | 20 | 4 | 0 |

AF-*Acetobacter fabarum*; AP-*Acetobacter pomorum*; AT-*Acetobacter tropicalis*; LB-*Lactobacillus brevis*; LP-*Lactobacillus plantarum*

**Table S3D. Summary statistics for Figure 3.**

| | f.value | p.value | Tukey comparison | p.value |
|---|---|---|---|---|
| **Fig 3A (Rich competitive)** | 84.7 | <2e-16 | co_produced-co_consumed | 0 |
| | | | cross_fed-co_consumed | 0 |
| | | | single_produced-co_consumed | 0 |
| | | | single_use-co_consumed | 0.0016512 |
| | | | cross_fed-co_produced | 0.9989412 |
| | | | single_produced-co_produced | 0.5057208 |
| | | | single_use-co_produced | 0 |
| | | | single_produced-cross_fed | 0.6749947 |
| | | | single_use-cross_fed | 0 |
| | | | single_use-single_produced | 0 |
| **Fig 3B (Base competitive)** | 58.56 | <2e-16 | co_produced-co_consumed | 0 |
| | | | cross_fed-co_consumed | 0 |
| | | | single_produced-co_consumed | 0 |
| | | | single_use-co_consumed | 0.0008057 |
| | | | cross_fed-co_produced | 0.6838525 |
| | | | single_produced-co_produced | 0.0444179 |
| | | | single_use-co_produced | 0 |
| | | | single_produced-cross_fed | 0.5763813 |
| | | | single_use-cross_fed | 0 |
| | | | single_use-single_produced | 0.0000001 |
| **Fig 3C (Minimal competitive)** | 2.576 | 0.0622 | co_produced-co_consumed | 0.3776919 |
| | | | cross_fed-co_consumed | 0.9309045 |
| | | | single_produced-co_consumed | 0.9730605 |
| | | | single_use-co_consumed | 0.0576864 |
| | | | cross_fed-co_produced | 0.8330905 |
| | | | single_produced-co_produced | 0.7349723 |
| | | | single_use-co_produced | 0.8358828 |
| | | | single_produced-cross_fed | 0.9997113 |
| | | | single_use-cross_fed | 0.2598555 |
| | | | single_use-single_produced | 0.1893655 |
| **Fig 3D (Rich parasitic)** | 134.2 | <2e-16 | co_produced-co_consumed | 0 |
| | | | cross_fed-co_consumed | 0 |
| | | | single_produced-co_consumed | 0 |
| | | | single_use-co_consumed | 0 |
| | | | cross_fed-co_produced | 0.9966574 |
| | | | single_produced-co_produced | 0.186073 |
| | | | single_use-co_produced | 0 |
| | | | single_produced-cross_fed | 0.3522236 |
| | | | single_use-cross_fed | 0 |
| | | | single_use-single_produced | 0 |
| **Fig 3E (Base parasitic)** | 50.27 | <2e-16 | co_produced-co_consumed | 0 |
| | | | cross_fed-co_consumed | 0 |
| | | | single_produced-co_consumed | 0 |
| | | | single_use-co_consumed | 0.0001462 |

| | | | | |
|---|---|---|---|---|
| | | | cross_fed-co_produced | 0.9879244 |
| | | | single_produced-co_produced | 0.129792 |
| | | | single_use-co_produced | 0 |
| | | | single_produced-cross_fed | 0.3350138 |
| | | | single_use-cross_fed | 0 |
| | | | single_use-single_produced | 0.0000259 |
| **Fig 3F (Minimal parasitic)** | 14.67 | 2.37E-10 | co_produced-co_consumed | 0.0002978 |
| | | | cross_fed-co_consumed | 0.0107213 |
| | | | single_produced-co_consumed | 0.9965538 |
| | | | single_use-co_consumed | 0.9566379 |
| | | | cross_fed-co_produced | 0 |
| | | | single_produced-co_produced | 0.0000684 |
| | | | single_use-co_produced | 0.0040358 |
| | | | single_produced-cross_fed | 0.0312083 |
| | | | single_use-cross_fed | 0.00094 |
| | | | single_use-single_produced | 0.8297301 |
| **Fig 3G (Minimal mutualisitic)** | 33.38 | <2e-16 | co_produced-co_consumed | 0.0003301 |
| | | | cross_fed-co_consumed | 0 |
| | | | single_produced-co_consumed | 0.999416 |
| | | | single_use-co_consumed | 0.0177718 |
| | | | cross_fed-co_produced | 0 |
| | | | single_produced-co_produced | 0.000141 |
| | | | single_use-co_produced | 0 |
| | | | single_produced-cross_fed | 0.0000001 |
| | | | single_use-cross_fed | 0.0066907 |
| | | | single_use-single_produced | 0.0334452 |

**Table S4A. Metabolite use pattern - rich medium.**

| | | Number of times metabolite is used | | | | | | Frequency of metabolite use | | | | |
|---|---|---|---|---|---|---|---|---|---|---|---|---|
| Present in medium | Metabolites | Single-use | Co-consumed | Cross-fed | Single-produced | Co-produced | Total number of times metabolite is used in all simulations | Single-use | Co-consumed | Cross-fed | Single-produced | Co-produced |
| Yes | meso-2,6-Diaminoheptanedioate | 32 | 0 | 0 | 0 | 0 | 32 | 1.0 | 0.0 | 0.0 | 0.0 | 0.0 |
| Yes | D-Alanine | 0 | 0 | 0 | 48 | 24 | 72 | 0.0 | 0.0 | 0.0 | 0.7 | 0.3 |
| Yes | Alanine | 46 | 7 | 0 | 0 | 0 | 53 | 0.9 | 0.1 | 0.0 | 0.0 | 0.0 |
| Yes | Arginine | 0 | 80 | 0 | 0 | 0 | 80 | 0.0 | 1.0 | 0.0 | 0.0 | 0.0 |
| Yes | Asparagine | 0 | 80 | 0 | 0 | 0 | 80 | 0.0 | 1.0 | 0.0 | 0.0 | 0.0 |
| Yes | Aspartate | 45 | 27 | 0 | 0 | 0 | 72 | 0.6 | 0.4 | 0.0 | 0.0 | 0.0 |
| Yes | Cysteine | 32 | 0 | 0 | 0 | 0 | 32 | 1.0 | 0.0 | 0.0 | 0.0 | 0.0 |
| Yes | Glutamine | 7 | 73 | 0 | 0 | 0 | 80 | 0.1 | 0.9 | 0.0 | 0.0 | 0.0 |
| Yes | Glutamate | 33 | 2 | 1 | 0 | 0 | 36 | 0.9 | 0.1 | 0.0 | 0.0 | 0.0 |
| Yes | Glycine | 48 | 24 | 0 | 0 | 0 | 72 | 0.7 | 0.3 | 0.0 | 0.0 | 0.0 |
| Yes | Homocysteine | 0 | 0 | 0 | 32 | 0 | 32 | 0.0 | 0.0 | 0.0 | 1.0 | 0.0 |
| Yes | Histidine | 32 | 0 | 0 | 0 | 0 | 32 | 1.0 | 0.0 | 0.0 | 0.0 | 0.0 |
| Yes | Isoleucine | 48 | 8 | 0 | 0 | 0 | 56 | 0.9 | 0.1 | 0.0 | 0.0 | 0.0 |
| Yes | Leucine | 49 | 7 | 0 | 0 | 0 | 56 | 0.9 | 0.1 | 0.0 | 0.0 | 0.0 |
| Yes | Lysine | 48 | 8 | 0 | 0 | 0 | 56 | 0.9 | 0.1 | 0.0 | 0.0 | 0.0 |
| Yes | Methionine | 15 | 0 | 6 | 26 | 7 | 54 | 0.3 | 0.0 | 0.1 | 0.5 | 0.1 |
| Yes | Ornithine | 0 | 0 | 0 | 48 | 25 | 73 | 0.0 | 0.0 | 0.0 | 0.7 | 0.3 |
| Yes | Phenylalanine | 0 | 80 | 0 | 0 | 0 | 80 | 0.0 | 1.0 | 0.0 | 0.0 | 0.0 |
| Yes | Proline | 1 | 79 | 0 | 0 | 0 | 80 | 0.0 | 1.0 | 0.0 | 0.0 | 0.0 |
| Yes | Serine | 0 | 48 | 32 | 0 | 0 | 80 | 0.0 | 0.6 | 0.4 | 0.0 | 0.0 |
| Yes | Threonine | 48 | 8 | 0 | 0 | 0 | 56 | 0.9 | 0.1 | 0.0 | 0.0 | 0.0 |
| Yes | Tryptophan | 0 | 80 | 0 | 0 | 0 | 80 | 0.0 | 1.0 | 0.0 | 0.0 | 0.0 |
| Yes | Tyrosine | 1 | 79 | 0 | 0 | 0 | 80 | 0.0 | 1.0 | 0.0 | 0.0 | 0.0 |
| Yes | Valine | 48 | 8 | 0 | 0 | 0 | 56 | 0.9 | 0.1 | 0.0 | 0.0 | 0.0 |
| Yes | (2-Aminoethyl)phosphonate | 32 | 0 | 0 | 0 | 0 | 32 | 1.0 | 0.0 | 0.0 | 0.0 | 0.0 |
| Yes | 2-Dehydro-3-deoxy-D-gluconate | 1 | 0 | 0 | 0 | 0 | 1 | 1.0 | 0.0 | 0.0 | 0.0 | 0.0 |
| Yes | L-2-hydroxyisocaproate | 1 | 0 | 0 | 0 | 0 | 1 | 1.0 | 0.0 | 0.0 | 0.0 | 0.0 |
| Yes | (R)-3-(4-Hydroxyphenyl)lactate | 1 | 0 | 0 | 0 | 0 | 1 | 1.0 | 0.0 | 0.0 | 0.0 | 0.0 |
| Yes | 4-Aminobutanoate | 47 | 6 | 0 | 0 | 0 | 53 | 0.9 | 0.1 | 0.0 | 0.0 | 0.0 |
| Yes | Acetate | 5 | 12 | 29 | 19 | 8 | 73 | 0.1 | 0.2 | 0.4 | 0.3 | 0.1 |
| Yes | Acetaldehyde | 50 | 6 | 0 | 0 | 0 | 56 | 0.9 | 0.1 | 0.0 | 0.0 | 0.0 |
| Yes | N-Acetyl-D-glucosamine | 50 | 22 | 0 | 0 | 0 | 72 | 0.7 | 0.3 | 0.0 | 0.0 | 0.0 |
| Yes | R Acetoin | 24 | 11 | 24 | 15 | 0 | 74 | 0.3 | 0.1 | 0.3 | 0.2 | 0.0 |
| Yes | S Acetoin | 46 | 2 | 0 | 0 | 0 | 48 | 1.0 | 0.0 | 0.0 | 0.0 | 0.0 |

| | | | | | | | | | | | |
|---|---|---|---|---|---|---|---|---|---|---|---|
| Yes | 2-Oxoglutarate | 5 | 3 | 28 | 20 | 24 | 80 | 0.1 | 0.0 | 0.4 | 0.3 | 0.3 |
| Yes | (R,R)-2,3-Butanediol | 39 | 0 | 0 | 0 | 0 | 39 | 1.0 | 0.0 | 0.0 | 0.0 | 0.0 |
| Yes | (S,S)-2,3-Butanediol | 45 | 9 | 0 | 0 | 0 | 54 | 0.8 | 0.2 | 0.0 | 0.0 | 0.0 |
| Yes | Citrate | 2 | 0 | 0 | 0 | 0 | 2 | 1.0 | 0.0 | 0.0 | 0.0 | 0.0 |
| Yes | Ethanol | 30 | 0 | 0 | 0 | 0 | 30 | 1.0 | 0.0 | 0.0 | 0.0 | 0.0 |
| Yes | Formaldehyde | | | | | | 0 | | | | | |
| Yes | Formate | 0 | 0 | 0 | 19 | 2 | 21 | 0.0 | 0.0 | 0.0 | 0.9 | 0.1 |
| Yes | D-Fructose | 25 | 0 | 0 | 0 | 0 | 25 | 1.0 | 0.0 | 0.0 | 0.0 | 0.0 |
| Yes | Fumarate | 1 | 0 | 0 | 0 | 0 | 1 | 1.0 | 0.0 | 0.0 | 0.0 | 0.0 |
| Yes | D-Glucose | 18 | 7 | 0 | 0 | 0 | 25 | 0.7 | 0.3 | 0.0 | 0.0 | 0.0 |
| Yes | D-Gluconate | 6 | 0 | 0 | 0 | 0 | 6 | 1.0 | 0.0 | 0.0 | 0.0 | 0.0 |
| Yes | Glycerol | 46 | 15 | 4 | 3 | 0 | 68 | 0.7 | 0.2 | 0.1 | 0.0 | 0.0 |
| Yes | Glycerol 3-phosphate | 48 | 4 | 0 | 0 | 0 | 52 | 0.9 | 0.1 | 0.0 | 0.0 | 0.0 |
| Yes | Glycolate | | | | | | 0 | | | | | |
| Yes | Imidazole lactate | 32 | 0 | 0 | 0 | 0 | 32 | 1.0 | 0.0 | 0.0 | 0.0 | 0.0 |
| Yes | D-Lactate | 0 | 0 | 0 | 6 | 0 | 6 | 0.0 | 0.0 | 0.0 | 1.0 | 0.0 |
| Yes | L-Lactate | 0 | 0 | 0 | 6 | 0 | 6 | 0.0 | 0.0 | 0.0 | 1.0 | 0.0 |
| Yes | Malate | 1 | 0 | 0 | 37 | 12 | 50 | 0.0 | 0.0 | 0.0 | 0.7 | 0.2 |
| Yes | Maltose | 42 | 6 | 0 | 0 | 0 | 48 | 0.9 | 0.1 | 0.0 | 0.0 | 0.0 |
| Yes | Maltotetraose | 32 | 0 | 0 | 0 | 0 | 32 | 1.0 | 0.0 | 0.0 | 0.0 | 0.0 |
| Yes | D-Mannose | 20 | 0 | 0 | 0 | 0 | 20 | 1.0 | 0.0 | 0.0 | 0.0 | 0.0 |
| Yes | D-Mannitol | 23 | 0 | 0 | 0 | 0 | 23 | 1.0 | 0.0 | 0.0 | 0.0 | 0.0 |
| Yes | Methylglyoxal | 35 | 2 | 0 | 0 | 0 | 37 | 0.9 | 0.1 | 0.0 | 0.0 | 0.0 |
| Yes | Pyruvate | | | | | | 0 | | | | | |
| Yes | D-Sorbitol | 14 | 0 | 0 | 0 | 0 | 14 | 1.0 | 0.0 | 0.0 | 0.0 | 0.0 |
| Yes | Succinate | 13 | 0 | 19 | 24 | 6 | 62 | 0.2 | 0.0 | 0.3 | 0.4 | 0.1 |
| Yes | Succinyl-CoA | 32 | 48 | 0 | 0 | 0 | 80 | 0.4 | 0.6 | 0.0 | 0.0 | 0.0 |
| Yes | Sucrose | 51 | 9 | 0 | 0 | 0 | 60 | 0.9 | 0.2 | 0.0 | 0.0 | 0.0 |
| Yes | Trehalose | 49 | 7 | 0 | 0 | 0 | 56 | 0.9 | 0.1 | 0.0 | 0.0 | 0.0 |
| Yes | Adenine | 48 | 8 | 0 | 0 | 0 | 56 | 0.9 | 0.1 | 0.0 | 0.0 | 0.0 |
| Yes | Adenosine | 32 | 0 | 0 | 0 | 0 | 32 | 1.0 | 0.0 | 0.0 | 0.0 | 0.0 |
| Yes | Cytosine | 1 | 0 | 0 | 0 | 0 | 1 | 1.0 | 0.0 | 0.0 | 0.0 | 0.0 |
| Yes | Cytidine | 32 | 0 | 0 | 0 | 0 | 32 | 1.0 | 0.0 | 0.0 | 0.0 | 0.0 |
| Yes | Deoxyadenosine | 26 | 0 | 0 | 0 | 0 | 26 | 1.0 | 0.0 | 0.0 | 0.0 | 0.0 |
| Yes | Deoxycytidine | 40 | 1 | 0 | 0 | 0 | 41 | 1.0 | 0.0 | 0.0 | 0.0 | 0.0 |
| Yes | Deoxyribose | 3 | 0 | 0 | 0 | 0 | 3 | 1.0 | 0.0 | 0.0 | 0.0 | 0.0 |
| No | dUMP | 12 | 19 | 39 | 7 | 0 | 77 | 0.2 | 0.2 | 0.5 | 0.1 | 0.0 |
| Yes | Deoxyuridine | 29 | 1 | 14 | 20 | 0 | 64 | 0.5 | 0.0 | 0.2 | 0.3 | 0.0 |
| Yes | Guanine | 48 | 8 | 0 | 0 | 0 | 56 | 0.9 | 0.1 | 0.0 | 0.0 | 0.0 |
| Yes | Hypoxanthine | 19 | 6 | 12 | 5 | 0 | 42 | 0.5 | 0.1 | 0.3 | 0.1 | 0.0 |
| Yes | Inosine | 48 | 0 | 0 | 0 | 0 | 48 | 1.0 | 0.0 | 0.0 | 0.0 | 0.0 |

| | | | | | | | | | | | | |
|---|---|---|---|---|---|---|---|---|---|---|---|---|
| Yes | Orotate | | | | | | 0 | | | | | |
| Yes | Thymidine | 48 | 8 | 0 | 0 | 0 | 56 | 0.9 | 0.1 | 0.0 | 0.0 | 0.0 |
| Yes | Uracil | 4 | 20 | 46 | 2 | 8 | 80 | 0.1 | 0.3 | 0.6 | 0.0 | 0.1 |
| Yes | Uridine | 45 | 8 | 3 | 0 | 0 | 56 | 0.8 | 0.1 | 0.1 | 0.0 | 0.0 |
| Yes | Xanthine | 8 | 2 | 9 | 23 | 0 | 42 | 0.2 | 0.0 | 0.2 | 0.5 | 0.0 |
| Yes | Biotin (B7) | 0 | 80 | 0 | 0 | 0 | 80 | 0.0 | 1.0 | 0.0 | 0.0 | 0.0 |
| Yes | Coenzyme A | 24 | 0 | 8 | 24 | 0 | 56 | 0.4 | 0.0 | 0.1 | 0.4 | 0.0 |
| Yes | Dihydropteroate | | | | | | 0 | | | | | |
| Yes | 1-deoxy-D-xylulose 5-phosphate | 32 | 0 | 0 | 0 | 0 | 32 | 1.0 | 0.0 | 0.0 | 0.0 | 0.0 |
| Yes | Folate (B9) | 48 | 8 | 0 | 0 | 0 | 56 | 0.9 | 0.1 | 0.0 | 0.0 | 0.0 |
| Yes | Nicotinate | | | | | | 0 | | | | | |
| Yes | Nicotinamide D-ribonucleotide | 0 | 80 | 0 | 0 | 0 | 80 | 0.0 | 1.0 | 0.0 | 0.0 | 0.0 |
| Yes | Pyridoxine 5-phosphate (B6) | 48 | 24 | 0 | 0 | 0 | 72 | 0.7 | 0.3 | 0.0 | 0.0 | 0.0 |
| Yes | Pantothenate (B5) | | | | | | 0 | | | | | |
| Yes | Pyridoxamine (B6) | 0 | 0 | 0 | 44 | 0 | 44 | 0.0 | 0.0 | 0.0 | 1.0 | 0.0 |
| Yes | Pyridoxal 5'-phosphate (B6) | 48 | 8 | 0 | 0 | 0 | 56 | 0.9 | 0.1 | 0.0 | 0.0 | 0.0 |
| Yes | Riboflavin (B2) | 48 | 24 | 0 | 0 | 0 | 72 | 0.7 | 0.3 | 0.0 | 0.0 | 0.0 |
| Yes | Tetrahydrofolate (B9) | 48 | 24 | 0 | 0 | 0 | 72 | 0.7 | 0.3 | 0.0 | 0.0 | 0.0 |
| Yes | Thiamin (B1) | 24 | 0 | 8 | 24 | 0 | 56 | 0.4 | 0.0 | 0.1 | 0.4 | 0.0 |
| No | Toxopyrimidine | 32 | 0 | 0 | 0 | 0 | 32 | 1.0 | 0.0 | 0.0 | 0.0 | 0.0 |
| Yes | Ammonium | 0 | 0 | 0 | 23 | 57 | 80 | 0.0 | 0.0 | 0.0 | 0.3 | 0.7 |
| No | L-Cysteinylglycine | 49 | 7 | 0 | 0 | 0 | 56 | 0.9 | 0.1 | 0.0 | 0.0 | 0.0 |
| Yes | L-methionyl-L-alanine | 2 | 78 | 0 | 0 | 0 | 80 | 0.0 | 1.0 | 0.0 | 0.0 | 0.0 |
| Yes | Hydrogen sulfide | 48 | 24 | 0 | 0 | 0 | 72 | 0.7 | 0.3 | 0.0 | 0.0 | 0.0 |
| Yes | Sulfate | 48 | 24 | 0 | 0 | 0 | 72 | 0.7 | 0.3 | 0.0 | 0.0 | 0.0 |

**Table S4B. Metabolite use pattern - base medium.**

| | | Number of times metabolite is used | | | | | | Frequency of metabolite use | | | | |
|---|---|---|---|---|---|---|---|---|---|---|---|---|
| **Present in medium** | **Metabolites** | **Single-use** | **Co-consumed** | **Cross-fed** | **Single-produced** | **Co-produced** | **Total number of times metabolite is used in all simulations** | **Single-use** | **Co-consumed** | **Cross-fed** | **Single-produced** | **Co-produced** |
| Yes | meso-2,6-Diaminoheptanedioate | 32 | 0 | 0 | 0 | 0 | 32 | 1.0 | 0.0 | 0.0 | 0.0 | 0.0 |
| No | D-Alanine | 0 | 0 | 0 | 48 | 24 | 72 | 0.0 | 0.0 | 0.0 | 0.7 | 0.3 |
| Yes | Alanine | 48 | 7 | 0 | 0 | 0 | 55 | 0.9 | 0.1 | 0.0 | 0.0 | 0.0 |
| Yes | Arginine | 0 | 80 | 0 | 0 | 0 | 80 | 0.0 | 1.0 | 0.0 | 0.0 | 0.0 |
| Yes | Asparagine | 0 | 80 | 0 | 0 | 0 | 80 | 0.0 | 1.0 | 0.0 | 0.0 | 0.0 |
| Yes | Aspartate | 37 | 20 | 0 | 0 | 0 | 57 | 0.6 | 0.4 | 0.0 | 0.0 | 0.0 |
| Yes | Cysteine | 48 | 8 | 0 | 0 | 0 | 56 | 0.9 | 0.1 | 0.0 | 0.0 | 0.0 |

| | | | | | | | | | | | | |
|---|---|---|---|---|---|---|---|---|---|---|---|---|
| Yes | Glutamine | 0 | 80 | 0 | 0 | 0 | 80 | 0.0 | 1.0 | 0.0 | 0.0 | 0.0 |
| Yes | Glutamate | 5 | 0 | 5 | 5 | 0 | 15 | 0.3 | 0.0 | 0.3 | 0.3 | 0.0 |
| Yes | Glycine | 18 | 18 | 34 | 3 | 1 | 74 | 0.2 | 0.2 | 0.5 | 0.0 | 0.0 |
| No | Homocysteine | 0 | 0 | 0 | 32 | 0 | 32 | 0.0 | 0.0 | 0.0 | 1.0 | 0.0 |
| Yes | Histidine | 32 | 0 | 0 | 0 | 0 | 32 | 1.0 | 0.0 | 0.0 | 0.0 | 0.0 |
| Yes | Isoleucine | 48 | 8 | 0 | 0 | 0 | 56 | 0.9 | 0.1 | 0.0 | 0.0 | 0.0 |
| Yes | Leucine | 48 | 8 | 0 | 0 | 0 | 56 | 0.9 | 0.1 | 0.0 | 0.0 | 0.0 |
| Yes | Lysine | 48 | 8 | 0 | 0 | 0 | 56 | 0.9 | 0.1 | 0.0 | 0.0 | 0.0 |
| Yes | Methionine | 0 | 80 | 0 | 0 | 0 | 80 | 0.0 | 1.0 | 0.0 | 0.0 | 0.0 |
| No | Ornithine | 0 | 0 | 0 | 37 | 38 | 75 | 0.0 | 0.0 | 0.0 | 0.5 | 0.5 |
| Yes | Phenylalanine | 0 | 80 | 0 | 0 | 0 | 80 | 0.0 | 1.0 | 0.0 | 0.0 | 0.0 |
| Yes | Proline | 0 | 80 | 0 | 0 | 0 | 80 | 0.0 | 1.0 | 0.0 | 0.0 | 0.0 |
| Yes | Serine | 0 | 54 | 26 | 0 | 0 | 80 | 0.0 | 0.7 | 0.3 | 0.0 | 0.0 |
| Yes | Threonine | 48 | 8 | 0 | 0 | 0 | 56 | 0.9 | 0.1 | 0.0 | 0.0 | 0.0 |
| Yes | Tryptophan | 0 | 80 | 0 | 0 | 0 | 80 | 0.0 | 1.0 | 0.0 | 0.0 | 0.0 |
| Yes | Tyrosine | 0 | 80 | 0 | 0 | 0 | 80 | 0.0 | 1.0 | 0.0 | 0.0 | 0.0 |
| Yes | Valine | 48 | 8 | 0 | 0 | 0 | 56 | 0.9 | 0.1 | 0.0 | 0.0 | 0.0 |
| No | (2-Aminoethyl)phosphonate | | | | | | 0 | | | | | |
| No | 2-Dehydro-3-deoxy-D-gluconate | | | | | | 0 | | | | | |
| No | L-2-hydroxyisocaproate | | | | | | 0 | | | | | |
| No | (R)-3-(4-Hydroxyphenyl)lactate | | | | | | 0 | | | | | |
| No | 4-Aminobutanoate | | | | | | 0 | | | | | |
| No | Acetate | 5 | 19 | 47 | 5 | 3 | 79 | 0.1 | 0.2 | 0.6 | 0.1 | 0.0 |
| No | Acetaldehyde | 13 | 0 | 11 | 16 | 2 | 42 | 0.3 | 0.0 | 0.3 | 0.4 | 0.0 |
| No | N-Acetyl-D-glucosamine | | | | | | 0 | | | | | |
| No | R Acetoin | 17 | 4 | 25 | 19 | 3 | 68 | 0.3 | 0.1 | 0.4 | 0.3 | 0.0 |
| No | S Acetoin | | | | | | 0 | | | | | |
| No | 2-Oxoglutarate | 1 | 0 | 6 | 33 | 39 | 79 | 0.0 | 0.0 | 0.1 | 0.4 | 0.5 |
| No | (R,R)-2,3-Butanediol | | | | | | 0 | | | | | |
| No | (S,S)-2,3-Butanediol | | | | | | 0 | | | | | |
| No | Citrate | | | | | | 0 | | | | | |
| No | Ethanol | | | | | | 0 | | | | | |
| No | Formaldehyde | | | | | | 0 | | | | | |
| No | Formate | 0 | 0 | 0 | 42 | 31 | 73 | 0.0 | 0.0 | 0.0 | 0.6 | 0.4 |
| No | D-Fructose | | | | | | 0 | | | | | |
| No | Fumarate | | | | | | 0 | | | | | |
| Yes | D-Glucose | 0 | 80 | 0 | 0 | 0 | 80 | 0.0 | 1.0 | 0.0 | 0.0 | 0.0 |
| No | D-Gluconate | | | | | | 0 | | | | | |
| Yes | Glycerol | 46 | 19 | 0 | 0 | 0 | 65 | 0.7 | 0.3 | 0.0 | 0.0 | 0.0 |
| No | Glycerol 3-phosphate | | | | | | 0 | | | | | |
| No | Glycolate | 0 | 0 | 0 | 39 | 23 | 62 | 0.0 | 0.0 | 0.0 | 0.6 | 0.4 |

| | | | | | | | | | | | | |
|---|---|---|---|---|---|---|---|---|---|---|---|---|
| No | Imidazole lactate | 12 | 0 | 8 | 12 | 0 | 32 | 0.4 | 0.0 | 0.3 | 0.4 | 0.0 |
| No | D-Lactate | | | | | | 0 | | | | | |
| No | L-Lactate | | | | | | 0 | | | | | |
| No | Malate | 0 | 0 | 0 | 17 | 3 | 20 | 0.0 | 0.0 | 0.0 | 0.9 | 0.2 |
| No | Maltose | | | | | | 0 | | | | | |
| No | Maltotetraose | | | | | | 0 | | | | | |
| No | D-Mannose | | | | | | 0 | | | | | |
| No | D-Mannitol | | | | | | 0 | | | | | |
| No | Methylglyoxal | | | | | | 0 | | | | | |
| No | Pyruvate | | | | | | 0 | | | | | |
| No | D-Sorbitol | | | | | | 0 | | | | | |
| No | Succinate | 4 | 0 | 16 | 4 | 7 | 31 | 0.1 | 0.0 | 0.5 | 0.1 | 0.2 |
| No | Succinyl-CoA | 17 | 3 | 19 | 14 | 0 | 53 | 0.3 | 0.1 | 0.4 | 0.3 | 0.0 |
| No | Sucrose | | | | | | 0 | | | | | |
| No | Trehalose | | | | | | 0 | | | | | |
| No | Adenine | 5 | 0 | 5 | 5 | 0 | 15 | 0.3 | 0.0 | 0.3 | 0.3 | 0.0 |
| No | Adenosine | | | | | | 0 | | | | | |
| No | Cytosine | | | | | | 0 | | | | | |
| No | Cytidine | | | | | | 0 | | | | | |
| No | Deoxyadenosine | | | | | | 0 | | | | | |
| No | Deoxycytidine | | | | | | 0 | | | | | |
| No | Deoxyribose | | | | | | 0 | | | | | |
| No | dUMP | 11 | 13 | 26 | 6 | 0 | 56 | 0.2 | 0.2 | 0.5 | 0.1 | 0.0 |
| No | Deoxyuridine | | | | | | 0 | | | | | |
| No | Guanine | | | | | | 0 | | | | | |
| No | Hypoxanthine | | | | | | 0 | | | | | |
| No | Inosine | | | | | | 0 | | | | | |
| No | Orotate | 0 | 24 | 32 | 0 | 0 | 56 | 0.0 | 0.4 | 0.6 | 0.0 | 0.0 |
| No | Thymidine | | | | | | 0 | | | | | |
| No | Uracil | | | | | | 0 | | | | | |
| No | Uridine | | | | | | 0 | | | | | |
| No | Xanthine | | | | | | 0 | | | | | |
| Yes | Biotin (B7) | 0 | 80 | 0 | 0 | 0 | 80 | 0.0 | 1.0 | 0.0 | 0.0 | 0.0 |
| No | Coenzyme A | 5 | 0 | 5 | 5 | 0 | 15 | 0.3 | 0.0 | 0.3 | 0.3 | 0.0 |
| Yes | Dihydropteroate | 32 | 0 | 0 | 0 | 0 | 32 | 1.0 | 0.0 | 0.0 | 0.0 | 0.0 |
| Yes | 1-deoxy-D-xylulose 5-phosphate | 32 | 0 | 0 | 0 | 0 | 32 | 1.0 | 0.0 | 0.0 | 0.0 | 0.0 |
| No | Folate (B9) | 12 | 0 | 8 | 12 | 0 | 32 | 0.4 | 0.0 | 0.3 | 0.4 | 0.0 |
| Yes | Nicotinate | 48 | 8 | 0 | 0 | 0 | 56 | 0.9 | 0.1 | 0.0 | 0.0 | 0.0 |
| No | Nicotinamide D-ribonucleotide | 12 | 18 | 36 | 7 | 0 | 73 | 0.2 | 0.2 | 0.5 | 0.1 | 0.0 |
| No | Pyridoxine 5-phosphate (B6) | 0 | 1 | 4 | 0 | 0 | 5 | 0.0 | 0.2 | 0.8 | 0.0 | 0.0 |
| Yes | Pantothenate (B5) | 46 | 31 | 0 | 0 | 0 | 77 | 0.6 | 0.4 | 0.0 | 0.0 | 0.0 |

| | | | | | | | | | | | | |
|---|---|---|---|---|---|---|---|---|---|---|---|---|
| Yes | Pyridoxamine (B6) | 0 | 0 | 0 | 48 | 3 | 51 | 0.0 | 0.0 | 0.0 | 0.9 | 0.1 |
| Yes | Pyridoxal 5'-phosphate (B6) | 48 | 8 | 0 | 0 | 0 | 56 | 0.9 | 0.1 | 0.0 | 0.0 | 0.0 |
| No | Riboflavin (B2) | 8 | 1 | 8 | 8 | 0 | 25 | 0.3 | 0.0 | 0.3 | 0.3 | 0.0 |
| No | Tetrahydrofolate (B9) | 9 | 2 | 10 | 8 | 0 | 29 | 0.3 | 0.1 | 0.3 | 0.3 | 0.0 |
| No | Thiamin (B1) | 9 | 0 | 7 | 25 | 0 | 41 | 0.2 | 0.0 | 0.2 | 0.6 | 0.0 |
| No | Toxopyrimidine | 32 | 0 | 0 | 0 | 0 | 32 | 1.0 | 0.0 | 0.0 | 0.0 | 0.0 |
| Yes | Ammonium | 0 | 0 | 0 | 48 | 24 | 72 | 0.0 | 0.0 | 0.0 | 0.7 | 0.3 |
| No | L-Cysteinylglycine | | | | | | 0 | | | | | |
| No | L-methionyl-L-alanine | | | | | | 0 | | | | | |
| Yes | Hydrogen sulfide | 48 | 24 | 0 | 0 | 0 | 72 | 0.7 | 0.3 | 0.0 | 0.0 | 0.0 |
| Yes | Sulfate | 48 | 23 | 0 | 0 | 0 | 71 | 0.7 | 0.3 | 0.0 | 0.0 | 0.0 |

**Table S4C. Metabolite use pattern - minimal medium.**

| | | Number of times metabolite is used | | | | | | Frequency of metabolite use | | | | |
|---|---|---|---|---|---|---|---|---|---|---|---|---|
| Present in medium | Metabolite name | Single-use | Co-consumed | Cross-fed | Single-produced | Co-produced | Total number of times metabolite is used in all simulations | Single-use | Co-consumed | Cross-fed | Single-produced | Co-produced |
| No | meso-2,6-Diaminoheptanedioate | 12 | 0 | 7 | 12 | 0 | 31 | 0.4 | 0.0 | 0.2 | 0.4 | 0.0 |
| No | D-Alanine | 0 | 0 | 0 | 36 | 18 | 54 | 0.0 | 0.0 | 0.0 | 0.7 | 0.3 |
| No | Alanine | | | | | | 0 | | | | | |
| No | Arginine | 2 | 21 | 34 | 2 | 1 | 60 | 0.0 | 0.4 | 0.6 | 0.0 | 0.0 |
| No | Asparagine | 13 | 8 | 22 | 10 | 0 | 53 | 0.2 | 0.2 | 0.4 | 0.2 | 0.0 |
| No | Aspartate | | | | | | 0 | | | | | |
| No | Cysteine | 12 | 0 | 7 | 12 | 0 | 31 | 0.4 | 0.0 | 0.2 | 0.4 | 0.0 |
| No | Glutamine | 1 | 0 | 1 | 1 | 0 | 3 | 0.3 | 0.0 | 0.3 | 0.3 | 0.0 |
| No | Glutamate | 11 | 0 | 8 | 11 | 1 | 31 | 0.4 | 0.0 | 0.3 | 0.4 | 0.0 |
| No | Glycine | 14 | 7 | 18 | 9 | 0 | 48 | 0.3 | 0.1 | 0.4 | 0.2 | 0.0 |
| No | Homocysteine | 0 | 0 | 0 | 19 | 0 | 19 | 0.0 | 0.0 | 0.0 | 1.0 | 0.0 |
| No | Histidine | 11 | 0 | 8 | 12 | 1 | 32 | 0.3 | 0.0 | 0.3 | 0.4 | 0.0 |
| No | Isoleucine | 8 | 7 | 28 | 4 | 7 | 54 | 0.1 | 0.1 | 0.5 | 0.1 | 0.1 |
| No | Leucine | 11 | 7 | 25 | 7 | 3 | 53 | 0.2 | 0.1 | 0.5 | 0.1 | 0.1 |
| No | Lysine | 12 | 0 | 7 | 12 | 0 | 31 | 0.4 | 0.0 | 0.2 | 0.4 | 0.0 |
| No | Methionine | 12 | 0 | 7 | 12 | 0 | 31 | 0.4 | 0.0 | 0.2 | 0.4 | 0.0 |
| No | Ornithine | 2 | 1 | 34 | 2 | 21 | 60 | 0.0 | 0.0 | 0.6 | 0.0 | 0.4 |
| No | Phenylalanine | 0 | 25 | 35 | 0 | 0 | 60 | 0.0 | 0.4 | 0.6 | 0.0 | 0.0 |
| No | Proline | 3 | 18 | 24 | 3 | 1 | 49 | 0.1 | 0.4 | 0.5 | 0.1 | 0.0 |
| No | Serine | 0 | 24 | 31 | 0 | 0 | 55 | 0.0 | 0.4 | 0.6 | 0.0 | 0.0 |
| No | Threonine | 12 | 0 | 7 | 12 | 0 | 31 | 0.4 | 0.0 | 0.2 | 0.4 | 0.0 |
| No | Tryptophan | 13 | 6 | 25 | 11 | 1 | 56 | 0.2 | 0.1 | 0.4 | 0.2 | 0.0 |

| | | | | | | | | | | | | |
|---|---|---|---|---|---|---|---|---|---|---|---|---|
| No | Tyrosine | 3 | 21 | 31 | 3 | 0 | 58 | 0.1 | 0.4 | 0.5 | 0.1 | 0.0 |
| No | Valine | 13 | 7 | 23 | 9 | 1 | 53 | 0.2 | 0.1 | 0.4 | 0.2 | 0.0 |
| No | (2-Aminoethyl)phosphonate | | | | | | 0 | | | | | |
| No | 2-Dehydro-3-deoxy-D-gluconate | | | | | | 0 | | | | | |
| No | L-2-hydroxyisocaproate | | | | | | 0 | | | | | |
| No | (R)-3-(4-Hydroxyphenyl)lactate | 7 | 0 | 3 | 7 | 0 | 17 | 0.4 | 0.0 | 0.2 | 0.4 | 0.0 |
| No | 4-Aminobutanoate | | | | | | 0 | | | | | |
| No | Acetate | 12 | 13 | 27 | 7 | 0 | 59 | 0.2 | 0.2 | 0.5 | 0.1 | 0.0 |
| No | Acetaldehyde | 2 | 0 | 3 | 2 | 0 | 7 | 0.3 | 0.0 | 0.4 | 0.3 | 0.0 |
| No | N-Acetyl-D-glucosamine | | | | | | 0 | | | | | |
| No | R Acetoin | 5 | 0 | 9 | 5 | 1 | 20 | 0.3 | 0.0 | 0.5 | 0.3 | 0.1 |
| No | S Acetoin | | | | | | 0 | | | | | |
| No | 2-Oxoglutarate | 6 | 0 | 29 | 9 | 16 | 60 | 0.1 | 0.0 | 0.5 | 0.2 | 0.3 |
| No | (R,R)-2,3-Butanediol | | | | | | 0 | | | | | |
| No | (S,S)-2,3-Butanediol | | | | | | 0 | | | | | |
| No | Citrate | | | | | | 0 | | | | | |
| No | Ethanol | | | | | | 0 | | | | | |
| No | Formaldehyde | 0 | 0 | 0 | 34 | 6 | 40 | 0.0 | 0.0 | 0.0 | 0.9 | 0.2 |
| No | Formate | 12 | 0 | 8 | 12 | 0 | 32 | 0.4 | 0.0 | 0.3 | 0.4 | 0.0 |
| No | D-Fructose | | | | | | 0 | | | | | |
| No | Fumarate | | | | | | 0 | | | | | |
| Yes | D-Glucose | 0 | 61 | 0 | 0 | 0 | 61 | 0.0 | 1.0 | 0.0 | 0.0 | 0.0 |
| No | D-Gluconate | | | | | | 0 | | | | | |
| Yes | Glycerol | 24 | 36 | 0 | 0 | 0 | 60 | 0.4 | 0.6 | 0.0 | 0.0 | 0.0 |
| No | Glycerol 3-phosphate | | | | | | 0 | | | | | |
| No | Glycolate | 0 | 0 | 0 | 26 | 32 | 58 | 0.0 | 0.0 | 0.0 | 0.4 | 0.6 |
| No | Imidazole lactate | 8 | 0 | 4 | 8 | 0 | 20 | 0.4 | 0.0 | 0.2 | 0.4 | 0.0 |
| No | D-Lactate | | | | | | 0 | | | | | |
| No | L-Lactate | | | | | | 0 | | | | | |
| No | Malate | 4 | 0 | 5 | 5 | 2 | 16 | 0.3 | 0.0 | 0.3 | 0.3 | 0.1 |
| No | Maltose | | | | | | 0 | | | | | |
| No | Maltotetraose | | | | | | 0 | | | | | |
| No | D-Mannose | | | | | | 0 | | | | | |
| No | D-Mannitol | | | | | | 0 | | | | | |
| No | Methylglyoxal | | | | | | 0 | | | | | |
| No | Pyruvate | 11 | 0 | 6 | 11 | 0 | 28 | 0.4 | 0.0 | 0.2 | 0.4 | 0.0 |
| No | D-Sorbitol | | | | | | 0 | | | | | |
| No | Succinate | 10 | 0 | 7 | 12 | 1 | 30 | 0.3 | 0.0 | 0.2 | 0.4 | 0.0 |
| No | Succinyl-CoA | 16 | 5 | 23 | 13 | 0 | 57 | 0.3 | 0.1 | 0.4 | 0.2 | 0.0 |
| No | Sucrose | | | | | | 0 | | | | | |

| | | | | | | | | | | | | |
|---|---|---|---|---|---|---|---|---|---|---|---|---|
| No | Trehalose | | | | | | 0 | | | | | |
| No | Adenine | 12 | 0 | 7 | 12 | 0 | 31 | 0.4 | 0.0 | 0.2 | 0.4 | 0.0 |
| No | Adenosine | | | | | | 0 | | | | | |
| No | Cytosine | | | | | | 0 | | | | | |
| No | Cytidine | | | | | | 0 | | | | | |
| No | Deoxyadenosine | | | | | | 0 | | | | | |
| No | Deoxycytidine | | | | | | 0 | | | | | |
| No | Deoxyribose | | | | | | 0 | | | | | |
| No | dUMP | 6 | 16 | 26 | 6 | 0 | 54 | 0.1 | 0.3 | 0.5 | 0.1 | 0.0 |
| No | Deoxyuridine | | | | | | 0 | | | | | |
| No | Guanine | | | | | | 0 | | | | | |
| No | Hypoxanthine | | | | | | 0 | | | | | |
| No | Inosine | | | | | | 0 | | | | | |
| No | Orotate | 5 | 2 | 8 | 4 | 0 | 19 | 0.3 | 0.1 | 0.4 | 0.2 | 0.0 |
| No | Thymidine | | | | | | 0 | | | | | |
| No | Uracil | | | | | | 0 | | | | | |
| No | Uridine | | | | | | 0 | | | | | |
| No | Xanthine | | | | | | 0 | | | | | |
| No | Biotin (B7) | 8 | 21 | 26 | 5 | 0 | 60 | 0.1 | 0.4 | 0.4 | 0.1 | 0.0 |
| No | Coenzyme A | 12 | 0 | 7 | 12 | 0 | 31 | 0.4 | 0.0 | 0.2 | 0.4 | 0.0 |
| No | Dihydropteroate | | | | | | 0 | | | | | |
| No | 1-deoxy-D-xylulose 5-phosphate | | | | | | 0 | | | | | |
| No | Folate (B9) | 12 | 0 | 7 | 12 | 0 | 31 | 0.4 | 0.0 | 0.2 | 0.4 | 0.0 |
| No | Nicotinate | | | | | | 0 | | | | | |
| No | Nicotinamide D-ribonucleotide | 15 | 8 | 25 | 10 | 0 | 58 | 0.3 | 0.1 | 0.4 | 0.2 | 0.0 |
| No | Pyridoxine 5-phosphate (B6) | 0 | 1 | 5 | 0 | 0 | 6 | 0.0 | 0.2 | 0.8 | 0.0 | 0.0 |
| No | Pantothenate (B5) | | | | | | 0 | | | | | |
| No | Pyridoxamine (B6) | 2 | 1 | 0 | 0 | 0 | 3 | 0.7 | 0.3 | 0.0 | 0.0 | 0.0 |
| Yes | Pyridoxal 5'-phosphate (B6) | 30 | 3 | 2 | 4 | 0 | 39 | 0.8 | 0.1 | 0.1 | 0.1 | 0.0 |
| No | Riboflavin (B2) | 10 | 2 | 12 | 9 | 0 | 33 | 0.3 | 0.1 | 0.4 | 0.3 | 0.0 |
| No | Tetrahydrofolate (B9) | 3 | 1 | 7 | 3 | 0 | 14 | 0.2 | 0.1 | 0.5 | 0.2 | 0.0 |
| No | Thiamin (B1) | 12 | 0 | 7 | 12 | 0 | 31 | 0.4 | 0.0 | 0.2 | 0.4 | 0.0 |
| No | Toxopyrimidine | | | | | | 0 | | | | | |
| Yes | Ammonium | 0 | 61 | 0 | 0 | 0 | 61 | 0.0 | 1.0 | 0.0 | 0.0 | 0.0 |
| No | L-Cysteinylglycine | | | | | | 0 | | | | | |
| No | L-methionyl-L-alanine | | | | | | 0 | | | | | |
| No | Hydrogen sulfide | 12 | 0 | 19 | 16 | 6 | 53 | 0.2 | 0.0 | 0.4 | 0.3 | 0.1 |
| Yes | Sulfate | 36 | 18 | 0 | 0 | 0 | 54 | 0.7 | 0.3 | 0.0 | 0.0 | 0.0 |

**Table S4D. Effect of community size and medium type on metabolite richness. Tests with significant p values are shown in bold.**

| | Consumption | | Production | |
|---|---|---|---|---|
| **ANOVA** | Effect test | P-value | Effect test | P-value |
| Community size | $F_{4,81.63} = 0.416$ | 0.7968 | $F_{4,97.96} = 0.227$ | 0.9229 |
| Medium type | $\mathbf{F_{2,196.72} = 42.572}$ | **$4.3 \times 10^{-16}$** | $\mathbf{F_{2,197.55} = 3.397}$ | **0.0354** |
| Interaction | $\mathbf{F_{8,196.72} = 3.803}$ | **0.0004** | $\mathbf{F_{8,197.55} = 2.404}$ | **0.0170** |
| **Analysis of deviance** | Effect test | P-value | Effect test | P-value |
| Microbial taxa | $\boldsymbol{\chi^2_1 = 85.86}$ | **$< 2.2 \times 10^{-16}$** | $\boldsymbol{\chi^2_1 = 7.89}$ | **0.0050** |
| Microbial treatment | $\chi^2_1 = 2.437$ | 0.1185 | $\chi^2_1 = 0.352$ | 0.5529 |
| **$R^2$** | | | | |
| Marginal | 0.5734709 | | 0.1072632 | |
| Conditional | 0.760849 | | 0.207347 | |

**Table S5A. Predicted total number of times metabolite is consumed or produced by individual bacteria in all simulations - rich medium.**

| | *Acetobacter fabarum* | *Acetobacter pomorum* | *Acetobacter tropicalis* | *Lactobacillus brevis* | *Lactobacillus plantarum* |
|---|---|---|---|---|---|
| **Total number of metabolites used in all simulations** | **41** | **40** | **52** | **65** | **64** |
| **Number of times used variably** | **0** | **0** | **1** | **2** | **1** |
| **% metabolite use variability** | **0** | **0** | **2** | **3** | **2** |

| **Metabolite** | **Consume** | **Produce** | **Consume** | **Produce** | **Consume** | **Produce** | **Consume** | **Produce** | **Consume** | **Produce** |
|---|---|---|---|---|---|---|---|---|---|---|
| meso-2,6-Diaminoheptanedioate | 0 | 0 | 0 | 0 | 0 | 0 | 16 | 0 | 0 | 0 |
| D-Alanine | 0 | 16 | 0 | 16 | 0 | 16 | 0 | 0 | 0 | 0 |
| Alanine | 0 | 0 | 0 | 0 | 0 | 0 | 16 | 0 | 12 | 0 |
| Arginine | 16 | 0 | 16 | 0 | 16 | 0 | 16 | 0 | 16 | 0 |
| Asparagine | 16 | 0 | 16 | 0 | 16 | 0 | 16 | 0 | 16 | 0 |
| Aspartate | 3 | 0 | 4 | 0 | 10 | 0 | 16 | 0 | 15 | 0 |
| Cysteine | 0 | 0 | 0 | 0 | 0 | 0 | 16 | 0 | 0 | 0 |
| Glutamine | 16 | 0 | 16 | 0 | 16 | 0 | 16 | 0 | 13 | 0 |
| Glutamate | 0 | 0 | 0 | 0 | 1 | 0 | 10 | 0 | 10 | 1 |
| Glycine | 16 | 0 | 16 | 0 | 16 | 0 | 0 | 0 | 0 | 0 |
| Homocysteine | 0 | 0 | 0 | 0 | 0 | 0 | 0 | 16 | 0 | 0 |
| Histidine | 0 | 0 | 0 | 0 | 0 | 0 | 16 | 0 | 0 | 0 |
| Isoleucine | 0 | 0 | 0 | 0 | 0 | 0 | 16 | 0 | 16 | 0 |
| Leucine | 0 | 0 | 0 | 0 | 0 | 0 | 16 | 0 | 15 | 0 |
| Lysine | 0 | 0 | 0 | 0 | 0 | 0 | 16 | 0 | 16 | 0 |
| Methionine | 0 | 7 | 0 | 16 | 0 | 5 | 8 | 0 | 1 | 0 |
| Ornithine | 0 | 16 | 0 | 16 | 0 | 16 | 0 | 2 | 0 | 1 |
| Phenylalanine | 16 | 0 | 16 | 0 | 16 | 0 | 16 | 0 | 16 | 0 |
| Proline | 16 | 0 | 16 | 0 | 16 | 0 | 16 | 0 | 15 | 0 |
| Serine | 16 | 0 | 16 | 0 | 16 | 0 | 0 | 16 | 16 | 0 |
| Threonine | 0 | 0 | 0 | 0 | 0 | 0 | 16 | 0 | 16 | 0 |
| Tryptophan | 16 | 0 | 16 | 0 | 16 | 0 | 16 | 0 | 16 | 0 |
| Tyrosine | 16 | 0 | 16 | 0 | 16 | 0 | 16 | 0 | 15 | 0 |
| Valine | 0 | 0 | 0 | 0 | 0 | 0 | 16 | 0 | 16 | 0 |
| (2-Aminoethyl)phosphonate | 0 | 0 | 0 | 0 | 0 | 0 | 0 | 0 | 16 | 0 |
| 2-Dehydro-3-deoxy-D-gluconate | 0 | 0 | 0 | 0 | 0 | 0 | 2 | 0 | 0 | 0 |
| L-2-hydroxyisocaproate | 0 | 0 | 0 | 0 | 0 | 0 | 0 | 0 | 1 | 0 |
| (R)-3-(4-Hydroxyphenyl)lactate | 0 | 0 | 0 | 0 | 0 | 0 | 0 | 0 | 1 | 0 |
| 4-Aminobutanoate | 0 | 0 | 0 | 0 | 0 | 0 | 14 | 0 | 12 | 0 |
| Acetate | 11 | 0 | 11 | 0 | 5 | 0 | 0 | 16 | 0 | 16 |
| Acetaldehyde | 8 | 0 | 12 | 0 | 8 | 0 | 4 | 0 | 5 | 0 |
| N-Acetyl-D-glucosamine | 16 | 0 | 0 | 0 | 0 | 0 | 16 | 0 | 14 | 0 |
| R Acetoin | 10 | 0 | 12 | 0 | 16 | 0 | 0 | 6 | 0 | 12 |
| S Acetoin | 6 | 0 | 8 | 0 | 15 | 0 | 0 | 0 | 0 | 0 |
| 2-Oxoglutarate | 0 | 16 | 0 | 16 | 0 | 16 | 3 | 0 | 16 | 0 |
| (R,R)-2,3-Butanediol | 0 | 0 | 0 | 0 | 0 | 0 | 6 | 0 | 12 | 0 |
| (S,S)-2,3-Butanediol | 11 | 0 | 8 | 0 | 15 | 0 | 0 | 0 | 0 | 0 |
| Citrate | 0 | 0 | 0 | 0 | 0 | 0 | 0 | 0 | 3 | 0 |
| Ethanol | 0 | 0 | 0 | 0 | 6 | 0 | 0 | 0 | 7 | 0 |
| Formaldehyde | | | | | | | | | | |
| Formate | 0 | 2 | 0 | 0 | 0 | 10 | 0 | 0 | 0 | 0 |
| D-Fructose | 0 | 0 | 0 | 0 | 0 | 0 | 2 | 0 | 12 | 0 |
| Fumarate | 0 | 0 | 0 | 0 | 1 | 0 | 0 | 0 | 0 | 0 |
| D-Glucose | 2 | 0 | 4 | 0 | 0 | 0 | 6 | 0 | 9 | 0 |
| D-Gluconate | 0 | 0 | 2 | 0 | 0 | 0 | 2 | 0 | 0 | 0 |
| Glycerol | 16 | 0 | 16 | 0 | 4 | 0 | 0 | 3 | 2 | 0 |

| | | | | | | | | | | |
|---|---|---|---|---|---|---|---|---|---|---|
| Glycerol 3-phosphate | 0 | 0 | 0 | 0 | 0 | 0 | 13 | 0 | 15 | 0 |
| Glycolate | | | | | | | | | | |
| Imidazole lactate | 0 | 0 | 0 | 0 | 0 | 0 | 0 | 0 | 16 | 0 |
| D-Lactate | 0 | 0 | 0 | 0 | 0 | 0 | 0 | 4 | 0 | 0 |
| L-Lactate | 0 | 0 | 0 | 0 | 0 | 0 | 0 | 3 | 0 | 0 |
| Malate | 0 | 14 | 0 | 14 | 0 | 6 | 0 | 0 | 2 | 0 |
| Maltose | 0 | 0 | 0 | 0 | 3 | 0 | 16 | 0 | 8 | 0 |
| Maltotetraose | 0 | 0 | 0 | 0 | 16 | 0 | 0 | 0 | 0 | 0 |
| D-Mannose | 0 | 0 | 0 | 0 | 6 | 0 | 3 | 0 | 2 | 0 |
| D-Mannitol | 0 | 0 | 0 | 0 | 0 | 0 | 0 | 0 | 10 | 0 |
| Methylglyoxal | 4 | 0 | 8 | 0 | 9 | 0 | 0 | 0 | 0 | 0 |
| Pyruvate | | | | | | | | | | |
| D-Sorbitol | 0 | 0 | 0 | 0 | 0 | 0 | 2 | 0 | 7 | 0 |
| Succinate | 0 | 4 | 0 | 4 | 0 | 4 | 2 | 14 | 15 | 0 |
| Succinyl-CoA | 16 | 0 | 16 | 0 | 16 | 0 | 0 | 0 | 16 | 0 |
| Sucrose | 8 | 0 | 15 | 0 | 9 | 0 | 0 | 0 | 4 | 0 |
| Trehalose | 0 | 0 | 0 | 0 | 6 | 0 | 10 | 0 | 15 | 0 |
| Adenine | 0 | 0 | 0 | 0 | 0 | 0 | 16 | 0 | 16 | 0 |
| Adenosine | 0 | 0 | 0 | 0 | 16 | 0 | 0 | 0 | 0 | 0 |
| Cytosine | 0 | 0 | 0 | 0 | 0 | 0 | 0 | 0 | 2 | 0 |
| Cytidine | 0 | 0 | 0 | 0 | 16 | 0 | 0 | 0 | 0 | 0 |
| Deoxyadenosine | 0 | 0 | 0 | 0 | 10 | 0 | 0 | 0 | 0 | 0 |
| Deoxycytidine | 0 | 0 | 0 | 0 | 5 | 0 | 16 | 0 | 0 | 0 |
| Deoxyribose | 0 | 0 | 0 | 0 | 0 | 0 | 2 | 0 | 0 | 0 |
| dUMP | 14 | 0 | 14 | 0 | 11 | 3 | 0 | 14 | 0 | 6 |
| Deoxyuridine | 0 | 0 | 0 | 0 | 0 | 11 | 9 | 3 | 16 | 0 |
| Guanine | 0 | 0 | 0 | 0 | 0 | 0 | 16 | 0 | 16 | 0 |
| Hypoxanthine | 12 | 0 | 11 | 0 | 0 | 0 | 0 | 8 | 0 | 5 |
| Inosine | 0 | 0 | 0 | 0 | 16 | 0 | 8 | 0 | 3 | 0 |
| Orotate | | | | | | | | | | |
| Thymidine | 0 | 0 | 0 | 0 | 0 | 0 | 16 | 0 | 16 | 0 |
| Uracil | 16 | 0 | 16 | 0 | 10 | 0 | 0 | 16 | 0 | 16 |
| Uridine | 0 | 0 | 0 | 0 | 0 | 2 | 16 | 0 | 16 | 0 |
| Xanthine | 4 | 0 | 5 | 0 | 0 | 16 | 0 | 0 | 0 | 0 |
| Biotin (B7) | 16 | 0 | 16 | 0 | 16 | 0 | 16 | 0 | 16 | 0 |
| Coenzyme A | 0 | 0 | 0 | 0 | 0 | 0 | 16 | 0 | 0 | 16 |
| Dihydropteroate | | | | | | | | | | |
| 1-deoxy-D-xylulose 5-phosphate | 0 | 0 | 0 | 0 | 0 | 0 | 16 | 0 | 0 | 0 |
| Folate (B9) | 0 | 0 | 0 | 0 | 0 | 0 | 16 | 0 | 16 | 0 |
| Nicotinate | | | | | | | | | | |
| Nicotinamide D-ribonucleotide | 16 | 0 | 16 | 0 | 16 | 0 | 16 | 0 | 16 | 0 |
| Pyridoxine 5-phosphate (B6) | 16 | 0 | 16 | 0 | 16 | 0 | 0 | 0 | 0 | 0 |
| Pantothenate (B5) | | | | | | | | | | |
| Pyridoxamine (B6) | 0 | 0 | 0 | 0 | 0 | 0 | 0 | 9 | 0 | 15 |
| Pyridoxal 5'-phosphate (B6) | 0 | 0 | 0 | 0 | 0 | 0 | 16 | 0 | 16 | 0 |
| Riboflavin (B2) | 16 | 0 | 16 | 0 | 16 | 0 | 0 | 0 | 0 | 0 |
| Tetrahydrofolate (B9) | 16 | 0 | 16 | 0 | 16 | 0 | 0 | 0 | 0 | 0 |
| Thiamin (B1) | 0 | 0 | 0 | 0 | 0 | 0 | 0 | 16 | 16 | 0 |
| Toxopyrimidine | 0 | 0 | 0 | 0 | 0 | 0 | 16 | 0 | 0 | 0 |
| Ammonium | 0 | 16 | 0 | 16 | 0 | 16 | 0 | 16 | 0 | 6 |
| L-Cysteinylglycine | 0 | 0 | 0 | 0 | 0 | 0 | 14 | 0 | 16 | 0 |
| L-methionyl-L-alanine | 16 | 0 | 16 | 0 | 16 | 0 | 15 | 0 | 15 | 0 |
| Hydrogen sulfide | 16 | 0 | 16 | 0 | 16 | 0 | 0 | 0 | 0 | 0 |
| Sulfate | 16 | 0 | 16 | 0 | 16 | 0 | 0 | 0 | 0 | 0 |

**Table S5B. Predicted total number of times metabolite is consumed or produced by individual bacteria in all simulations - base medium.**

| | *Acetobacter fabarum* | *Acetobacter pomorum* | *Acetobacter tropicalis* | *Lactobacillus brevis* | *Lactobacillus plantarum* |
|---|---|---|---|---|---|
| **Total number of metabolites used in all simulations** | **35** | **35** | **35** | **47** | **43** |
| **Number of times used variably** | **1** | **1** | **2** | **1** | **1** |
| **% metabolite use variability** | **3** | **3** | **6** | **2** | **2** |

| | *Acetobacter fabarum* | | *Acetobacter pomorum* | | *Acetobacter tropicalis* | | *Lactobacillus brevis* | | *Lactobacillus plantarum* | |
|---|---|---|---|---|---|---|---|---|---|---|
| **Metabolite** | **Consume** | **Produce** | **Consume** | **Produce** | **Consume** | **Produce** | **Consume** | **Produce** | **Consume** | **Produce** |
| meso-2,6-Diaminoheptanedioate | 0 | 0 | 0 | 0 | 0 | 0 | 16 | 0 | 0 | 0 |
| D-Alanine | 0 | 16 | 0 | 16 | 0 | 16 | 0 | 0 | 0 | 0 |
| Alanine | 0 | 0 | 0 | 0 | 0 | 0 | 16 | 0 | 14 | 0 |
| Arginine | 16 | 0 | 16 | 0 | 16 | 0 | 16 | 0 | 16 | 0 |
| Asparagine | 16 | 0 | 16 | 0 | 16 | 0 | 16 | 0 | 16 | 0 |
| Aspartate | 6 | 0 | 8 | 0 | 12 | 0 | 2 | 0 | 12 | 0 |
| Cysteine | 0 | 0 | 0 | 0 | 0 | 0 | 16 | 0 | 16 | 0 |
| Glutamine | 16 | 0 | 16 | 0 | 16 | 0 | 16 | 0 | 16 | 0 |
| Glutamate | 0 | 0 | 0 | 0 | 0 | 0 | 0 | 5 | 6 | 0 |
| Glycine | 12 | 0 | 12 | 0 | 16 | 0 | 0 | 10 | 0 | 9 |
| Homocysteine | 0 | 0 | 0 | 0 | 0 | 0 | 0 | 16 | 0 | 0 |
| Histidine | 0 | 0 | 0 | 0 | 0 | 0 | 16 | 0 | 0 | 0 |
| Isoleucine | 0 | 0 | 0 | 0 | 0 | 0 | 16 | 0 | 16 | 0 |
| Leucine | 0 | 0 | 0 | 0 | 0 | 0 | 16 | 0 | 16 | 0 |
| Lysine | 0 | 0 | 0 | 0 | 0 | 0 | 16 | 0 | 16 | 0 |
| Methionine | 16 | 0 | 16 | 0 | 16 | 0 | 16 | 0 | 16 | 0 |
| Ornithine | 0 | 16 | 0 | 16 | 0 | 16 | 0 | 6 | 0 | 0 |
| Phenylalanine | 16 | 0 | 16 | 0 | 16 | 0 | 16 | 0 | 16 | 0 |
| Proline | 16 | 0 | 16 | 0 | 16 | 0 | 16 | 0 | 16 | 0 |
| Serine | 16 | 0 | 16 | 0 | 16 | 0 | 6 | 10 | 16 | 0 |
| Threonine | 0 | 0 | 0 | 0 | 0 | 0 | 16 | 0 | 16 | 0 |
| Tryptophan | 16 | 0 | 16 | 0 | 16 | 0 | 16 | 0 | 16 | 0 |
| Tyrosine | 16 | 0 | 16 | 0 | 16 | 0 | 16 | 0 | 16 | 0 |
| Valine | 0 | 0 | 0 | 0 | 0 | 0 | 16 | 0 | 16 | 0 |
| (2-Aminoethyl)phosphonate | | | | | | | | | | |
| 2-Dehydro-3-deoxy-D-gluconate | | | | | | | | | | |
| L-2-hydroxyisocaproate | | | | | | | | | | |
| (R)-3-(4-Hydroxyphenyl)lactate | | | | | | | | | | |
| 4-Aminobutanoate | | | | | | | | | | |
| Acetate | 14 | 0 | 14 | 0 | 12 | 3 | 0 | 10 | 0 | 16 |
| Acetaldehyde | 2 | 0 | 1 | 0 | 6 | 0 | 0 | 5 | 0 | 6 |
| N-Acetyl-D-glucosamine | | | | | | | | | | |
| R Acetoin | 7 | 0 | 9 | 0 | 6 | 2 | 0 | 7 | 0 | 12 |
| S Acetoin | | | | | | | | | | |
| 2-Oxoglutarate | 0 | 16 | 0 | 16 | 0 | 16 | 0 | 8 | 5 | 3 |
| (R,R)-2,3-Butanediol | | | | | | | | | | |
| (S,S)-2,3-Butanediol | | | | | | | | | | |
| Citrate | | | | | | | | | | |
| Ethanol | | | | | | | | | | |
| Formaldehyde | | | | | | | | | | |
| Formate | 0 | 14 | 0 | 13 | 0 | 14 | 0 | 0 | 0 | 9 |
| D-Fructose | | | | | | | | | | |
| Fumarate | | | | | | | | | | |
| D-Glucose | 16 | 0 | 16 | 0 | 16 | 0 | 16 | 0 | 16 | 0 |
| D-Gluconate | | | | | | | | | | |
| Glycerol | 16 | 0 | 16 | 0 | 2 | 0 | 3 | 0 | 10 | 0 |

| | | | | | | | | | | |
|---|---|---|---|---|---|---|---|---|---|---|
| Glycerol 3-phosphate | | | | | | | | | | |
| Glycolate | 0 | 10 | 0 | 12 | 0 | 16 | 0 | 0 | 0 | 8 |
| Imidazole lactate | 0 | 0 | 0 | 0 | 0 | 0 | 0 | 8 | 8 | 0 |
| D-Lactate | | | | | | | | | | |
| L-Lactate | | | | | | | | | | |
| Malate | 0 | 2 | 0 | 1 | 0 | 4 | 0 | 6 | 0 | 0 |
| Maltose | | | | | | | | | | |
| Maltotetraose | | | | | | | | | | |
| D-Mannose | | | | | | | | | | |
| D-Mannitol | | | | | | | | | | |
| Methylglyoxal | | | | | | | | | | |
| Pyruvate | | | | | | | | | | |
| D-Sorbitol | | | | | | | | | | |
| Succinate | 0 | 4 | 0 | 5 | 0 | 4 | 0 | 4 | 10 | 0 |
| Succinyl-CoA | 6 | 1 | 6 | 2 | 0 | 13 | 0 | 0 | 7 | 0 |
| Sucrose | | | | | | | | | | |
| Trehalose | | | | | | | | | | |
| Adenine | 0 | 0 | 0 | 0 | 0 | 0 | 0 | 5 | 5 | 0 |
| Adenosine | | | | | | | | | | |
| Cytosine | | | | | | | | | | |
| Cytidine | | | | | | | | | | |
| Deoxyadenosine | | | | | | | | | | |
| Deoxycytidine | | | | | | | | | | |
| Deoxyribose | | | | | | | | | | |
| dUMP | 8 | 0 | 8 | 0 | 8 | 0 | 0 | 15 | 2 | 0 |
| Deoxyuridine | | | | | | | | | | |
| Guanine | | | | | | | | | | |
| Hypoxanthine | | | | | | | | | | |
| Inosine | | | | | | | | | | |
| Orotate | 8 | 0 | 8 | 0 | 8 | 0 | 0 | 15 | 8 | 0 |
| Thymidine | | | | | | | | | | |
| Uracil | | | | | | | | | | |
| Uridine | | | | | | | | | | |
| Xanthine | | | | | | | | | | |
| Biotin (B7) | 16 | 0 | 16 | 0 | 16 | 0 | 16 | 0 | 16 | 0 |
| Coenzyme A | 0 | 0 | 0 | 0 | 0 | 0 | 0 | 5 | 5 | 0 |
| Dihydropteroate | 0 | 0 | 0 | 0 | 0 | 0 | 16 | 0 | 0 | 0 |
| 1-deoxy-D-xylulose 5-phosphate | 0 | 0 | 0 | 0 | 0 | 0 | 16 | 0 | 0 | 0 |
| Folate (B9) | 0 | 0 | 0 | 0 | 0 | 0 | 0 | 8 | 8 | 0 |
| Nicotinate | 0 | 0 | 0 | 0 | 0 | 0 | 16 | 0 | 16 | 0 |
| Nicotinamide D-ribonucleotide | 12 | 0 | 12 | 0 | 12 | 0 | 0 | 11 | 0 | 10 |
| Pyridoxine 5-phosphate (B6) | 2 | 0 | 2 | 0 | 0 | 3 | 0 | 0 | 0 | 0 |
| Pantothenate (B5) | 10 | 0 | 10 | 0 | 16 | 0 | 16 | 0 | 4 | 0 |
| Pyridoxamine (B6) | 0 | 0 | 0 | 0 | 0 | 0 | 0 | 11 | 0 | 16 |
| Pyridoxal 5'-phosphate (B6) | 0 | 0 | 0 | 0 | 0 | 0 | 16 | 0 | 16 | 0 |
| Riboflavin (B2) | 6 | 0 | 2 | 0 | 0 | 7 | 0 | 0 | 0 | 0 |
| Tetrahydrofolate (B9) | 6 | 0 | 4 | 0 | 0 | 8 | 0 | 0 | 0 | 0 |
| Thiamin (B1) | 0 | 0 | 0 | 0 | 0 | 0 | 0 | 16 | 7 | 0 |
| Toxopyrimidine | 0 | 0 | 0 | 0 | 0 | 0 | 16 | 0 | 0 | 0 |
| Ammonium | 0 | 16 | 0 | 16 | 0 | 16 | 0 | 0 | 0 | 1 |
| L-Cysteinylglycine | | | | | | | | | | |
| L-methionyl-L-alanine | | | | | | | | | | |
| Hydrogen sulfide | 16 | 0 | 16 | 0 | 16 | 0 | 0 | 0 | 0 | 0 |
| Sulfate | 14 | 0 | 14 | 0 | 16 | 0 | 0 | 0 | 0 | 0 |

**Table S5C. Predicted total number of times metabolite is consumed or produced by individual bacteria in all simulations - minimal medium.**

| | *Acetobacter fabarum* | *Acetobacter pomorum* | *Acetobacter tropicalis* | *Lactobacillus brevis* | *Lactobacillus plantarum* |
|---|---|---|---|---|---|
| **Total number of metabolites used in all simulations** | **39** | **38** | **38** | **39** | **41** |
| **Number of times used variably** | **11** | **11** | **7** | **1** | **1** |
| **% metabolite use variability** | **28** | **29** | **18** | **3** | **2** |

| **Metabolite** | **Consume** | **Produce** | **Consume** | **Produce** | **Consume** | **Produce** | **Consume** | **Produce** | **Consume** | **Produce** |
|---|---|---|---|---|---|---|---|---|---|---|
| meso-2,6-Diaminoheptanedioate | 0 | 0 | 0 | 0 | 0 | 0 | 7 | 0 | 0 | 7 |
| D-Alanine | 0 | 11 | 0 | 12 | 0 | 12 | 0 | 0 | 0 | 0 |
| Alanine | | | | | | | | | | |
| Arginine | 4 | 4 | 10 | 1 | 4 | 6 | 0 | 7 | 14 | 0 |
| Asparagine | 11 | 0 | 4 | 1 | 0 | 5 | 7 | 0 | 0 | 9 |
| Aspartate | | | | | | | | | | |
| Cysteine | 0 | 3 | 0 | 2 | 0 | 2 | 7 | 0 | 0 | 0 |
| Glutamine | 0 | 0 | 1 | 0 | 0 | 1 | 0 | 0 | 0 | 0 |
| Glutamate | 0 | 1 | 0 | 2 | 0 | 1 | 7 | 0 | 0 | 4 |
| Glycine | 2 | 0 | 4 | 0 | 5 | 0 | 7 | 0 | 0 | 11 |
| Homocysteine | 0 | 0 | 0 | 0 | 0 | 0 | 0 | 7 | 0 | 0 |
| Histidine | 0 | 2 | 0 | 4 | 0 | 2 | 7 | 0 | 0 | 0 |
| Isoleucine | 0 | 6 | 0 | 7 | 0 | 7 | 7 | 0 | 14 | 0 |
| Leucine | 0 | 4 | 0 | 8 | 0 | 5 | 7 | 0 | 14 | 0 |
| Lysine | 0 | 0 | 0 | 0 | 0 | 0 | 7 | 0 | 0 | 7 |
| Methionine | 0 | 3 | 0 | 1 | 0 | 3 | 7 | 0 | 0 | 0 |
| Ornithine | 4 | 4 | 1 | 10 | 6 | 4 | 7 | 0 | 0 | 14 |
| Phenylalanine | 10 | 0 | 10 | 0 | 8 | 3 | 7 | 0 | 0 | 14 |
| Proline | 4 | 1 | 4 | 0 | 8 | 0 | 7 | 0 | 0 | 11 |
| Serine | 8 | 0 | 8 | 0 | 8 | 0 | 7 | 0 | 0 | 14 |
| Threonine | 0 | 3 | 0 | 1 | 0 | 3 | 7 | 0 | 0 | 0 |
| Tryptophan | 2 | 4 | 3 | 8 | 0 | 7 | 7 | 0 | 12 | 0 |
| Tyrosine | 9 | 0 | 7 | 0 | 8 | 3 | 7 | 0 | 0 | 13 |
| Valine | 0 | 3 | 0 | 8 | 0 | 4 | 7 | 0 | 14 | 0 |
| (2-Aminoethyl)phosphonate | | | | | | | | | | |
| 2-Dehydro-3-deoxy-D-gluconate | | | | | | | | | | |
| L-2-hydroxyisocaproate | | | | | | | | | | |
| (R)-3-(4-Hydroxyphenyl)lactate | 0 | 0 | 0 | 0 | 0 | 0 | 0 | 3 | 3 | 0 |
| 4-Aminobutanoate | | | | | | | | | | |
| Acetate | 10 | 0 | 8 | 0 | 8 | 2 | 1 | 0 | 0 | 14 |
| Acetaldehyde | 3 | 0 | 0 | 0 | 0 | 0 | 0 | 0 | 0 | 3 |
| N-Acetyl-D-glucosamine | | | | | | | | | | |
| R Acetoin | 6 | 0 | 2 | 1 | 0 | 1 | 0 | 1 | 0 | 6 |
| S Acetoin | | | | | | | | | | |
| 2-Oxoglutarate | 2 | 4 | 1 | 8 | 0 | 10 | 0 | 7 | 14 | 0 |
| (R,R)-2,3-Butanediol | | | | | | | | | | |
| (S,S)-2,3-Butanediol | | | | | | | | | | |
| Citrate | | | | | | | | | | |
| Ethanol | | | | | | | | | | |
| Formaldehyde | 0 | 6 | 0 | 5 | 0 | 8 | 0 | 0 | 0 | 0 |
| Formate | 0 | 0 | 0 | 0 | 8 | 0 | 0 | 0 | 0 | 8 |
| D-Fructose | | | | | | | | | | |
| Fumarate | | | | | | | | | | |
| D-Glucose | 11 | 0 | 12 | 0 | 12 | 0 | 7 | 0 | 14 | 0 |
| D-Gluconate | | | | | | | | | | |
| Glycerol | 11 | 0 | 12 | 0 | 7 | 0 | 0 | 0 | 14 | 0 |

| | | | | | | | | | | |
|---|---|---|---|---|---|---|---|---|---|---|
| Glycerol 3-phosphate | | | | | | | | | | |
| Glycolate | 0 | 6 | 0 | 9 | 0 | 12 | 0 | 0 | 0 | 14 |
| Imidazole lactate | 0 | 0 | 0 | 0 | 0 | 0 | 0 | 4 | 4 | 0 |
| D-Lactate | | | | | | | | | | |
| L-Lactate | | | | | | | | | | |
| Malate | 0 | 3 | 0 | 0 | 0 | 2 | 0 | 0 | 3 | 0 |
| Maltose | | | | | | | | | | |
| Maltotetraose | | | | | | | | | | |
| D-Mannose | | | | | | | | | | |
| D-Mannitol | | | | | | | | | | |
| Methylglyoxal | | | | | | | | | | |
| Pyruvate | 0 | 0 | 0 | 0 | 0 | 0 | 6 | 0 | 0 | 6 |
| D-Sorbitol | | | | | | | | | | |
| Succinate | 0 | 4 | 0 | 2 | 0 | 0 | 0 | 1 | 6 | 0 |
| Succinyl-CoA | 6 | 3 | 3 | 4 | 0 | 11 | 0 | 0 | 14 | 0 |
| Sucrose | | | | | | | | | | |
| Trehalose | | | | | | | | | | |
| Adenine | 0 | 0 | 0 | 0 | 0 | 0 | 7 | 0 | 0 | 7 |
| Adenosine | | | | | | | | | | |
| Cytosine | | | | | | | | | | |
| Cytidine | | | | | | | | | | |
| Deoxyadenosine | | | | | | | | | | |
| Deoxycytidine | | | | | | | | | | |
| Deoxyribose | | | | | | | | | | |
| dUMP | 5 | 2 | 7 | 0 | 3 | 6 | 0 | 7 | 11 | 0 |
| Deoxyuridine | | | | | | | | | | |
| Guanine | | | | | | | | | | |
| Hypoxanthine | | | | | | | | | | |
| Inosine | | | | | | | | | | |
| Orotate | 4 | 0 | 3 | 0 | 0 | 0 | 0 | 1 | 1 | 5 |
| Thymidine | | | | | | | | | | |
| Uracil | | | | | | | | | | |
| Uridine | | | | | | | | | | |
| Xanthine | | | | | | | | | | |
| Biotin (B7) | 5 | 4 | 7 | 2 | 4 | 8 | 7 | 0 | 14 | 0 |
| Coenzyme A | 0 | 0 | 0 | 0 | 0 | 0 | 7 | 0 | 0 | 7 |
| Dihydropteroate | | | | | | | | | | |
| 1-deoxy-D-xylulose 5-phosphate | | | | | | | | | | |
| Folate (B9) | 0 | 0 | 0 | 0 | 0 | 0 | 7 | 0 | 0 | 7 |
| Nicotinate | | | | | | | | | | |
| Nicotinamide D-ribonucleotide | 2 | 3 | 2 | 3 | 0 | 11 | 7 | 0 | 14 | 0 |
| Pyridoxine 5-phosphate (B6) | 3 | 0 | 2 | 1 | 0 | 3 | 0 | 0 | 0 | 0 |
| Pantothenate (B5) | | | | | | | | | | |
| Pyridoxamine (B6) | 0 | 0 | 0 | 0 | 0 | 0 | 1 | 0 | 1 | 0 |
| Pyridoxal 5'-phosphate (B6) | 0 | 0 | 0 | 0 | 0 | 0 | 3 | 2 | 13 | 0 |
| Riboflavin (B2) | 7 | 1 | 5 | 1 | 0 | 8 | 0 | 0 | 0 | 0 |
| Tetrahydrofolate (B9) | 4 | 1 | 3 | 0 | 0 | 5 | 0 | 0 | 0 | 0 |
| Thiamin (B1) | 0 | 0 | 0 | 0 | 0 | 0 | 7 | 0 | 0 | 7 |
| Toxopyrimidine | | | | | | | | | | |
| Ammonium | 11 | 0 | 12 | 0 | 12 | 0 | 7 | 0 | 14 | 0 |
| L-Cysteinylglycine | | | | | | | | | | |
| L-methionyl-L-alanine | | | | | | | | | | |
| Hydrogen sulfide | 0 | 8 | 0 | 3 | 0 | 8 | 0 | 0 | 14 | 0 |
| Sulfate | 11 | 0 | 12 | 0 | 12 | 0 | 0 | 0 | 0 | 0 |

**Table S5D. Effect of taxa, community size, and medium type on metabolite consumption and release rates. Tests with significant p values are shown in bold.**

| **Multivariate correlation** | Consumption | | Production | |
|---|---|---|---|---|
| | Effect size ($R^2$) | P-value | Effect size ($R^2$) | P-value |
| AF | 0.033 | 0.020 | **0.061** | **0.002** |
| AP | 0.026 | 0.030 | 0.027 | 0.046 |
| AT | **0.155** | **0.001** | **0.047** | **0.006** |
| LB | **0.263** | **0.001** | **0.097** | **0.001** |
| LP | 0.018 | 0.117 | **0.066** | **0.001** |
| **PERMANOVA** | Effect test | Effect size ($R^2$) | Effect test | Effect size ($R^2$) |
| Taxon | **$F_{4,221} = 13.96$, $p = 0.001$** | **0.119** | **$F_{4,221} = 10.78$, $p = 0.001$** | **0.115** |
| Community size | $F_{4,221} = 0.918$, $p = 0.508$ | 0.008 | $F_{4,221} = 2.04$, $p = 0.037$ | 0.022 |
| Medium type | **$F_{2,221} = 90.67$, $p = 0.001$** | **0.387** | **$F_{2,221} = 43.56$, $p = 0.001$** | **0.232** |
| Community size * medium type | $F_{8,221} = 0.82$, $p = 0.681$ | 0.014 | **$F_{8,221} = 2.10$, $p = 0.016$** | **0.045** |

Table S6. Metabolites predicted to be available to the host.

| **Metabolite group** | **Metabolite** | **Nutrient replete** | **Base** | **Nutrient depleted** |
|---|---|---|---|---|
| **Amino acid** | D-Alanine | ■ | ■ | ■ |
| | Homocysteine | ■ | ■ | ■ |
| | Methionine | ■ | □ | □ |
| | Ornithine | ■ | ■ | □ |
| | Serine | ■ | □ | □ |
| **Central carbon** | Acetate | ■ | ■ | ■ |
| | 2-Oxoglutarate | ■ | ■ | □ |
| | Formaldehyde | □ | □ | ■ |
| | Formate | ■ | ■ | □ |
| | Glycolate | □ | ■ | ■ |
| | D-Lactate | ■ | □ | □ |
| | L-Lactate | ■ | □ | □ |
| | Malate | ■ | ■ | □ |
| | Succinate | ■ | ■ | □ |
| **Nitrogen** | Ammonium | ■ | ■ | □ |
| **Nucleotide** | Hypoxanthine | ■ | □ | □ |
| | Uracil | ■ | □ | □ |
| | Xanthine | ■ | □ | □ |
| **Vitamin & cofactor** | CoA | ■ | □ | □ |
| | Pyridoxamine | ■ | ■ | □ |
| | Thiamin | ■ | ■ | □ |
| **Total number of metabolites** | | 19 | 12 | 5 |

Black rectangles indicate metabolite availability to host, grey rectangles indicate absence

| Table S7A. List of components - rich medium. | | |
|---|---|---|
| **Metabolite name** | **Exchange bound** | **Metabolite group** |
| L-meso-2,6-Diaminoheptanedioate | -0.5 | Amino acid |
| D-Alanine | -0.05 | Amino acid |
| L-Alanine | -0.5 | Amino acid |
| L-Arginine | -0.5 | Amino acid |
| L-Asparagine | -0.5 | Amino acid |
| L-Aspartate | -0.5 | Amino acid |
| Choline | -0.05 | Amino acid |
| Chorismate | -0.05 | Amino acid |
| L-Cysteine | -0.5 | Amino acid |
| L-Glutamine | -0.5 | Amino acid |
| L-Glutamate | -0.5 | Amino acid |
| Glycine | -0.5 | Amino acid |
| Glycine betaine | -0.05 | Amino acid |
| L-Homocysteine | -0.05 | Amino acid |
| L-Histidine | -0.5 | Amino acid |
| Histamine | -0.05 | Amino acid |
| L-Isoleucine | -0.5 | Amino acid |
| L-Leucine | -0.5 | Amino acid |
| L-Lysine | -0.5 | Amino acid |
| D-Methionine | -0.05 | Amino acid |
| L-Methionine | -0.5 | Amino acid |
| L-Methionine Sulfoxide | -0.05 | Amino acid |
| Ornithine | -0.05 | Amino acid |
| L-Phenylalanine | -0.5 | Amino acid |
| L-Proline | -0.5 | Amino acid |
| D-Serine | -0.5 | Amino acid |
| L-Serine | -0.5 | Amino acid |
| L-Threonine | -0.5 | Amino acid |
| L-Tryptophan | -0.5 | Amino acid |
| L-Tyrosine | -0.5 | Amino acid |
| L-Valine | -0.5 | Amino acid |
| Propane-1,2-diol | -0.05 | Carbon |
| (2-Aminoethyl)phosphonate | -0.05 | Carbon |
| 2-Dehydro-3-deoxy-D-gluconate | -0.05 | Carbon |
| L-2-hydroxyisocaproate | -0.05 | Carbon |
| 2-methyl butanoic acid | -0.05 | Carbon |
| 2-Methylbutanal | -0.05 | Carbon |
| 2-methylbutanol | -0.05 | Carbon |
| 2-methylpropanoic acid | -0.05 | Carbon |
| 2-methylpropanal | -0.05 | Carbon |
| 2-methylpropanol | -0.05 | Carbon |
| (R)-3-(4-Hydroxyphenyl)lactate | -0.05 | Carbon |
| 3-methylbutanoic acid | -0.05 | Carbon |
| 3-methylbutanal | -0.05 | Carbon |
| 3-methylbutanol | -0.05 | Carbon |
| 4-Aminobutanoate | -0.05 | Carbon |
| 4-Aminobenzoate | -0.05 | Carbon |
| Acetoacetyl-CoA | -0.05 | Carbon |
| Acetate | -0.05 | Carbon |
| Acetaldehyde | -0.05 | Carbon |
| Acetol | -0.05 | Carbon |
| N-Acetyl-D-galactosamine | -0.05 | Carbon |
| N-Acetyl-D-glucosamine | -0.05 | Carbon |
| (R)-Acetoin | -0.05 | Carbon |
| (S)-Acetoin | -0.05 | Carbon |

| | | |
|---|---|---|
| 2-Oxoglutarate | -0.05 | Carbon |
| L-Arabinose | -0.05 | Carbon |
| (R,R)-2,3-Butanediol | -0.05 | Carbon |
| (S,S)-2,3-Butanediol | -0.05 | Carbon |
| Benzaldehyde | -0.05 | Carbon |
| cellobiose | -0.05 | Carbon |
| Citrate | -0.05 | Carbon |
| Dihydroxyacetone | -0.05 | Carbon |
| Diacetyl | -0.05 | Carbon |
| Ethanolamine | -0.05 | Carbon |
| ethanol | -0.05 | Carbon |
| formaldehyde | -0.05 | Carbon |
| formate | -0.05 | Carbon |
| D-Fructose | -0.05 | Carbon |
| L-Fucose | -0.05 | Carbon |
| Fumarate | -0.05 | Carbon |
| D-Galactose | -0.05 | Carbon |
| Galactitol | -0.05 | Carbon |
| D-Glucose | -1 | Carbon |
| D-Gluconate | -0.05 | Carbon |
| Glycerol | -0.5 | Carbon |
| Glycerol 3-phosphate | -0.05 | Carbon |
| Glycolate | -0.05 | Carbon |
| imidazole lactate | -0.05 | Carbon |
| Indolelactate | -0.05 | Carbon |
| D-Lactate | -0.05 | Carbon |
| L-Lactate | -0.05 | Carbon |
| Lactose | -0.05 | Carbon |
| D-Malate | -0.05 | Carbon |
| L-Malate | -0.05 | Carbon |
| maltose | -0.05 | Carbon |
| maltohexaose | -0.05 | Carbon |
| maltopentaose | -0.05 | Carbon |
| Maltotriose | -0.05 | Carbon |
| maltotetraose | -0.05 | Carbon |
| D-Mannose | -0.05 | Carbon |
| Melibiose | -0.05 | Carbon |
| Methional | -0.05 | Carbon |
| D-Mannitol | -0.05 | Carbon |
| Methylglyoxal | -0.05 | Carbon |
| Orotate | -0.05 | Carbon |
| Phenylacetaldehyde | -0.05 | Carbon |
| Phenylethyl alcohol | -0.05 | Carbon |
| Phenol | -0.05 | Carbon |
| Phenyl lactate | -0.05 | Carbon |
| Pyruvate | -0.05 | Carbon |
| D-Ribose | -0.05 | Carbon |
| D-Sorbitol | -0.05 | Carbon |
| Succinate | -0.05 | Carbon |
| Succinyl-CoA | -0.05 | Carbon |
| sucrose | -0.05 | Carbon |
| trehalose | -0.05 | Carbon |
| D-Xylose | -0.05 | Carbon |
| arsenite | -0.05 | Inorganic ion |
| Calcium | -0.1 | Inorganic ion |
| Cobinamide | -0.05 | Inorganic ion |
| Cob(I)alamin | -0.05 | Inorganic ion |

| | | |
|---|---|---|
| Cadmium | -0.05 | Inorganic ion |
| Chloride | -0.1 | Inorganic ion |
| CO2 | -0.05 | Inorganic ion |
| Co2+ | -0.1 | Inorganic ion |
| Cu+ | -0.05 | Inorganic ion |
| Cu2+ | -0.1 | Inorganic ion |
| Fe2+ | -0.1 | Inorganic ion |
| Fe3+ | -0.1 | Inorganic ion |
| H+ | -0.1 | Inorganic ion |
| H2O | -1 | Inorganic ion |
| Hg2+ | -0.05 | Inorganic ion |
| potassium | -0.1 | Inorganic ion |
| Magnesium | -0.1 | Inorganic ion |
| Mn2+ | -0.1 | Inorganic ion |
| Molybdate | -0.1 | Inorganic ion |
| Sodium | -0.1 | Inorganic ion |
| nickel | -0.1 | Inorganic ion |
| o2 | -2 | Inorganic ion |
| Lead | -0.05 | Inorganic ion |
| Phosphate | -0.1 | Inorganic ion |
| Zinc | -0.1 | Inorganic ion |
| Ammonium | -0.5 | Nitrogen |
| Nitrite | -0.05 | Nitrogen |
| Nitrate | -0.05 | Nitrogen |
| Urea | -0.05 | Nitrogen |
| 5-Methylthio-D-ribose | -0.05 | Nucleotide |
| Adenine | -0.05 | Nucleotide |
| adenosine | -0.05 | Nucleotide |
| Allantoin | -0.05 | Nucleotide |
| Cytosine | -0.05 | Nucleotide |
| cytidine | -0.05 | Nucleotide |
| deoxyadenosine | -0.05 | Nucleotide |
| deoxycytidine | -0.05 | Nucleotide |
| Deoxyribose | -0.05 | Nucleotide |
| deoxyuridine | -0.05 | Nucleotide |
| Guanine | -0.05 | Nucleotide |
| Hypoxanthine | -0.05 | Nucleotide |
| inosine | -0.05 | Nucleotide |
| thymidine | -0.05 | Nucleotide |
| Uracil | -0.05 | Nucleotide |
| uridine | -0.05 | Nucleotide |
| Xanthine | -0.05 | Nucleotide |
| L-alanyl-L-aspartate | -0.05 | Peptide |
| L-alanyl-L-glutamine | -0.05 | Peptide |
| L-alanyl-L-glutamate | -0.05 | Peptide |
| L-alanylglycine | -0.05 | Peptide |
| L-alanyl-L-histidine | -0.05 | Peptide |
| L-alanyl-L-leucine | -0.05 | Peptide |
| L-alanyl-L-threonine | -0.05 | Peptide |
| Cys-Gly | -0.05 | Peptide |
| Glycyl-L-asparagine | -0.05 | Peptide |
| Glycyl-L-aspartate | -0.05 | Peptide |
| Gly-Cys | -0.05 | Peptide |
| Glycyl-L-glutamine | -0.05 | Peptide |
| Glycyl-L-glutamate | -0.05 | Peptide |
| Glycylleucine | -0.05 | Peptide |
| Glycyl-L-methionine | -0.05 | Peptide |

| | | |
|---|---|---|
| Glycylphenylalanine | -0.05 | Peptide |
| Glycylproline | -0.05 | Peptide |
| Glycyl-L-tyrosine | -0.05 | Peptide |
| L-methionyl-L-alanine | -0.05 | Peptide |
| butanesulfonate | -0.05 | Sulfur |
| ethanesulfonate | -0.05 | Sulfur |
| Hydrogen sulfide | -0.5 | Sulfur |
| Hexanesulfonate | -0.05 | Sulfur |
| Isethionic acid | -0.05 | Sulfur |
| L-Cysteate | -0.05 | Sulfur |
| methanesulfonate | -0.05 | Sulfur |
| Putrescine | -0.5 | Sulfur |
| Sulfate | -0.5 | Sulfur |
| Spermidine | -0.5 | Sulfur |
| sulfoacetate | -0.05 | Sulfur |
| Taurine | -0.05 | Sulfur |
| Thiosulfate | -0.05 | Sulfur |
| 4-Amino-5-hydroxymethyl-2-methylpyrimidine | -0.1 | Vitamin |
| 5-Methyltetrahydrofolate | -0.1 | Vitamin |
| Adenosylcobalamin | -0.05 | Vitamin |
| Biotin | -0.1 | Vitamin |
| CoA | -0.05 | Vitamin |
| Dihydropteroate | -0.1 | Vitamin |
| 1-deoxy-D-xylulose 5-phosphate | -0.1 | Vitamin |
| Folate | -0.05 | Vitamin |
| Nicotinate | -0.1 | Vitamin |
| Nicotinamide D-ribonucleotide | -0.05 | Vitamin |
| Pyridoxine 5-phosphate | -0.05 | Vitamin |
| Pantothenate | -0.1 | Vitamin |
| Pyridoxamine | -0.1 | Vitamin |
| Pyridoxal 5'-phosphate | -0.1 | Vitamin |
| Pyridoxine | -0.1 | Vitamin |
| riboflavin | -0.05 | Vitamin |
| 5,6,7,8-Tetrahydrofolate | -0.05 | Vitamin |
| Thiamin | -0.05 | Vitamin |

**Table S7B. List of components - base medium.**

| Metabolite name | Exchange bound | Metabolite group |
|---|---|---|
| L-meso-2,6-Diaminoheptanedioate | -0.05 | Amino acid |
| L-Alanine | -0.05 | Amino acid |
| L-Arginine | -0.05 | Amino acid |
| L-Asparagine | -0.05 | Amino acid |
| L-Aspartate | -0.05 | Amino acid |
| L-Cysteine | -0.05 | Amino acid |
| L-Glutamine | -0.05 | Amino acid |
| L-Glutamate | -0.05 | Amino acid |
| Glycine | -0.05 | Amino acid |
| L-Histidine | -0.05 | Amino acid |
| L-Isoleucine | -0.05 | Amino acid |
| L-Leucine | -0.05 | Amino acid |
| L-Lysine | -0.05 | Amino acid |
| L-Methionine | -0.05 | Amino acid |
| L-Phenylalanine | -0.05 | Amino acid |
| L-Proline | -0.05 | Amino acid |
| D-Serine | -0.05 | Amino acid |
| L-Serine | -0.05 | Amino acid |

| | | |
|---|---|---|
| L-Threonine | -0.05 | Amino acid |
| L-Tryptophan | -0.05 | Amino acid |
| L-Tyrosine | -0.05 | Amino acid |
| L-Valine | -0.05 | Amino acid |
| D-Glucose | -0.1 | Carbon |
| Glycerol | -0.05 | Carbon |
| Calcium | -0.01 | Inorganic ion |
| Chloride | -0.01 | Inorganic ion |
| Co2+ | -0.01 | Inorganic ion |
| Cu2+ | -0.01 | Inorganic ion |
| Fe2+ | -0.01 | Inorganic ion |
| Fe3+ | -0.01 | Inorganic ion |
| H+ | -0.01 | Inorganic ion |
| H2O | -0.1 | Inorganic ion |
| potassium | -0.01 | Inorganic ion |
| Magnesium | -0.01 | Inorganic ion |
| Mn2+ | -0.01 | Inorganic ion |
| Molybdate | -0.01 | Inorganic ion |
| Sodium | -0.01 | Inorganic ion |
| nickel | -0.01 | Inorganic ion |
| o2 | -2 | Inorganic ion |
| Phosphate | -0.01 | Inorganic ion |
| Zinc | -0.01 | Inorganic ion |
| Ammonium | -0.05 | Nitrogen |
| Putrescine | -0.05 | Sulfur |
| Spermidine | -0.05 | Sulfur |
| Hydrogen sulfide | -0.05 | Sulfur |
| Sulfate | -0.05 | Sulfur |
| 4-Amino-5-hydroxymethyl-2-methylpyrimidine | -0.01 | Vitamin |
| 5-Methyltetrahydrofolate | -0.01 | Vitamin |
| Biotin | -0.01 | Vitamin |
| Dihydropteroate | -0.01 | Vitamin |
| 1-deoxy-D-xylulose 5-phosphate | -0.01 | Vitamin |
| Nicotinate | -0.01 | Vitamin |
| Pantothenate | -0.01 | Vitamin |
| Pyridoxamine | -0.01 | Vitamin |
| Pyridoxal 5'-phosphate | -0.01 | Vitamin |
| Pyridoxine | -0.01 | Vitamin |

**Table S7C. List of components - minimal medium.**

| Metabolite name | Exchange bound | Metabolite group |
|---|---|---|
| D-Glucose | -0.1 | Carbon |
| Glycerol | -0.05 | Carbon |
| Calcium | -0.01 | Inorganic ion |
| Chloride | -0.01 | Inorganic ion |
| Co2+ | -0.01 | Inorganic ion |
| Cu2+ | -0.01 | Inorganic ion |
| Fe2+ | -0.01 | Inorganic ion |
| Fe3+ | -0.01 | Inorganic ion |
| H+ | -0.01 | Inorganic ion |
| H2O | -0.1 | Inorganic ion |
| potassium | -0.01 | Inorganic ion |
| Magnesium | -0.01 | Inorganic ion |
| Mn2+ | -0.01 | Inorganic ion |
| Molybdate | -0.01 | Inorganic ion |
| Sodium | -0.01 | Inorganic ion |

| | | |
|---|---|---|
| nickel | -0.01 | Inorganic ion |
| o2 | -2 | Inorganic ion |
| Phosphate | -0.01 | Inorganic ion |
| Zinc | -0.01 | Inorganic ion |
| Ammonium | -0.05 | Nitrogen |
| Sulfate | -0.05 | Sulfur |
| Pyridoxal 5'-phosphate | -0.01 | Vitamin |
